# A machine learning based approach to the segmentation of micro CT data in archaeological and evolutionary sciences

**DOI:** 10.1101/859983

**Authors:** Thomas O’Mahoney, Lidija Mcknight, Tristan Lowe, Maria Mednikova, Jacob Dunn

## Abstract

Segmentation of high-resolution tomographic data is often an extremely time-consuming task and until recently, has usually relied upon researchers manually selecting materials of interest slice by slice. With the exponential rise in datasets being acquired, this is clearly not a sustainable workflow. In this paper, we apply the Trainable Weka Segmentation (a freely available plugin for the multiplatform program ImageJ) to typical datasets found in archaeological and evolutionary sciences. We demonstrate that Trainable Weka Segmentation can provide a fast and robust method for segmentation and is as effective as other leading-edge machine learning segmentation techniques.

## Introduction

Three-dimensional imaging using micro CT scanning has rapidly become mainstream in the archaeological and evolutionary sciences. It enables the high-resolution and non-destructive analysis of internal structures of scientific interest. In archaeological sciences it has been used for a variety of purposes, from imaging pottery (Barron et al., 2017; Tuniz and Zanini, 2018) understanding soil compaction (McBride and Mercer, 2012) imaging early bone tools (Bello et al., 2013) and mummies, both human and animal (Charlier et al., 2014; Du Plessis et al., 2015; Romell et al., 2018). In evolutionary sciences, it is employed even more widely, from scanning hominin remains for morphological reconstruction (Gunz et al., 2012; Hershkovitz et al., 2018, 2015; Hublin et al., 2017) to diagnosing ancient pathologies (Anné et al., 2015; Odes et al., 2016; Randolph-Quinney et al., 2016). It is used extensively in vertebrate paleontology (Abel et al., 2012; Chapelle et al., 2019; Hechenleitner et al., 2016; Laloy et al., 2013) and invertebrate paleontology (Garwood and Dunlop, 2014; Wacey et al., 2017) and increasingly, paleopalynology (Collinson et al., 2016).

In comparative anatomy, Micro-CT scanning is now part of the standard non-destructive analytical toolkit, creating data for use in analyses such as geometric morphometrics (GMM) and finite element analysis (FEA) (Borgard et al., 2019; Brassey et al., 2018, 2013; Brocklehurst et al., 2019; Cuff et al., 2015; Marshall et al., 2019; Polly et al., 2016). Unfortunately for researchers, if one wishes to quantify biological structures, the data does not simply appear from scanners ready to use. It requires processing through the segmentation of the structures of interest, followed (usually) by the generation of 3-dimensional models.

There are 5 main approaches to image segmentation.

- Global thresholding based upon greyscale values in scans (greyscale thresholding).
- Watershed based segmentation
- Locally adaptive segmentation/edge-based segmentation
- Manual segmentation of structures
- Label based segmentation, in conjunction with machine learning.

One can broadly classify greyscale segmentation and edge-based segmentation as passive approaches, as very little input is required from the user, and Region or label-based segmentation as active, in that they require more explicit input from the user. In the following discussion, we use screenshots from Avizo (Thermo Fisher Inc. Dallas, TX) to illustrate our examples. All the algorithms (with the exceptions of local c-means and Weka segmentation) are available within popular commercial software packages as well as ImageJ, through their respective segmentation toolboxes. (For a complementary review of segmentation methods, which goes into more detail in places, please see Noruzi et al., (2014)).

### Greyscale thresholding

Greyscale thresholding is the oldest approach to the processing of tomographic data (Spoor et al., 1993) and has been refined to the use of the half width, full maximum height approach based upon stack histograms (Spoor et al., 1993). This however is only useful for materials which have a single range of X-ray absorption and several passes are therefore required for the segmentation of multiple tissue types. This approach is available in freeware (e.g. ImageJ, MIA (Wollny et al., 2013), Stradview (Treece et al., 2000),3D slicer (Fedorov et al., 2012)) and commercial software (e.g. Avizo, Matlab, DragonFly, Mimics). An example is shown in figure 1.

**Figure 1.**
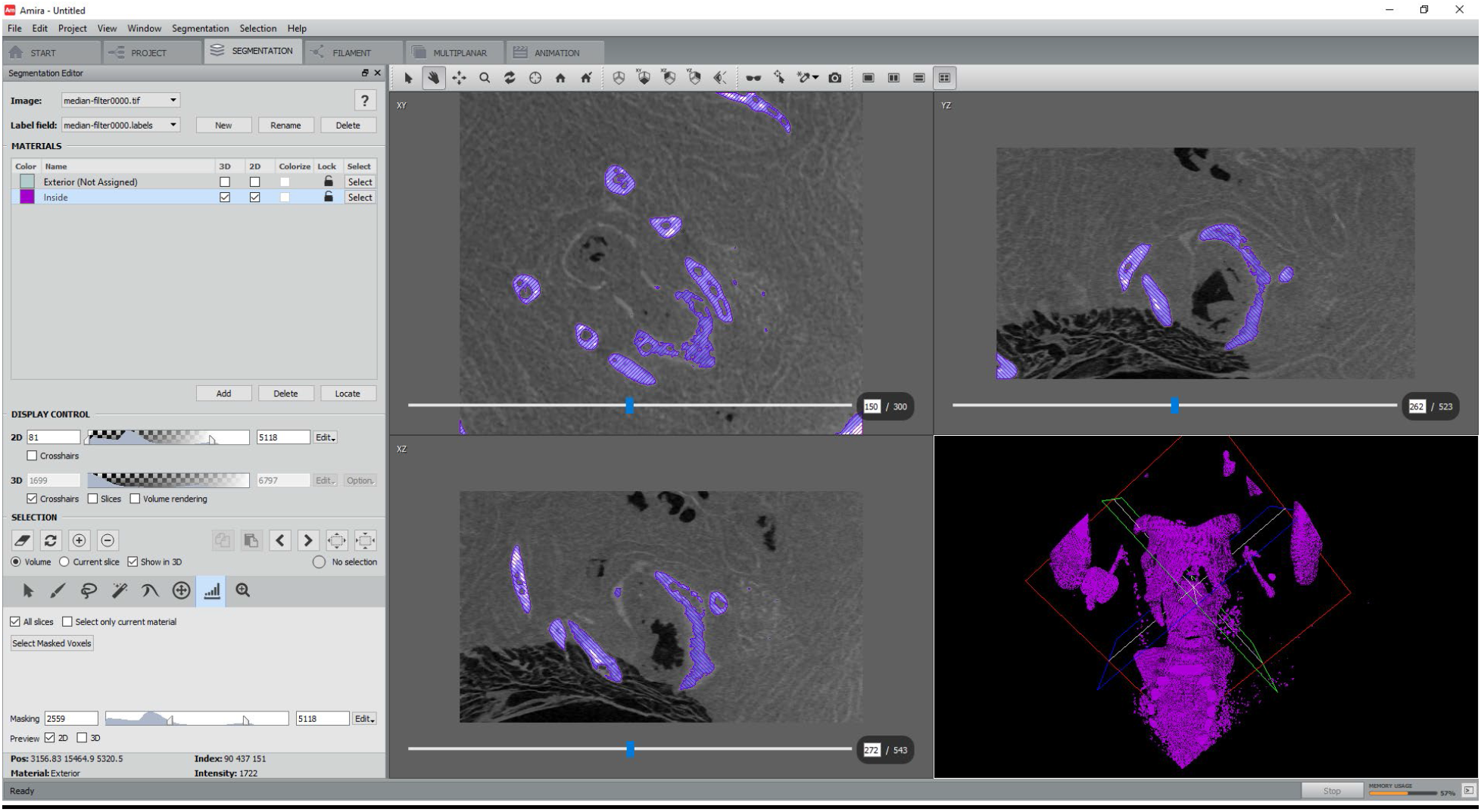
Example of half maximum height grey scale thresholding. Specimen shown is a Nycticebus pygmaeus vocal tract DLC 2901 available from http://www.morphosource.org

### Watershed based segmentation

Watershed based segmentation has enjoyed a lot of popularity for segmenting complex structures such as brain folds but is also of some utility when segmenting fossil structures. A recent innovation has been the application of ray-casting and similar techniques to the processing of data (Dunmore et al., 2018; Scherf and Tilgner, 2009) which helps to ameliorate problems with fuzzy data and automates the processing of this. A problem is that it is only feasible to process a single material (although the others are also detected) and an aspect of ‘re-looping’ the procedure is then required, which can create a bottleneck for scans where multiple materials are of equal interest (for example, mummies, where the skeleton, desiccated flesh and wrappings are all of equal scientific interest. This approach is available in freeware (e.g. ImageJ, 3D Slicer) and commercial software (e.g. Avizo, Matlab, DragonFly, Mimics - see figure 2 for an example of this).

**Figure 2.**
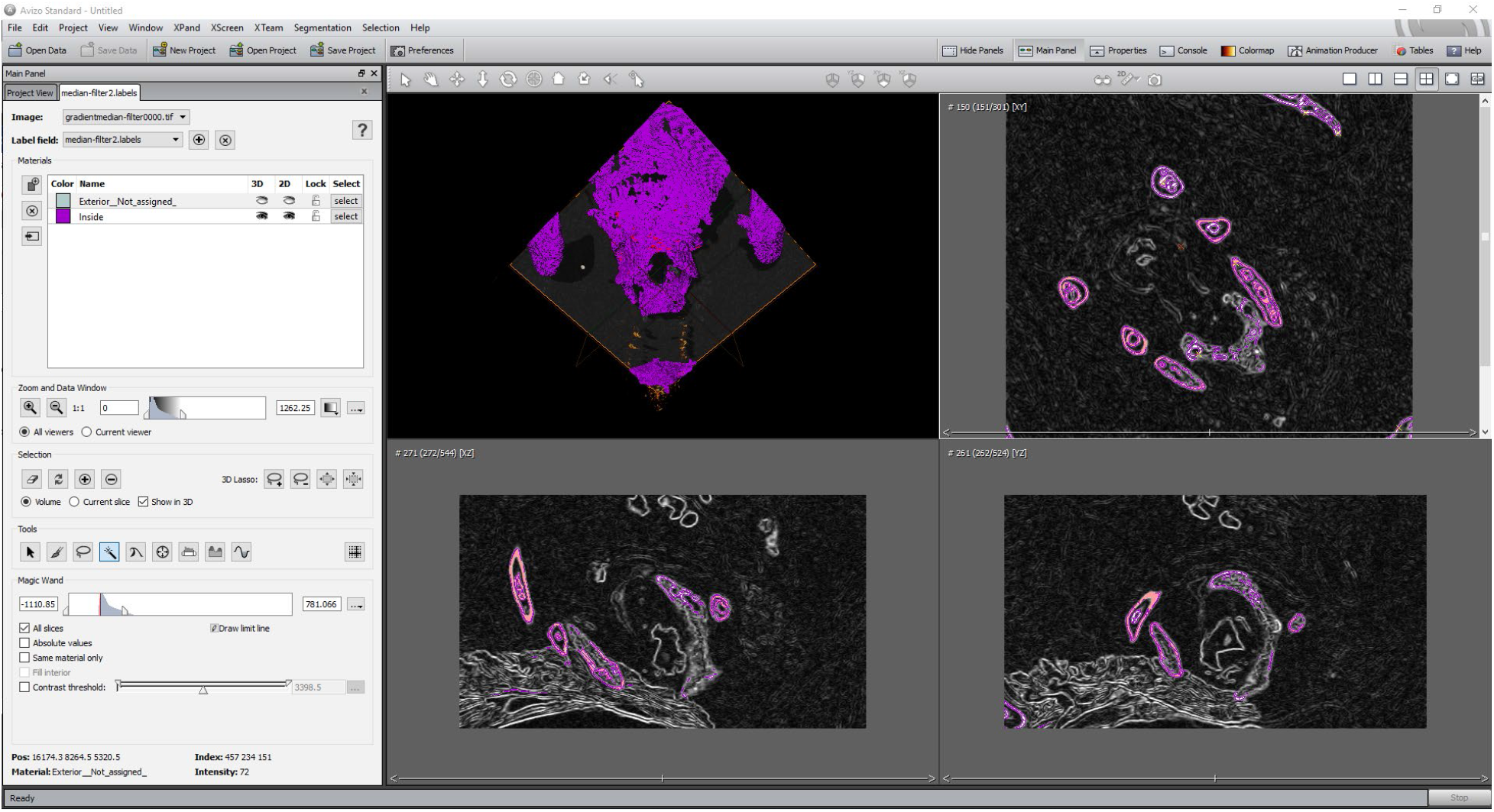
Watershed segmentation. Specimen shown is a Nycticebus pygmaeus vocal tract DLC 2901 available from http://www.morphosource.org

### Locally adaptive segmentation

Locally adaptive segmentation is increasingly carried out using deep learning in an automated fashion. Algorithms use combinations of edge detection, texture similarity and image contrast to create rules for the classification of different materials. (Prasoon et al., 2013; Radford et al., 2015; Suzani et al., 2015). It has become increasingly popular with the availability of massive datasets from healthcare providers and several recent reviews cover this suite of techniques in-depth (Greenspan et al., 2016; Litjens et al., 2017; Shen et al., 2017; Suzuki, 2017). A criticism of unsupervised methods such as convoluted neural networks is that they can demand huge computing resources while still often yielding false positives including highlighting artefacts in data (e.g. ring artefacts in CT scans (Nguyen et al., 2015; Szegedy et al., 2013; Wang, 2016). Another set of related techniques include kmeans and c-means clustering algorithms. K-means is known as a ‘hard’ clustering algorithm, introduced independently by Forgy and MacQueen (Forgy, 1965; MacQueen, 1967). This method, and extensions of it, have been used widely in MRI processing (e.g. (Dimitriadou et al., 2004; Juang and Wu, 2010; Singh et al., 1996). An interesting note is that both Forgy and MacQueen (Forgy, 1965; MacQueen, 1967) cautioned against using k-means clustering as a definitive algorithm, but as an aid to the user in interpreting clusters of data. As such, one whould always apply a ‘sense check’ to data resulting from this clustering method.

Another popular clustering algorithm is that of fuzzy c-means (Bezdek, 1980, 1980, 1975; Pham and Prince, 1999) which is an example of ‘soft’ clustering methods, where probabilities of group allocation are given. Again, this is popular for the automated segmentation of MRI data (e.g. Dimitriadou et al., 2004) and computational speed can be further improved by an initial clustering using k-means partitioning (Stetco et al., 2015).

Both k-means and c-means clustering can be applied in two different fashions-either slice by slice (two-dimensional clustering) or to the volume as a whole, as a three-dimensional matrix (three-dimensional clustering). Recent work by Dunmore et al., (2018) further refined this process by using a multi-step methodology. This comprised of initial clustering of data through k-means (which is a very common method of initializing local c-means clustering). This is followed by a global c-means clustering, with a final step of localized c-means clustering of overlapping sub volumes. Materials are assigned based on these summed probabilities, following a user-defined probability threshold (Dunmore et al., 2018). An example of results obtained through 3D kmeans is shown in figure 3. Different implementations of clustering are available in freeware (e.g. ImageJ, 3D Slicer, MIA (Wollny et al., 2013)) and commercial software (e.g. Avizo, Matlab, DragonFly, Mimics).

**Figure 3.**
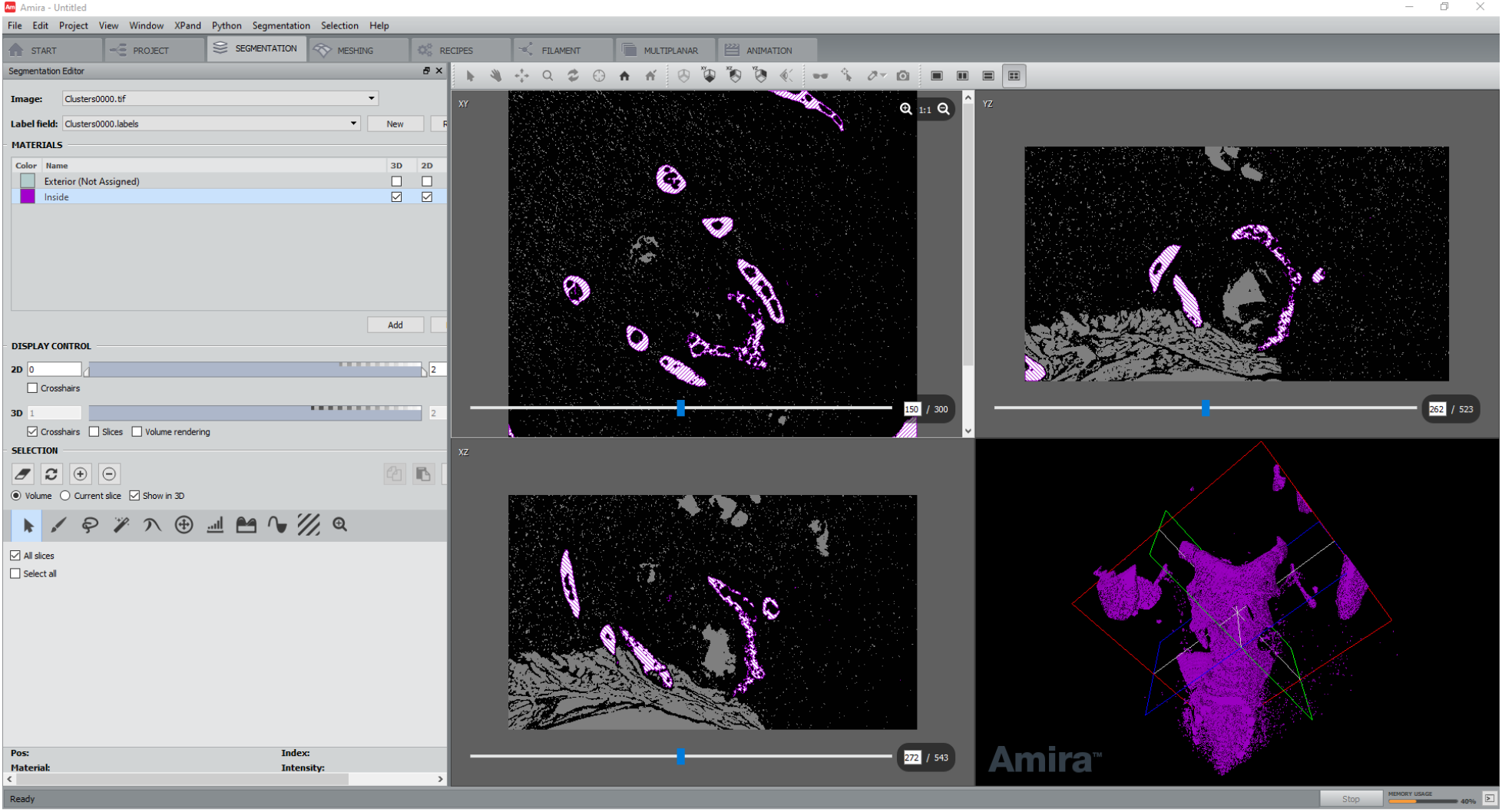
3D k-means segmentation. Specimen shown is a Nycticebus pygmaeus vocal tract DLC 2901 available from http://www.morphosource.org

### Manual segmentation

Manual segmentation is usually carried out using a graphics tablet and the contours of each material of interest are manually traced by a researcher who is familiar with the characteristics of the material of interest. This technique is intrinsically reliant on the skill of the researcher carrying out the segmentation and is also extremely time consuming for large datasets, although it can be ameliorated slightly by the use of interpolation functions, so only every nth slice (e.g. every 5^th^ or 10^th^ slice) is fully manually processed. This functionality is available in all image processing software, although the user interface may differ. An example of manual segmentation is shown in figure 4.

**Figure 4.**
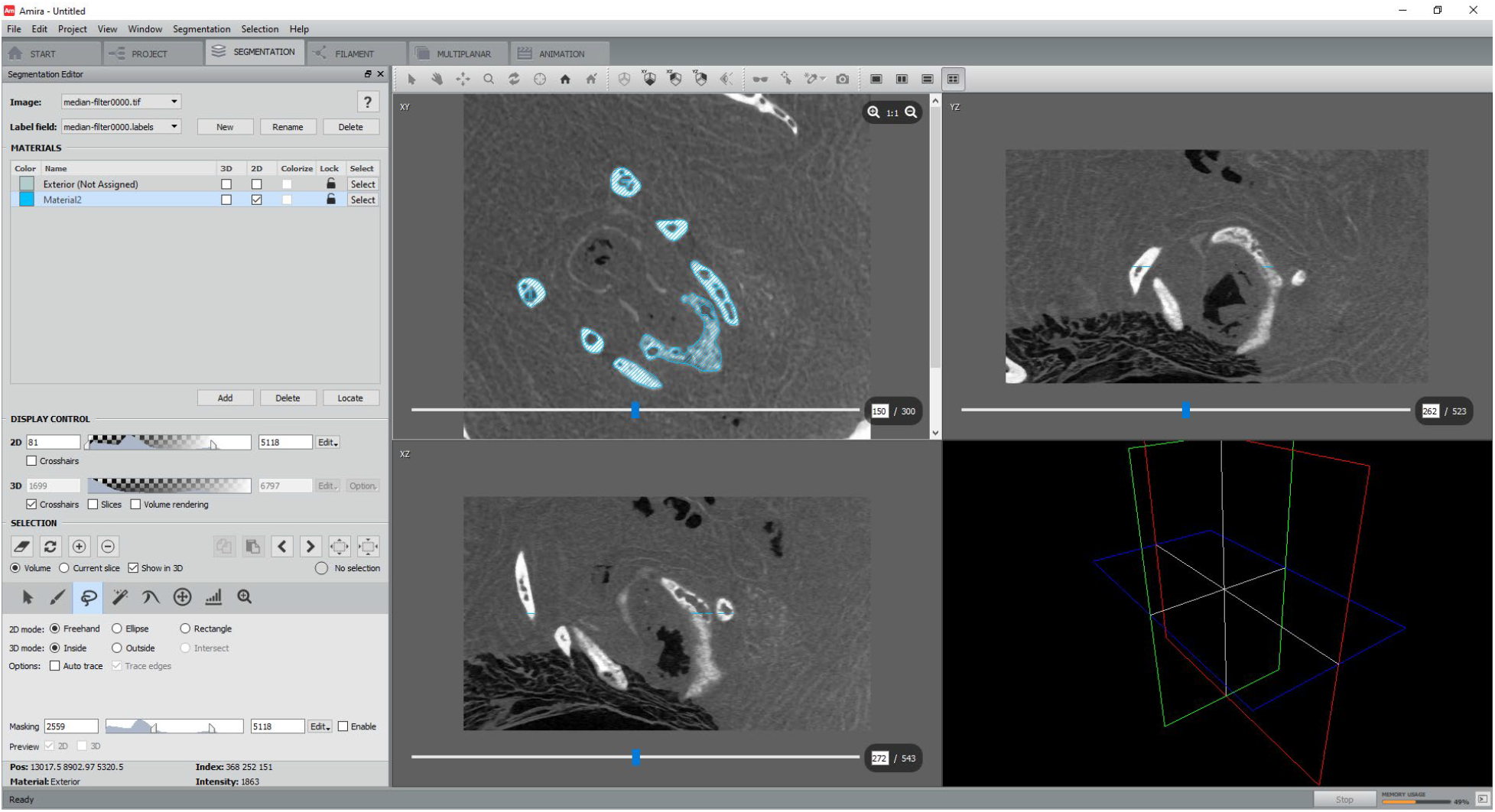
Manual segmentation. Specimen shown is a Nycticebus pygmaeus vocal tract DLC 2901 available from http://www.morphosource.org

### Label based segmentation

Label based segmentation is commonly used to ‘seed’ areas of interest and a contour is propagated until a significant difference in the material absorption is observed (within user set parameters such as kernel size and diffuseness of boundaries) See figure 5 for an example of this. In more recent applications, these approaches have been combined with machine learning, such as in (Arganda-Carreras et al., 2017; Glocker et al., 2013). Most user guided approaches utilize a variant of the Random Forest Algorithm for training (Breiman, 2001; Tin Kam Ho, 1998). Multiple different approaches to this, using different algorithms are available in freeware software (e.g. ImageJ, 3D slicer) and commercial packages (e.g, Avizo, DragonFly and Mimics). The Weka (Waikato Environment for Knowledge Analysis) segmentation method available in ImageJ is an example of supervised label based segmentation, augmented by machine learning and as such, is our preferred technique for the segmentation of complex anatomical and archaeological or paleontological data which may suffer from artefacts in scanning and material inhomogeneity (defined here as differences in material x-ray absorption). It has previously been tested on ground truth images applicable to geological samples and found to perform at least as well as other leading algorithms (Berg et al., 2018). It provides an easy to interpret overlay, (also known as a mask) on training datasets (see figure 6) and can be used to rapidly process multiple complex materials simultaneously.

**Figure 5.**
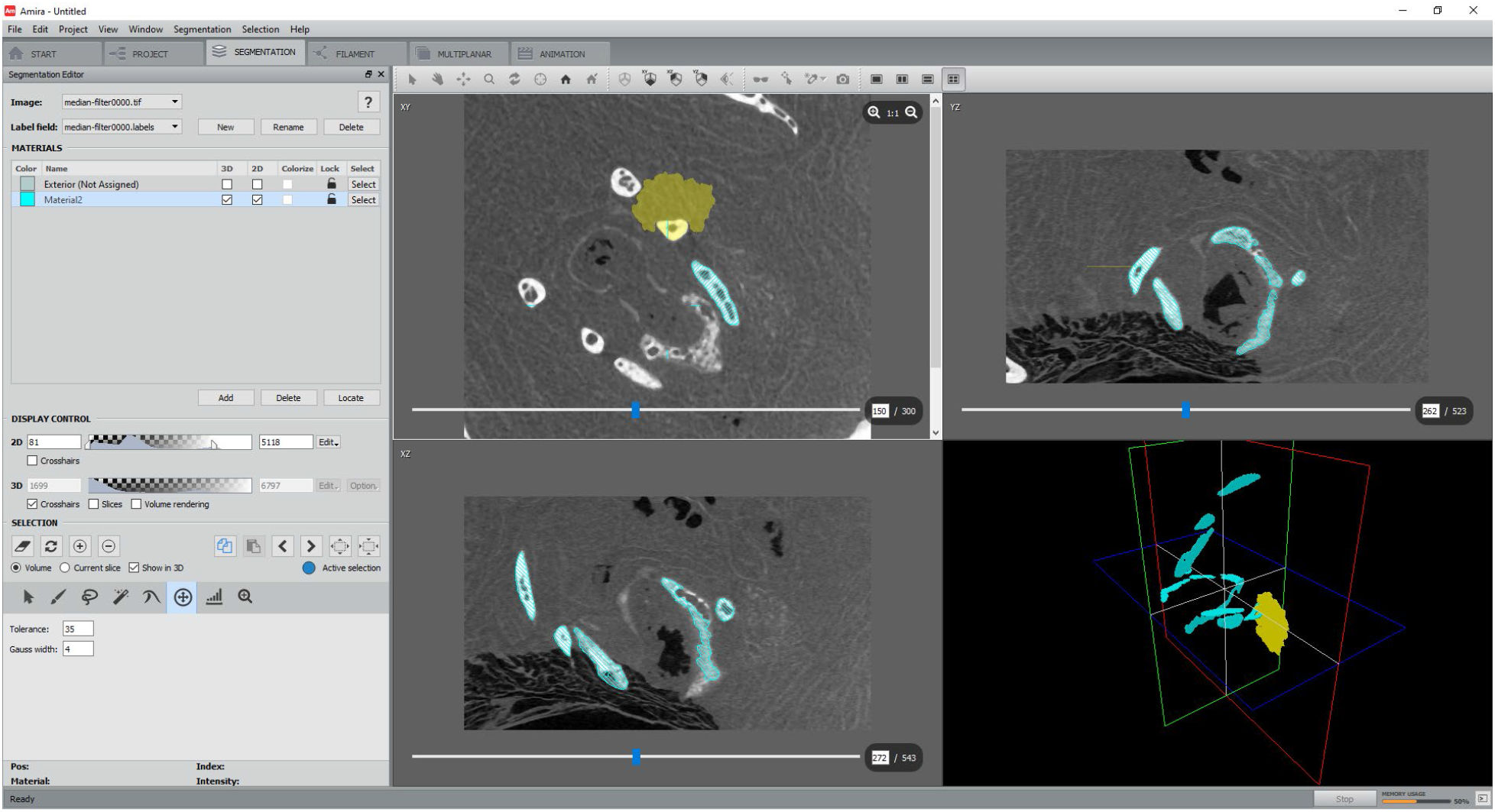
Segmentation by propagating contour. Specimen shown is a Nycticebus pygmaeus vocal tract DLC 2901 available from http://www.morphosource.org

**Figure 6.**
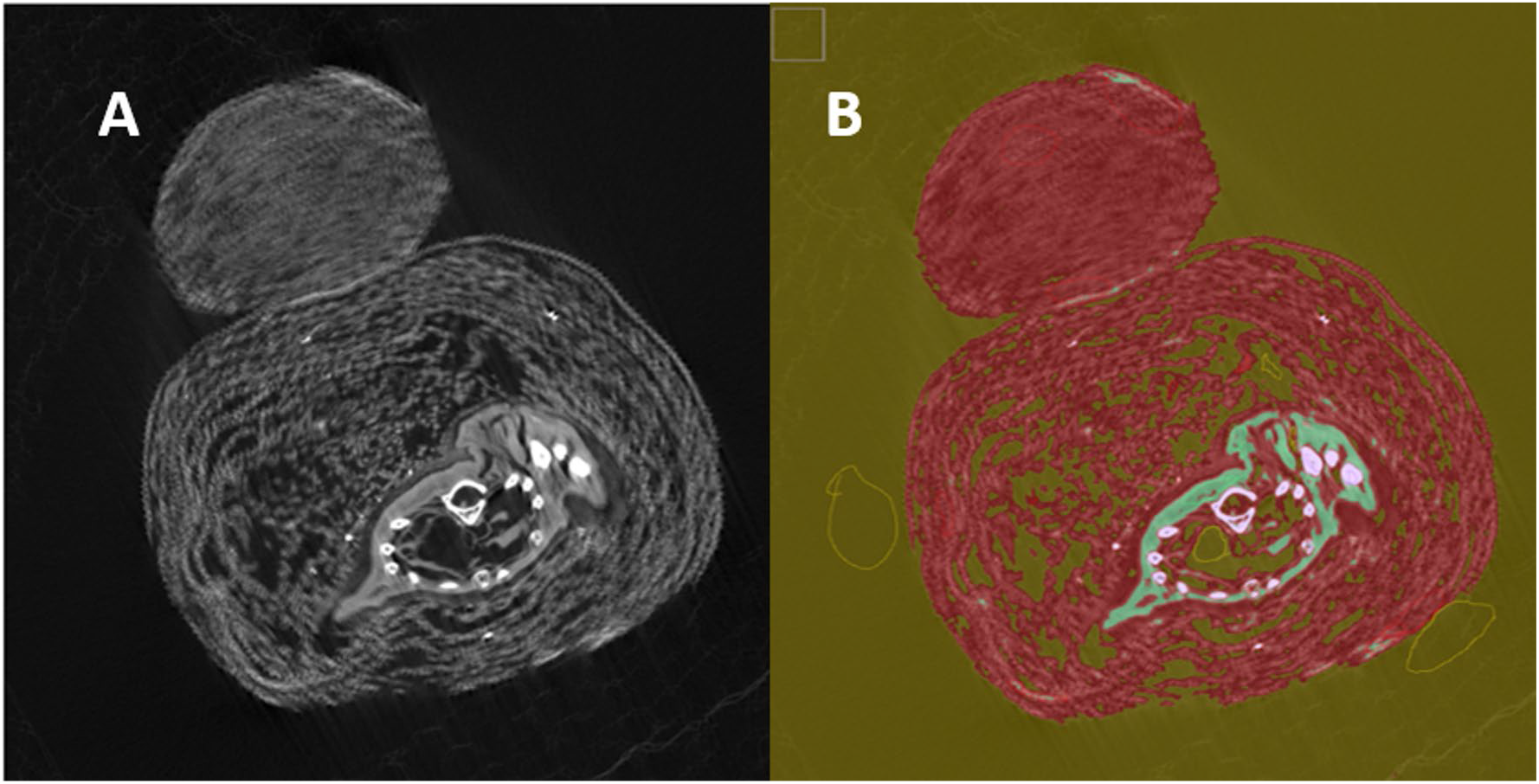
Weka segmentation (A=original, B=segmentation). Specimen shown is a small animal mummy in wrappings, Manchester Museum no. 6033

In summary, a significant roadblock to more rapid and precise advances in micro CT imaging in archaeological and evolutionary sciences is that the structures of interest are often non-homogenous in nature and until recently, have required extensive manual processing of slices. Recent advances in machine learning, combined with user-friendly interfaces mean that an acceleration of data processing is seriously possible, especially when combined with the potential to process data through either clusters or multiple GPUs. In this article, we demonstrate for the implementation of the Trainable Weka Segmentation (Arganda-Carreras et al., 2017), to typical micro CT data encountered in archaeological and evolutionary sciences. The Trainable Weka Segmentation is available through the FIJI fork of ImageJ (Schindelin et al., 2012).

We demonstrate the efficacy of these algorithms as applied to six distinct examples: an entirely synthetic dataset; micro CT scans of a machine wire phantom; a defleshed mouse tibia; a lemur vocal tract; a juvenile Neanderthal humerus (Kiik-Koba 2) and a small animal mummy. These represent a range of sample types commonly encountered by researchers working in imaging in evolutionary sciences and each presents different segmentation challenges. To further demonstrate the efficacy of Trainable Weka Segmentation, we compare this algorithm with other conventional and machine learning methods that have typically been used, including the half-maximum height protocol, k-means clustering and c-means clustering. We have chosen these to compare with Weka segmentation as they are all available through freeware (thereby maximizing user accessibility) and are relatively simple to implement. While commercial software such as Avizo/Amira (Thermo Fisher Inc., Dallas, TX), Mimics (Materialise N.V, Belgium); Matlab (Mathworks inc., Nattick, MA) and Dragonfly (ORS, Montreal) all offer conventional greyscale segmentation and some machine learning capability, we are interested in directly testing algorithms where the source code is publicly available.

We wish to address the following specific questions:

1. How effective are supervised and unsupervised machine learning segmentation algorithms at segmenting different types of material. Are they actually an improvement over a simple greyscale thresholding using half maximum height of the stack histogram?
2. Which machine learning algorithm is most effective for each type of scan?
3. How do the different algorithms effect the results of biomechanical analyses based upon these segmentations?
4. How well do these algorithms cope with scan noise, both real and artificially introduced?
5. Are two dimensional approaches more effective than three dimensional ones for unsupervised clustering?
6. How repeatable are segmentations performed using the Weka toolkit, both between users and within users?

## Materials & Methods

A series of datasets were selected which represent a range of material. These include entirely synthetized data (to ground truth methods); known size calibration artefacts and biological samples of interest to those working in archaeological science and evolutionary sciences. This allows users of the different segmentation methods tested here to be aware of their utility (and drawbacks) when applied to different data types.

### Synthetic Dataset

1. A synthetic dataset of 12 images of white triangle outlines on a black background was made. The original was kept as the ground truth. To simulate partial volume averaging and scanner noise, the following filters were applied in ImageJ: Noise: Salt and Pepper; Gaussian Blur of radius 2.0 pixels; Shadow from south (base) of the image. The original data, the data with noise added, and, all segmentations are included in the supplementary material. A render is in figure 7 with an arbitrary voxel depth of 10.

**Figure 7.**
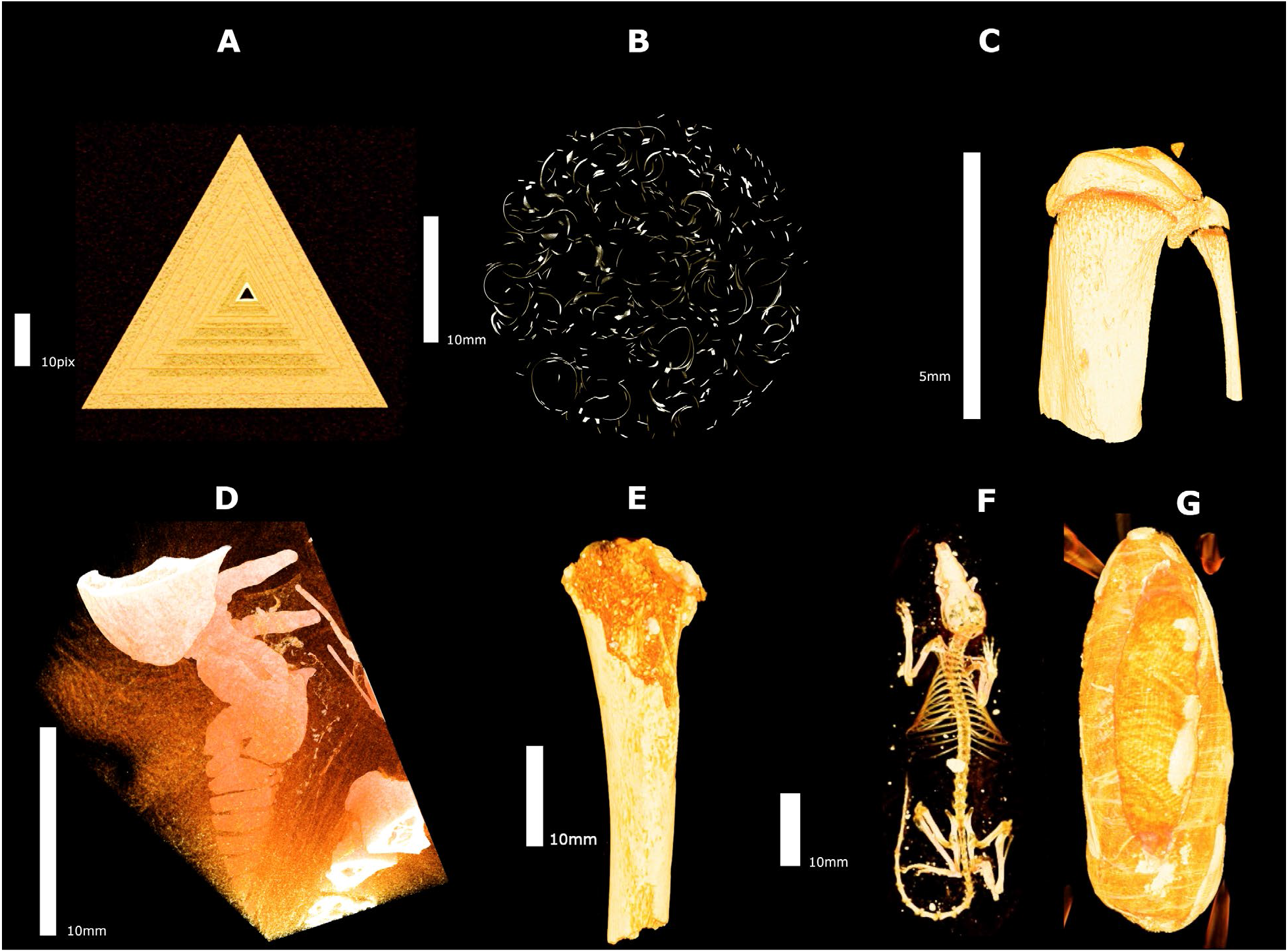
Volume renders of items analysed. A: Synthetic dataset; B: Machine Wire; C: Wild mouse type tibia section; D: Nycticebus pygmaeus vocal tract; E: Kiik Koba 2 partial humerus; E: Mummified rodent.

### MicroCT scans

2. A wire phantom object from (Dunmore et al., 2018). This is a coil of randomly crunched stainless steel wire of thickness 4mm.
3. A wild type mouse proximal tibia and from (Ranzoni et al., 2018). A 100×100×100 pixel cubic region of interest (ROI) was also selected from this scan for further analyses.
4. Lemur larynx -this is a scan of a wet preserved *Nycticebus pygmaeus* individual from the Duke Lemur Center, catalogue DLC_2901 and is more fully described in (Yapuncich et al., 2019).
5. A partial proximal humerus from a juvenile Neanderthal from the site of Kiik-Koba. It has been described in detail by (Trinkaus et al., 2016) and has matrix and consolidant adhering to it which obscure some more detailed aspects of its morphology.
6. An animal mummy, Manchester Museum number 6033. This is thought to be a shrew, based upon size of the wrappings and earlier medical X-Rays (Adams, 2015).

Full scan parameters are shown in Table 1 and volume renders of the tested datasets are shown in Figure 7.

**Table 1.** Scan parameters for each object

| *Object* | *Reference no.* | *Scanned with* | *Ke V* | *μA* | *Voxel size/μm* | *Filter* | *Scanning Medium* |
| --- | --- | --- | --- | --- | --- | --- | --- |
| *40μm thickness machine wire artefact* | *n/a* | *Skyscan1172* | *100* | *100* | *7.86* | *None* | *Air* |
| *Wild type mouse tibia* | *Number 6* | *Skyscan1172* | *49* | *200* | *5.06* | *Aluminium* | *70% EtOH* |
| *Nycticebus pygmaeus vocal tract* | *Duke Lemur Center 2901* | *Nikon XTH225* | *155* | *110* | *35.47* | *None* | *Air* |
| *Kiik-Koba 2 Neanderthal humerus* | *KunstKamera Museum* | *Custom MicroCT (MPKT-01)* | *135* | *35* | *30* | *None* | *Air* |
| *Rodent mummy in wrappings* | *Manchester museum 6033* | *Nikon XTH225* | *45* | *270* | *59.81* | *None* | *Air* |

These datasets were processed using a series of competing algorithms in ImageJ:

- Trainable Weka Segmentation
- Greyscale thresholding (using the half-maximum-height algorithm)
- K-means segmentation
- C-means segmentation

The datasets were also processed using localised fuzzy c-means segmentation, with pre-selection through k-means clustering using the Debian Linux package MIA -tools (Dunmore et al., 2018). We also provide a full step-by-step guide on using the Weka segmentation for MicroCT segmentation, including the settings chosen here as supplementary information (S2).

### Sample processing

All samples were subjected to segmentation using the Interactive Weka Segmentation editor plugin in ImageJ (Arganda-Carreras et al., 2017) with the following settings (adjusted after Somasundaram et al., 2018 who have applied this to medical images): Gaussian blur, Sobel Filter, Hessian Filter, Membrane projections (Thickness 1, patch size 10, difference of Gaussian filters, median filter (minimum sigma=1.0, maximum sigma=4.0); Kuwahara filter (Kuwahara et al., 1976). These filters help to counteract potential artefacts in the original scan slices and were found by (Somasundaram et al., 2018) to give the closest values to their ‘gold standard’ which was manual segmentation by a specialist.

Individual slices which contained all the materials of interest were trained using the Weka segmentation plugin. Briefly, areas containing each material of interest were selected using a graphics tablet and areas at the interface between materials were also selected (e.g. where a bone came into contact with air, near the edge of the bone was selected and added to the ‘bone’ label and a part near the edge of the air was selected and added to the ‘air’ label). This helped the algorithm to effectively select the correct labels at interfaces between materials. To propagate this label selection across the whole image, the Random Forest Algorithm was used, with 200 hundred trees. Although the use of fewer trees is more computationally efficient, the trade-off between efficiency and efficacy starts to plateau after ~250 trees (Probst and Boulesteix, 2018). All images were then segmented using the appropriate training dataset.

All stacks were processed on of two machines with 32GB RAM, PCIeM2 SSD and either a 6 core i7 at 3.6GHz (4.2GHz at boost) or an 8 core AMD 2700 at 3.2 GHz (4Ghz at boost). Due to the way the Java virtual machine is configured, graphic card parameters are not currently relevant for this workflow.

Two-dimensional k-means and c-means segmentation used the ImageJ plugin available from https://github.com/arranger1044/SFCM. Three-dimensional k-means segmentation used the plugin in the MorphoLibJ toolkit, version 1.41 (Argenda-Carreras et al., 2019, Leglang et al., 2016).

Spatial fuzzy c-means with k-means initialisation in MIA-tools used the settings recommended by Dunmore et al. (2018). This was processed as follows. Using Dunmore et al.’s (2018) recommendations, we pre-filtered the datasets with a median filter with a radius of 2 pixels and set the grid spacing at 7 pixels for all samples apart from the synthetic dataset and animal mummy which used a grid spacing of 12 pixels.

For visual purposes, 3D isosurfaces of each of the complete models were created using Avizo with no smoothing applied and the distances between the Weka segmentation and meshes generated with competing algorithms was visualised using the package Rvcg in R (Schlager, 2017).

To test the repeatability of the Weka segmentation, all 5 authors performed a segmentation of the synthetic dataset and 3 authors segmented the machine wire CT scan using identical settings in the Weka segmentation editor. The machine wire dataset was also segmented three times on separate occasions by the lead author. Statistics of overall accuracy (for the synthetic dataset) and intra and inter observer variation for both image stacks are reported.

### Statistical comparisons

The effects of the varying segmentation algorithms on real world results is the most important consideration, as it is sensible to anticipate that users will be most concerned about the accuracy of these. Given that many of the errors in segmentation were related to the artificial noise introduced into the dataset and are the type of noise that would be removed from a 3d model by a user, it was decided that automated measurement of the object in question was desirable. We wrote a script in Matlab 2019 (Mathworks inc.) which automatically measured the thickness of the main object in each slice in a radial fashion at 20 fixed positions per rotation, following (Bondioli et al., 2010). The resulting matrices were subtracted from the matrix for the original synthetic data to visualise where deviation occurred. Principal components analysis was also conducted on the matrices in Matlab 2019, and visualised in PAST v3 (Hammer et al., 2001) as a easier way to visualise how close each segmentation was to the original synthetic dataset. In the case of the wire phantom and the tibia ROI it was also possible to compare average wire/trabecular thickness and thickness distribution of the samples (with the wire also having a ground truth thickness of 40μm). One further real-world test was a comparison of the ellipsoidness (after Salmon et al., 2015) and degree of anisotropy in the trabecular ROI, to demonstrate what effect the segmentation would have on biomechanical analyses. All thickness and anisotropy calculations were calculated with BoneJ 2 (Doube et al., 2010). We also assessed the degree of bone volume in the trabecular ROI, as many publications use this as a proxy for levels of bone formation in response to weight bearing or mechanical stimulation (e.g. Acquaah et al., 2015; Farooq et al., 2017; Li et al., 2016; Milovanovic et al., 2017; Turner, 2002). Violin plots of deviation from the Weka segmentation in the Tibia ROI were generated using PAST statistics v3 (Hammer et al., 2001).

## Results

### Segmentation times

Segmentation times for Weka segmentation are listed for representative individual slices from each stack in table 2.

**Table 2.**
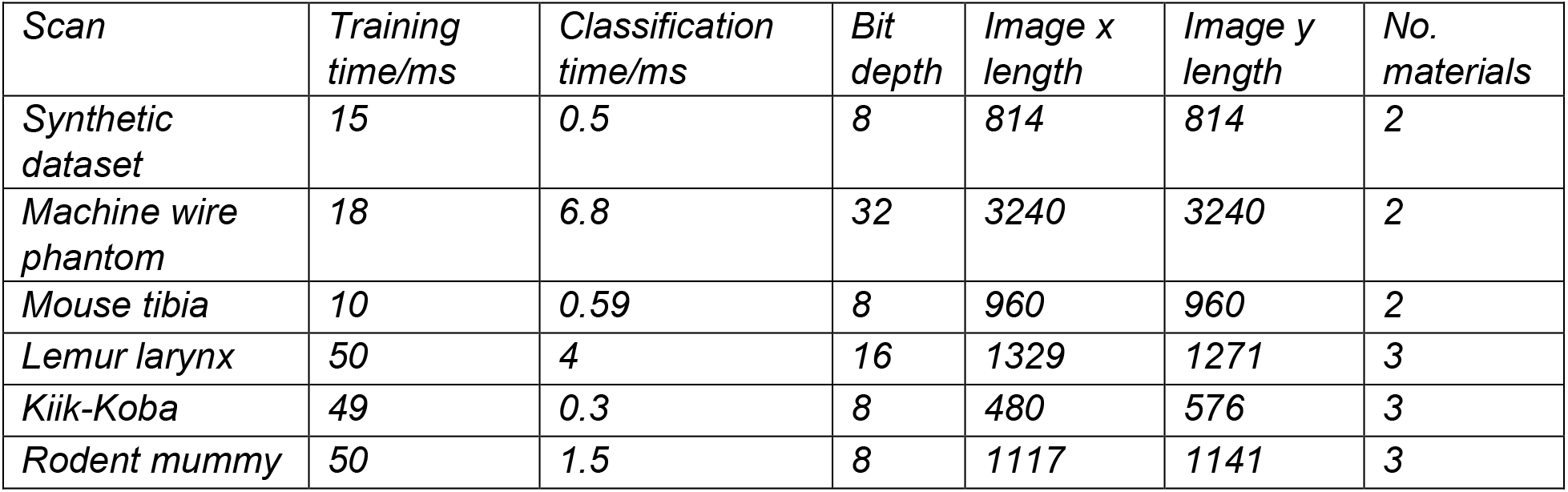
Segmentation times for each dataset

| *Scan* | *Training time/ms* | *Classification time/ms* | *Bit depth* | *Image x length* | *Image y length* | *No. materials* |
| --- | --- | --- | --- | --- | --- | --- |
| *Synthetic dataset* | *15* | *0.5* | *8* | *814* | *814* | *2* |
| *Machine wire phantom* | *18* | *6.8* | *32* | *3240* | *3240* | *2* |
| *Mouse tibia* | *10* | *0.59* | *8* | *960* | *960* | *2* |
| *Lemur larynx* | *50* | *4* | *16* | *1329* | *1271* | *3* |
| *Kiik-Koba* | *49* | *0.3* | *8* | *480* | *576* | *3* |
| *Rodent mummy* | *50* | *1.5* | *8* | *1117* | *1141* | *3* |

### Synthesised dataset

The majority of the data segmented relatively easily, but both two-dimensional k-means and local c-means struggled with the smaller triangles, where noise was closer to the dimensions of the object of interest (Figure 8). Repeatability and error of this segmentation is shown in table 3. Principal componenets analysis (Figure 9) demonstrated that the two-dimensional c-means and k-means segmentation resulted in large measurement errors due to noise. The most accurate segmentations (in terms of measurement accuracy) were those produced by Weka and three-dimensional local c-means. Interestingly, the most accurate overall segmentation produced by Weka segmentation had the worst precision in terms of linear measurements for this segmentation type (the outlier in figure 10), demonstrating that overall accuracy is probably not an ideal real-world statistic to report on when one is interested in particular features of images.

**Table 3.** Error of overall segmentations of synthetic datasets.

| Technique /user | Weka |  |  |  |  | 3D local c-means (MIA) | Half maximum height | 2D Kmeans | 2D local c-means | 3D kmeans |
| --- | --- | --- | --- | --- | --- | --- | --- | --- | --- | --- |
|  | JCD | LMM | MM | TL | TO'M |  |  |  |  |  |
| No. misclassified | 20362 | 27172 | 3745 | 19894 | 22712 | 17731 | 43270 | 483248 | 531470 | 541918 |
| % error | 0.245821 | 0.328035 | 0.045212 | 0.240171 | 0.274191 | 0.2141 | 0.522378618 | 5.834028711 | 6.416191 | 6.542324378 |

**Figure 8.**
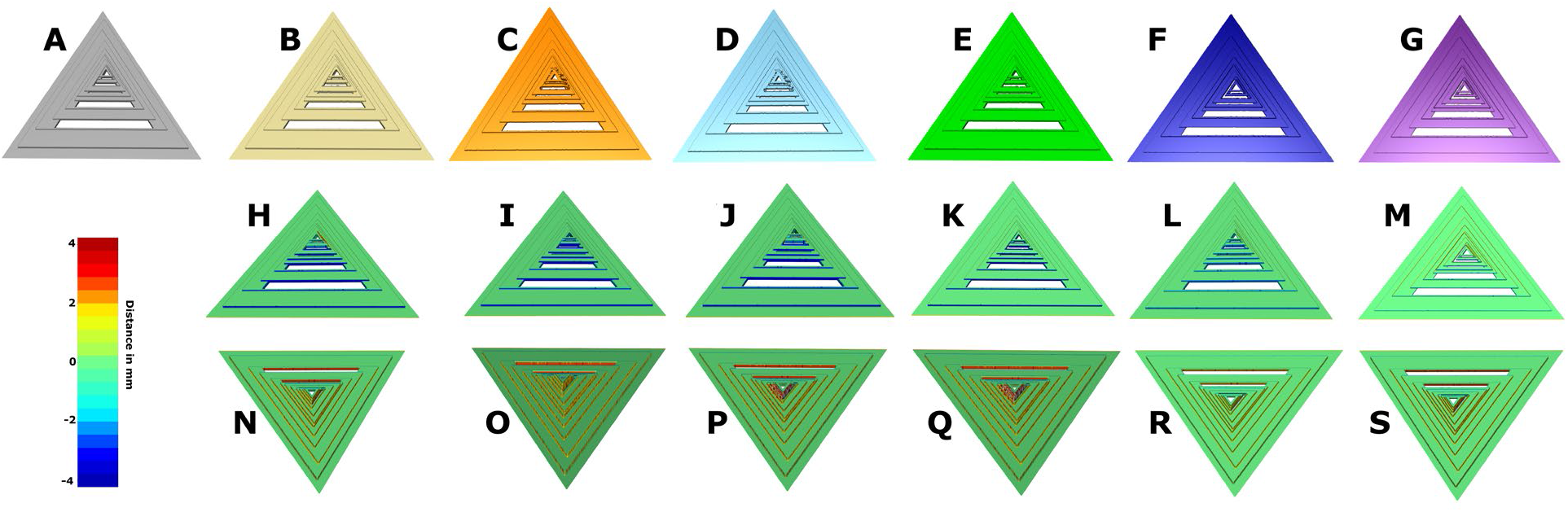
Synthesised data results. A: Original non-modified data; B: Weka Segmentation; C: local c-means; D: k-means; E: Half-maximum height; F: 3d k-means; G: 3d c-means. H-M: comparisons of above meshes with original data.

**Figure 9.**
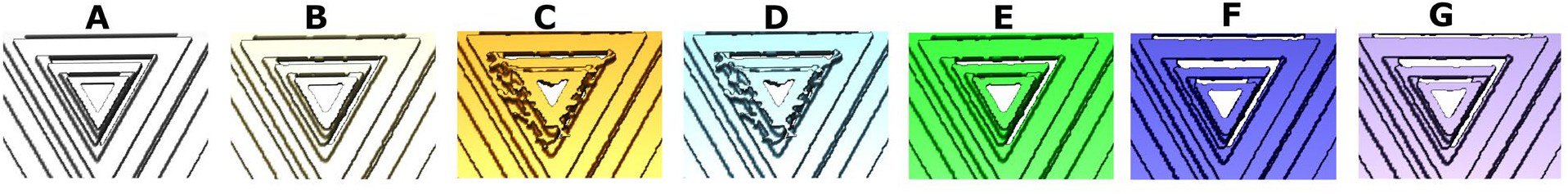
Closeups of the segmentations. A: Originl non-modified data; B: Weka Segmentation; C: local c-means; D: k-means; E: Half-maximum height; F: 3d k-means; G: 3d c-means.

**Figure 10.**
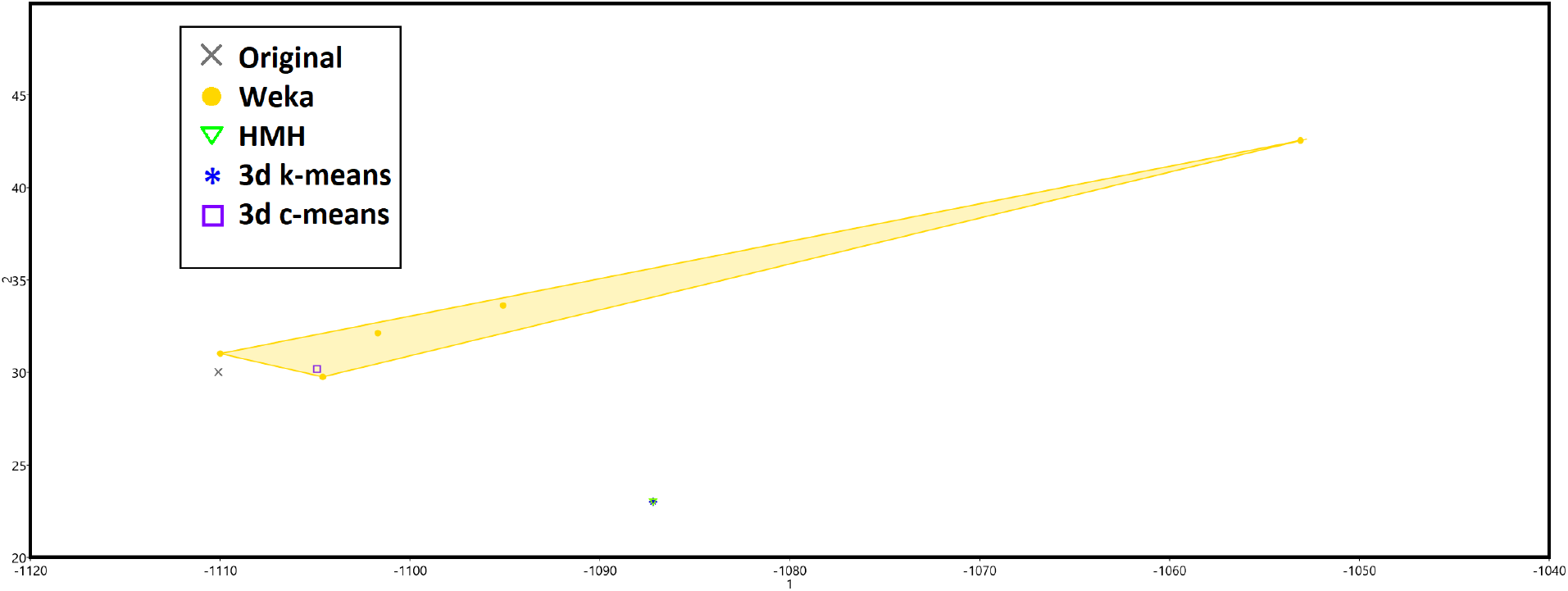
Principal Component Analysis of thickness measurements from all slices.

### Wire phantom

Three-dimensional local c-means yielded the most accurate segmentation, as shown in table 4. The Weka segmentation performed as well as the two-dimensional local c-means segmentation and improved some aspects of fine detail retrieval (figure 11). Intra-observer repeatability was within 5.63 microns, which is lower than voxel size.

**Table 4.**
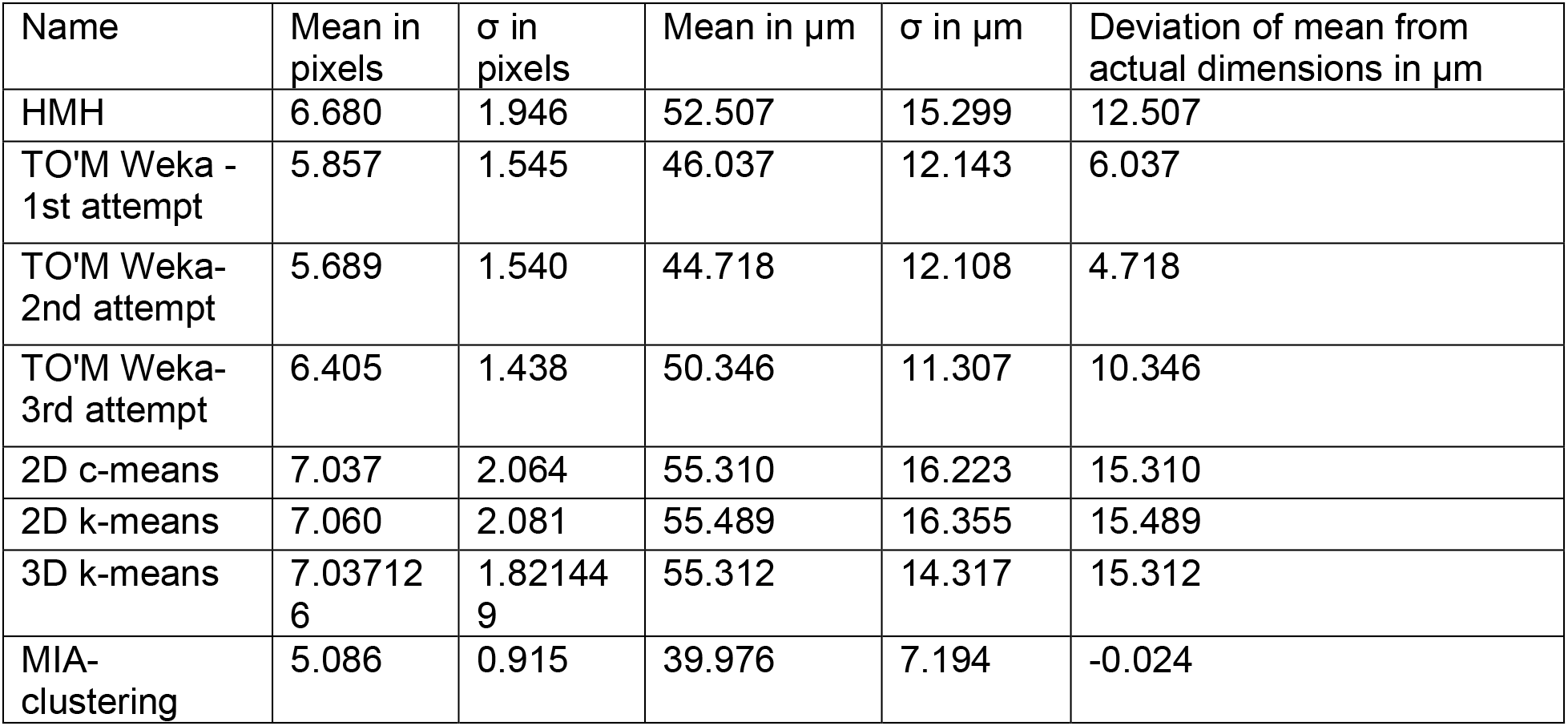
Accuracy and repeatability statistics for the different segmentations of the wire artefact.

| Name | Mean in pixels | $\sigma$ in pixels | Mean in $\mu\text{m}$ | $\sigma$ in $\mu\text{m}$ | Deviation of mean from actual dimensions in $\mu\text{m}$ |
| --- | --- | --- | --- | --- | --- |
| HMH | 6.680 | 1.946 | 52.507 | 15.299 | 12.507 |
| TO'M Weka - 1st attempt | 5.857 | 1.545 | 46.037 | 12.143 | 6.037 |
| TO'M Weka- 2nd attempt | 5.689 | 1.540 | 44.718 | 12.108 | 4.718 |
| TO'M Weka- 3rd attempt | 6.405 | 1.438 | 50.346 | 11.307 | 10.346 |
| 2D c-means | 7.037 | 2.064 | 55.310 | 16.223 | 15.310 |
| 2D k-means | 7.060 | 2.081 | 55.489 | 16.355 | 15.489 |
| 3D k-means | 7.037126 | 1.821449 | 55.312 | 14.317 | 15.312 |
| MIA-clustering | 5.086 | 0.915 | 39.976 | 7.194 | -0.024 |

**Figure 11.**
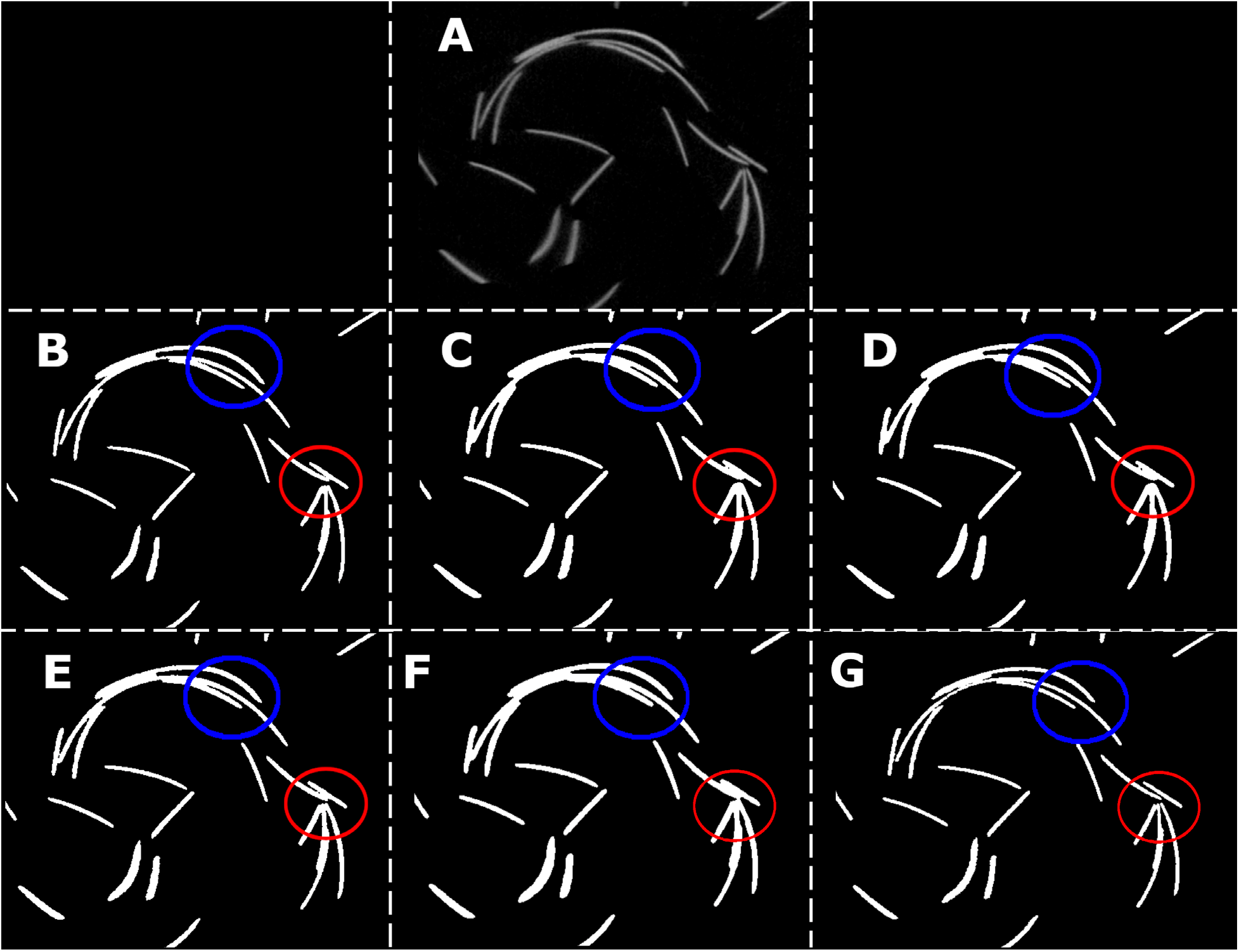
Comparisons of wire artefact segmentation. A: Original non-modified data; B: Weka Segmentation; C: local c-means; D: k-means; E: Half-maximum height; F: 3d k-means; G: 3d c-means. Modified after Dunmore et al., (2018) Red and blue circles indicating areas which require fine detail segmentation.

### Wild type mouse tibia

The Weka segmentation performed better than most of the other types of segmentation, with improved quality on fine features (Figure 12). The three-dimensional local c-means segmentation generated the smallest external cortex of the bone (so overall the object was subtly smaller) (Figures 13 and 14).

**Figure 12.**
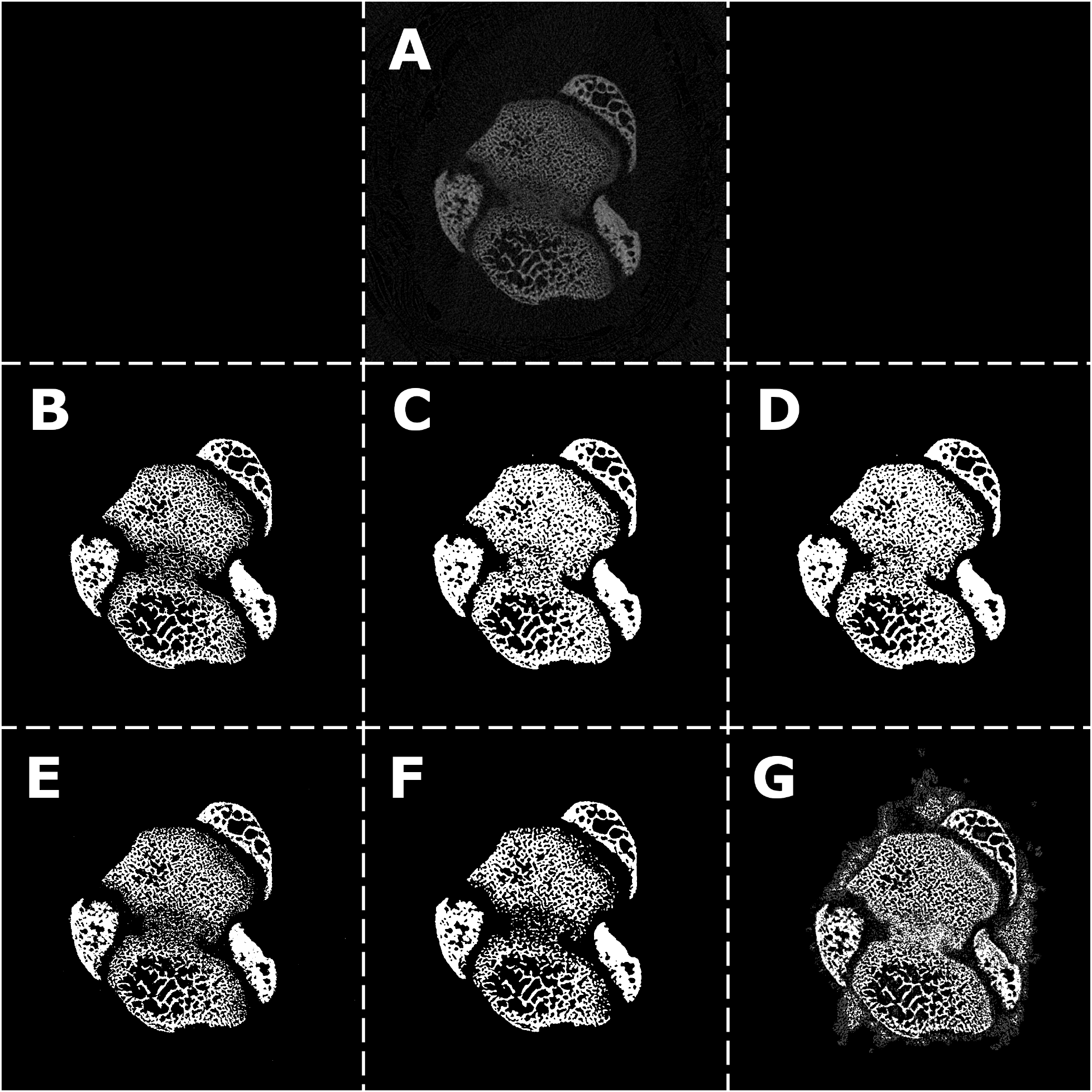
Comparisons of segmentations of mouse tibia-raw section views. A: Original; B: Weka Segmentation; C: local c-means; D: k-means; E: Half-maximum height; F: 3d k-means; G: 3d c-means.

**Figure 13.**
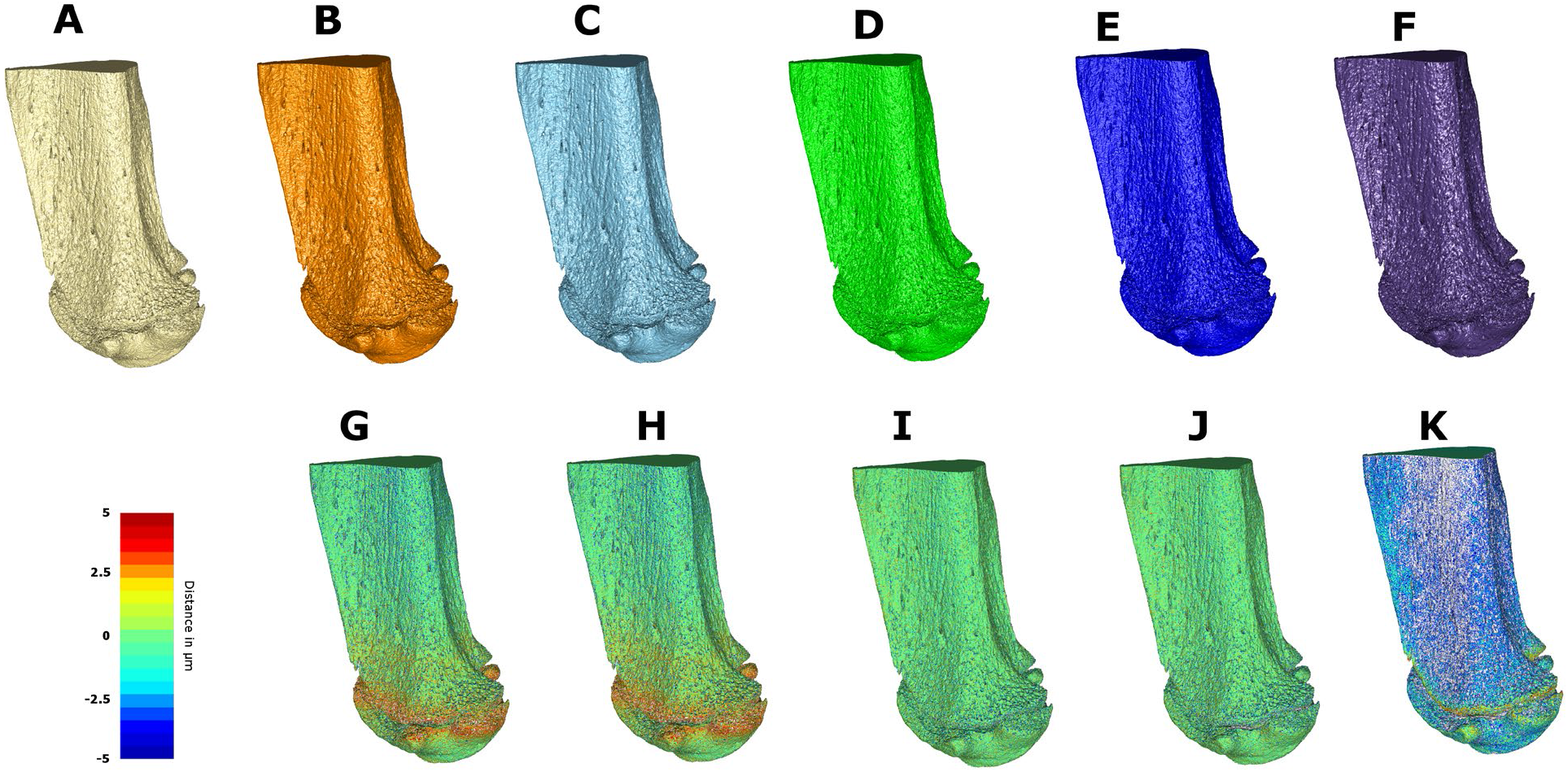
Comparisons of mouse tibia segmentation. A: Weka Segmentation; B: local c-means; C: k-means; D: Half-maximum height; E: 3d k-means; F: 3d c-means. G-K: comparisons of above meshes with original data. Blue to red scale, Blue indicates values which are concave compared with Weka; Red indicates areas that are convex compared with Weka.

**Figure 14.**
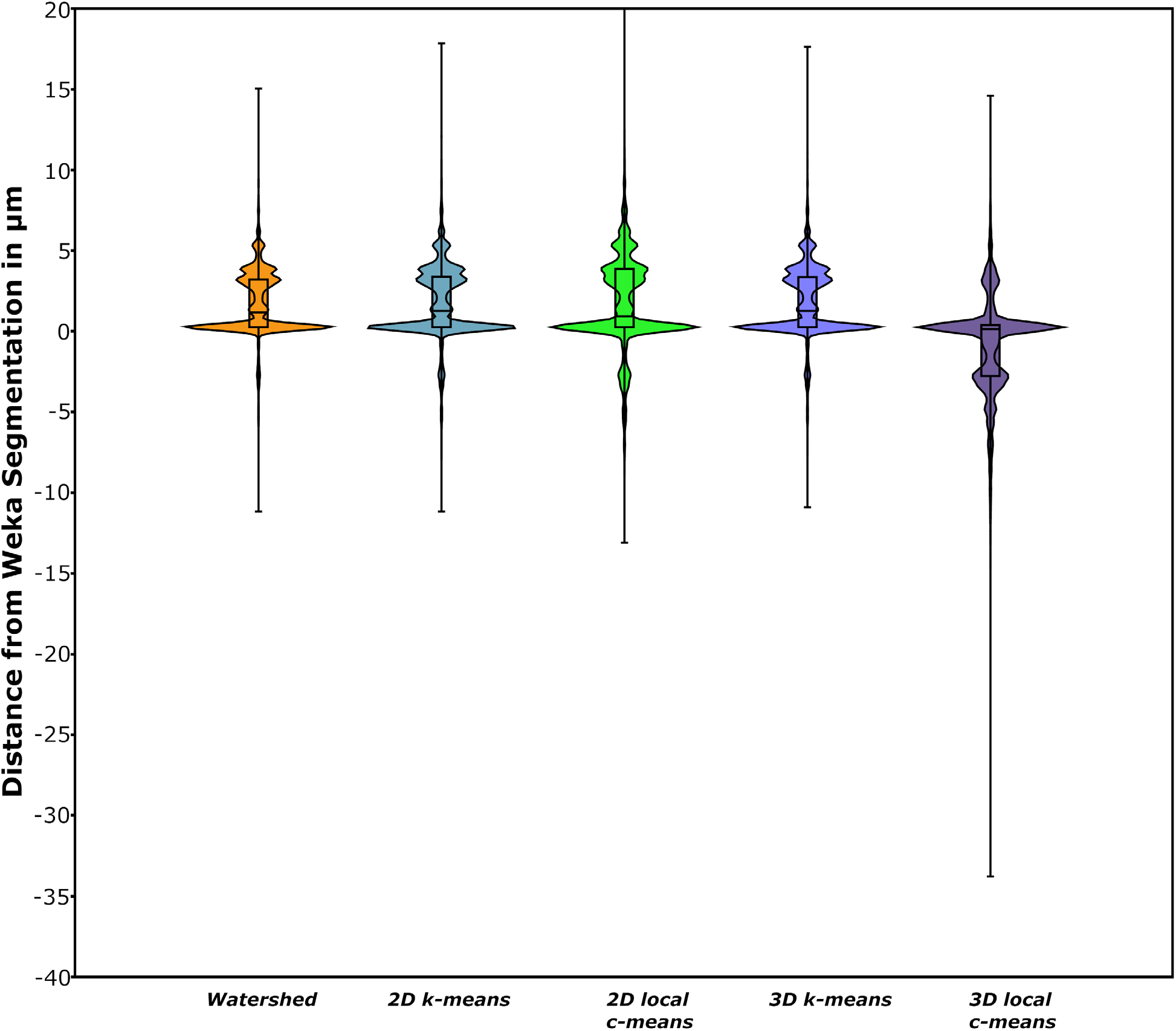
Violin plots with box and whiskers overlaid comparing deviations of segmentation from Weka result.

Violin plots indicate that alternative segmentation methods have subtly different distributions in terms of distance from the Weka segmentation (Figure 15). All suffer from arbitrary spiking in distribution of deviation of the data relative to the Weka segmentation.

**Figure 15.**
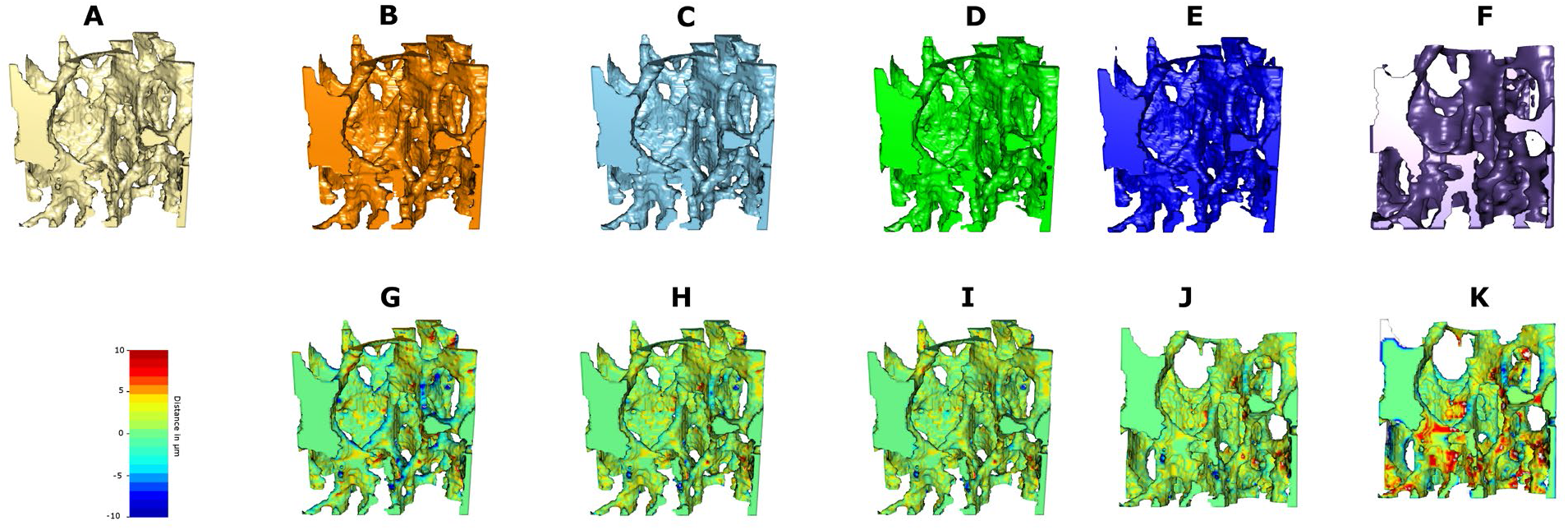
Comparisons of mouse tibia ROI segmentation. (A) Weka Segmentation. (B) 2D local c-means. (C) 2D k-means. (D) Half-maximum height. (E) 3d k-means. (F) 3d c-means. (G-K) comparisons of above meshes with original data. Blue to red scale, Blue indicates values which are concave compared with Weka; Red indicates areas that are convex compared with Weka.

It is also apparent that differing segmentation techniques have a marked effect on the degree of anisotropy detected in trabecular bone, with Weka tending towards more anisotropic structures. This may be because of the lack of spiking in the resulting segmentation when compared with the other methods here. The Ellipsoid Factor (a replacement for the Structure model index (Doube, 2015; Salmon et al., 2015) also varies considerably, with a difference of almost 4% between Weka and watershed segmentation (table 5). It is noticeable also that Weka segmentation classifies a relatively low percentage of bone and also trabecular thickness. Three-dimensional local c-means was very dependent on the grid size employed and tended to be very conservative at the grid size chosen (7) in terms of pixels classified as bone.

**Table 5.** Comparison of standard biomechanical results for different segmentation techniques for tibia ROI.

| *Segmentation technique* | *Degree of anisotropy (average of 10 runs)* | *% of foreground volume filled with ellipsoids* | *Median ellipsoid factor* | *% of ROI classified as bone* | *Trabecular thickness/$\mu\text{m}$* | *Trabecular thickness S.D. /$\mu\text{m}$* |
| --- | --- | --- | --- | --- | --- | --- |
| *Weka* | *0.74* | *28.4* | *-0.007* | *12.7* | *37.2* | *12.4* |
| *2D Local c-means* | *0.686* | *36.90* | *0.111* | *15.3* | *42.99* | *12.62* |
| *2D K-means* | *0.686* | *35.7* | *0.168* | *15.2* | *42.86* | *12.65* |
| *3D K-means* | *0.711* | *34.75* | *0.069* | *15.0* | *41.3* | *12.1* |
| *3D Local c-means* | *0.628* | *31.39* | *0.137* | *16.8* | *39.55* | *11.05* |
| *Conventional watershed* | *0.707* | *32.33* | *0.084* | *15.1* | *41.19* | *12.04* |

### Lemur larynx

The Weka segmentation was able to account for the ring artefacts in the scan and successfully segmented the materials of interest. It was also more successful at segmenting the finer structures in the larynx (see Figure 16 and 17). It also generated much cleaner data than all other segmentations. Three dimensional local c-means created a ‘halo’ around the bone, which means that any calculations of relative proportions of tissues would be likely to be incorrect.

**Figure 16.**
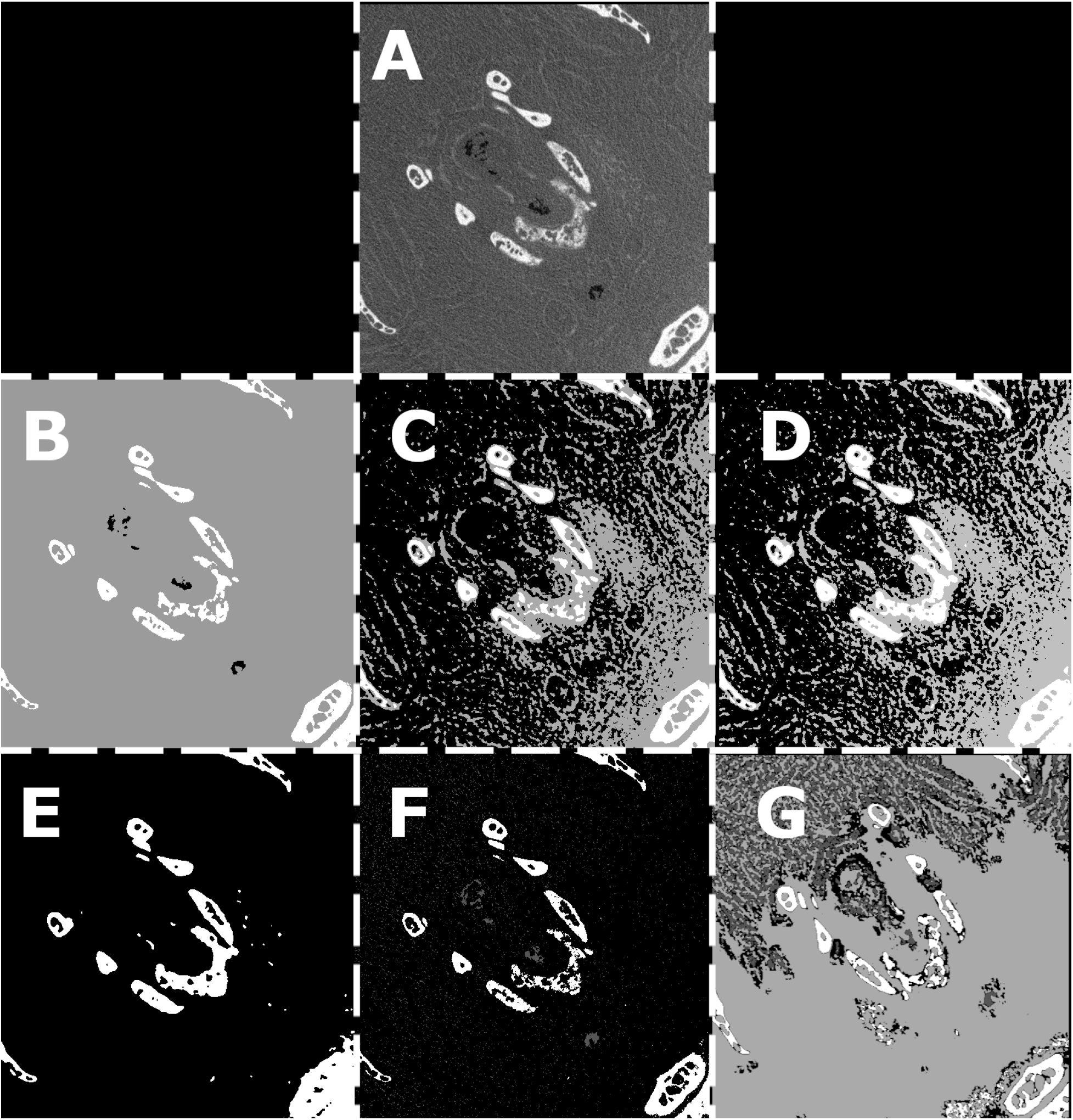
Comparisons of segmentations of Nycticebus pygmaeus -raw section views. (A) Original slice. (B) Weka Segmentation. (C) 2D local c-means. (D) 2D k-means. (E) Half-maximum height. (F) 3d k-means. (G) 3d c-means

**Figure 17.**
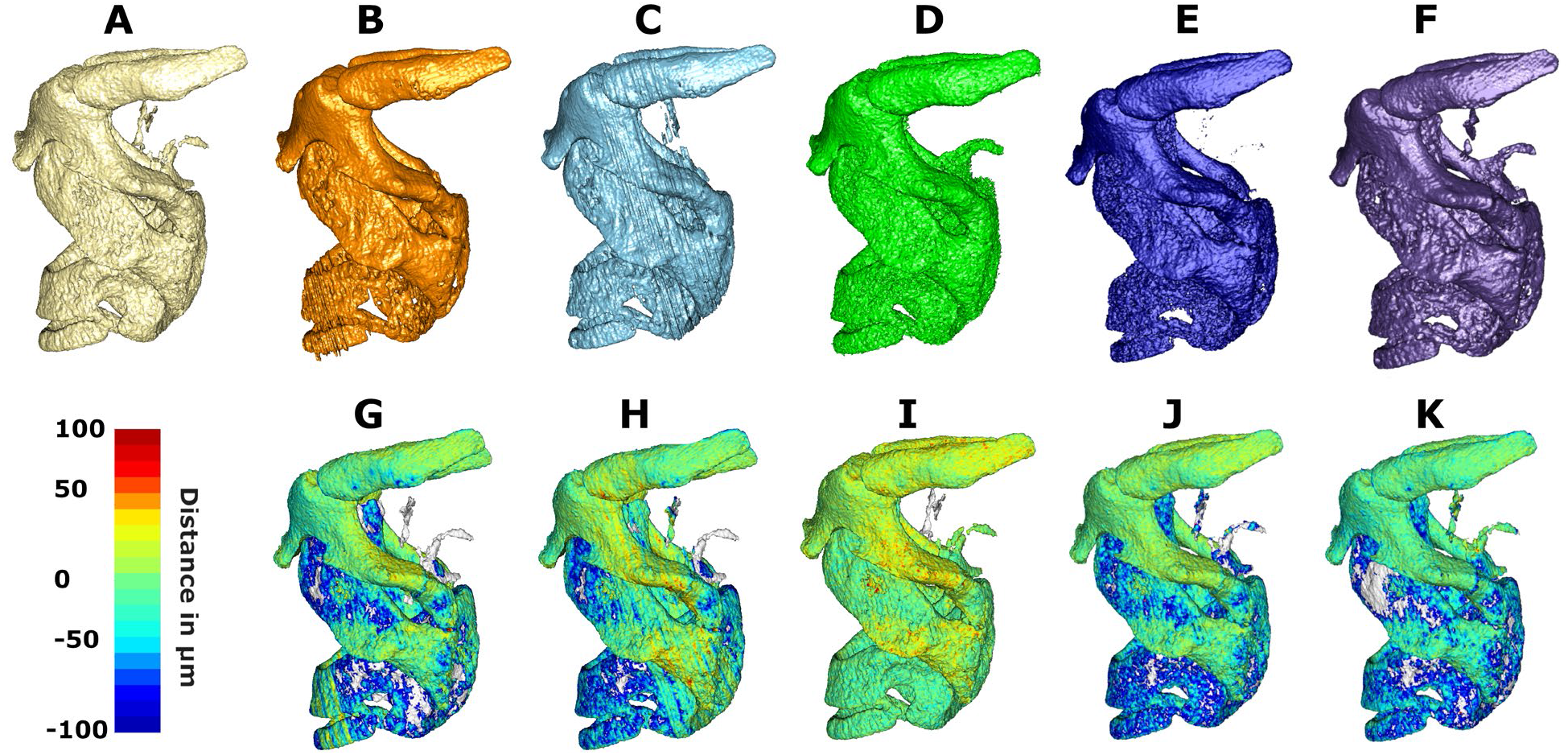
Comparison of Nycticebus pygmaeus scan segmentations. (A) Weka Segmentation. (B) 2D local c-means. (C) 2D k-means. (D) Half-maximum height. (E) 3d k-means. (F) 3d c-means. (G-K) comparisons of above meshes with original data. Blue to red scale, Blue indicates values which are concave compared with Weka; Red indicates areas that are convex compared with Weka.

### Kiik-Koba Neanderthal humerus

The Weka segmentation was able to track trabecular structure successfully, without eroding the material. It also was able to take into account the slight ‘halo’ effect on the bone/air interface, which conventional segmentation used to create an external border of the matrix material. The c-means and k-means segmentation both created this ‘halo’ like border (Figures 18 and 19). The watershed segmentation performed very well in this instance as the scan was relatively clean with good contrast between the different materials. The three-dimensional local c-means segmentation was unable to compensate for the ring artefacts in the scan and classified these with the fossil matrix. It also had the ‘halo’ seen in two dimensional c-means.

**Figure 18.**
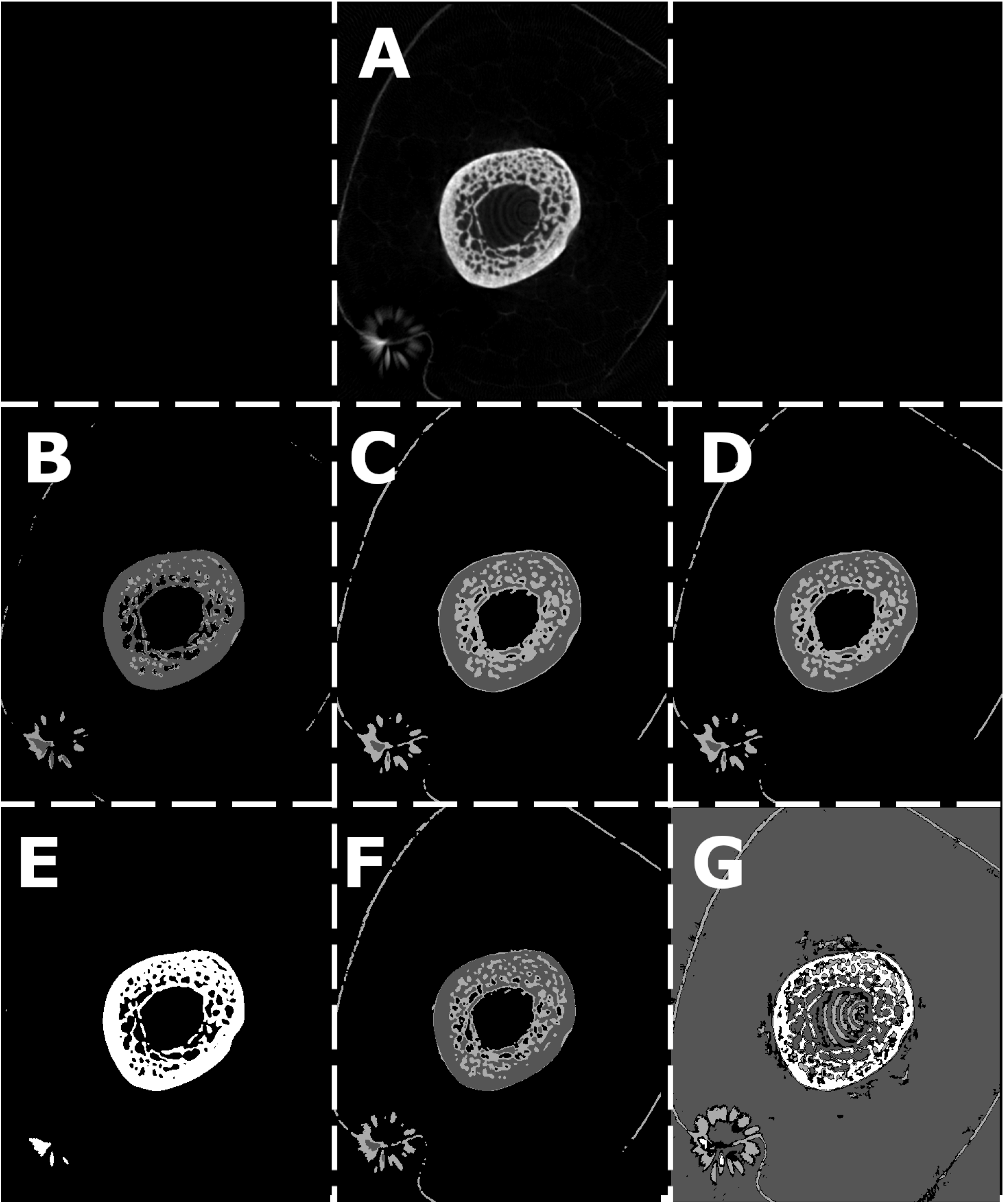
Comparisons of segmentations of Kiik-Koba 2 Neanderthal humerus -raw section views. (A) Original slice. Weka Segmentation. (C) 2D local c-means. (D) 2D k-means. (E) Half-maximum height. (F) 3d k-means. (G) 3d c-means

**Figure 19.**
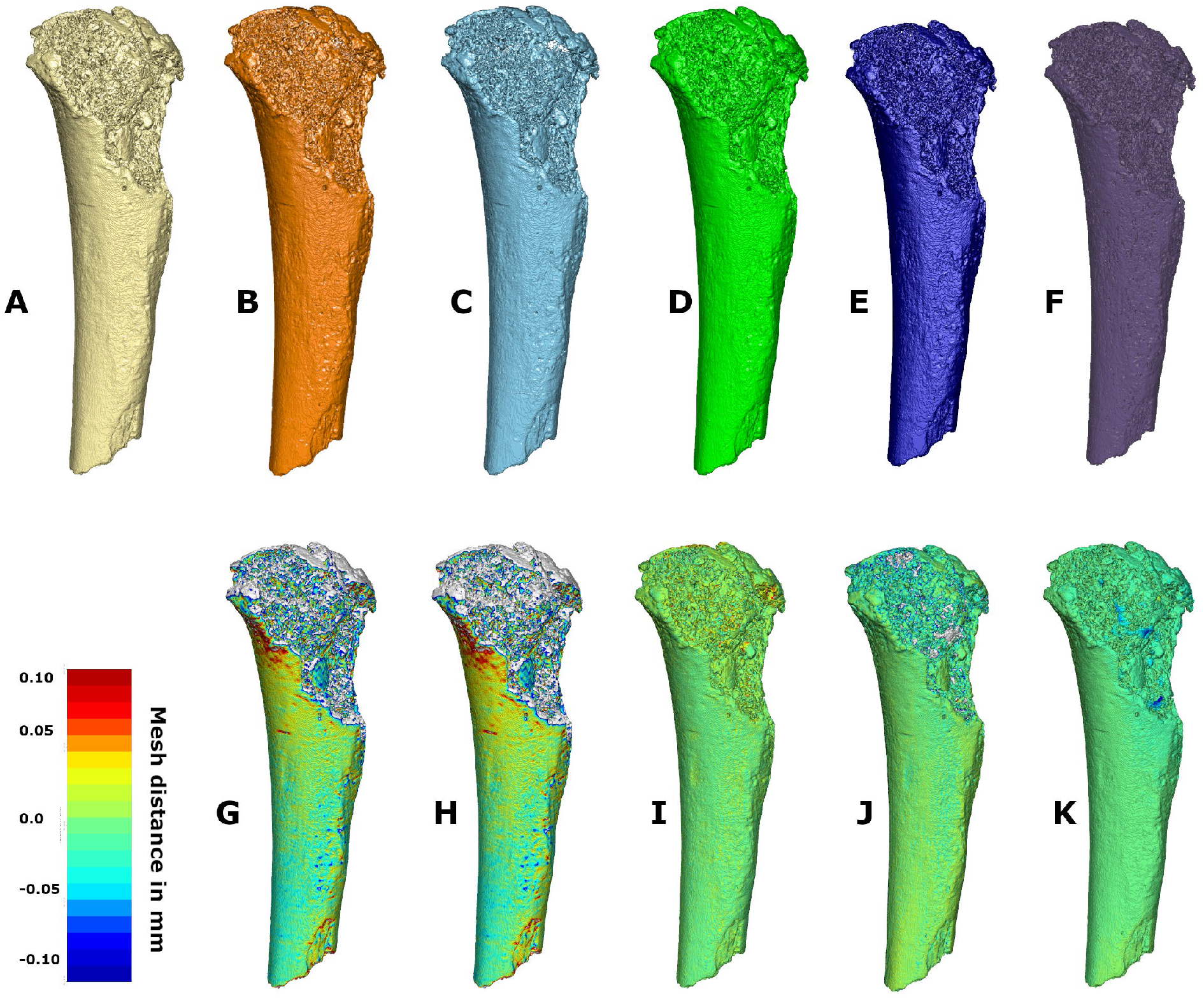
Comparison of Kiik-Koba 2 Neanderthal humerus scan segmentations. (A) Weka Segmentation. (B) 2D local c-means. (C) 2D k-means. (D) Half-maximum height. (E) 3d k-means. (F) 3d c-means. (G-K) comparisons of above meshes with original data. Blue to red scale, Blue indicates values which are concave compared with Weka; Red indicates areas that are convex compared with Weka.

### Animal mummy

The Weka segmentation was able to detect the majority of the skeletal features and also discriminate the mummified tissues from the outer wrappings. There were a few artefacts around the front paws and mandible which would require some manual correction to fully delineate the structures. The smoothing steps introduced in the segmentation were able to remove many of the scanning artefacts which made borders of materials harder to resolve with conventional methods. Two dimensional K-means segmentation was able to discriminate materials relatively well also, although it did misclassify several slices. Two dimensional localised C-means was unsuccessful in several slices, missing bone material entirely, where other segmentation techniques succeeded. Three-dimensional c-means created a halo around bone once again. The conventional half maximum height segmentation performed extremely well at identifying the bone in this instance (see figures 20 and 21).

**Figure 20.**
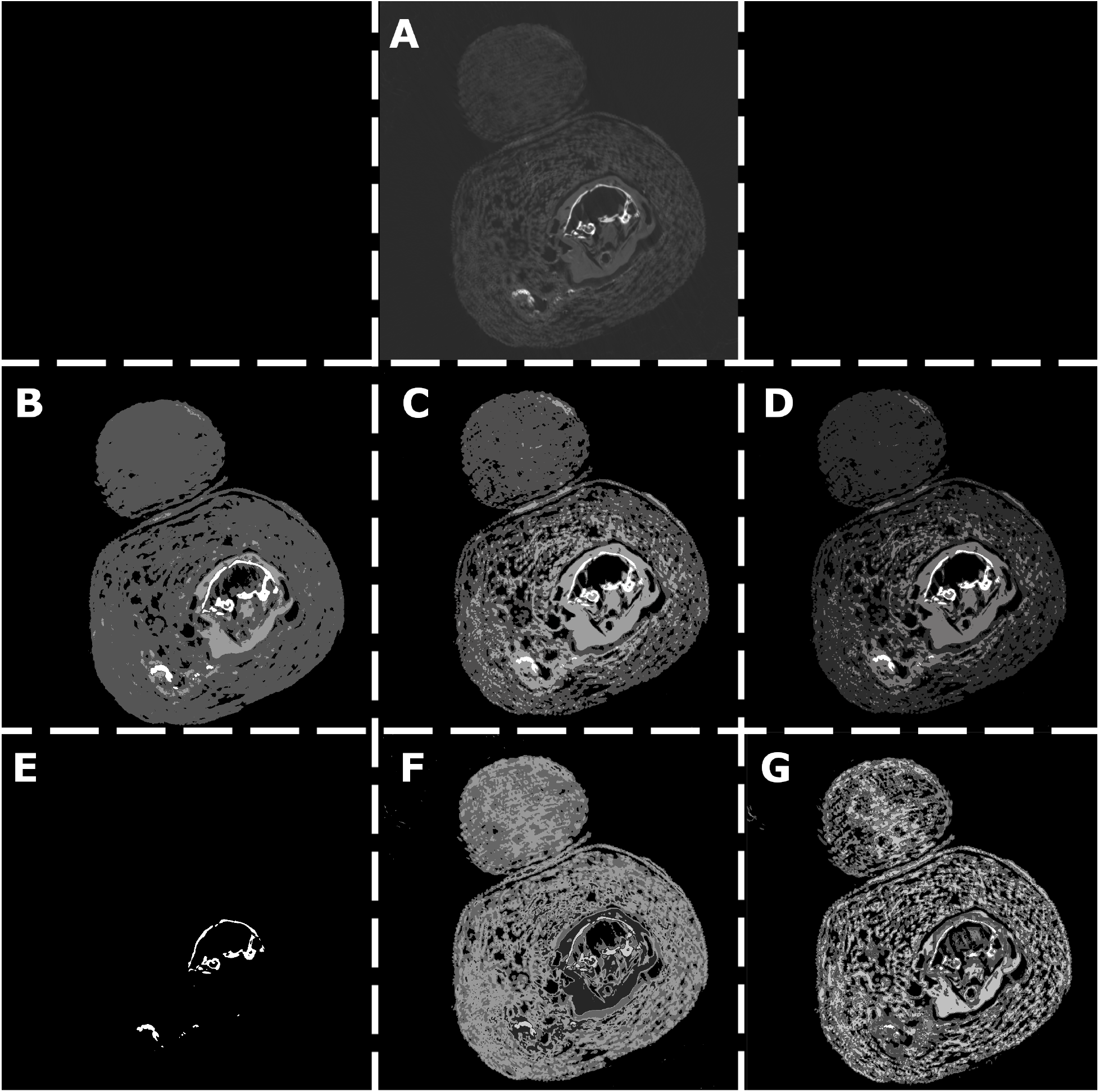
Comparisons of segmentations of animal mummy -raw section views. (A) Original slice. (B) Weka Segmentation. (C) 2D local c-means. (D) 2D k-means. (E) Half-maximum height. (F) 3d k-means. (G) 3d c-means.

**Figure 21.**
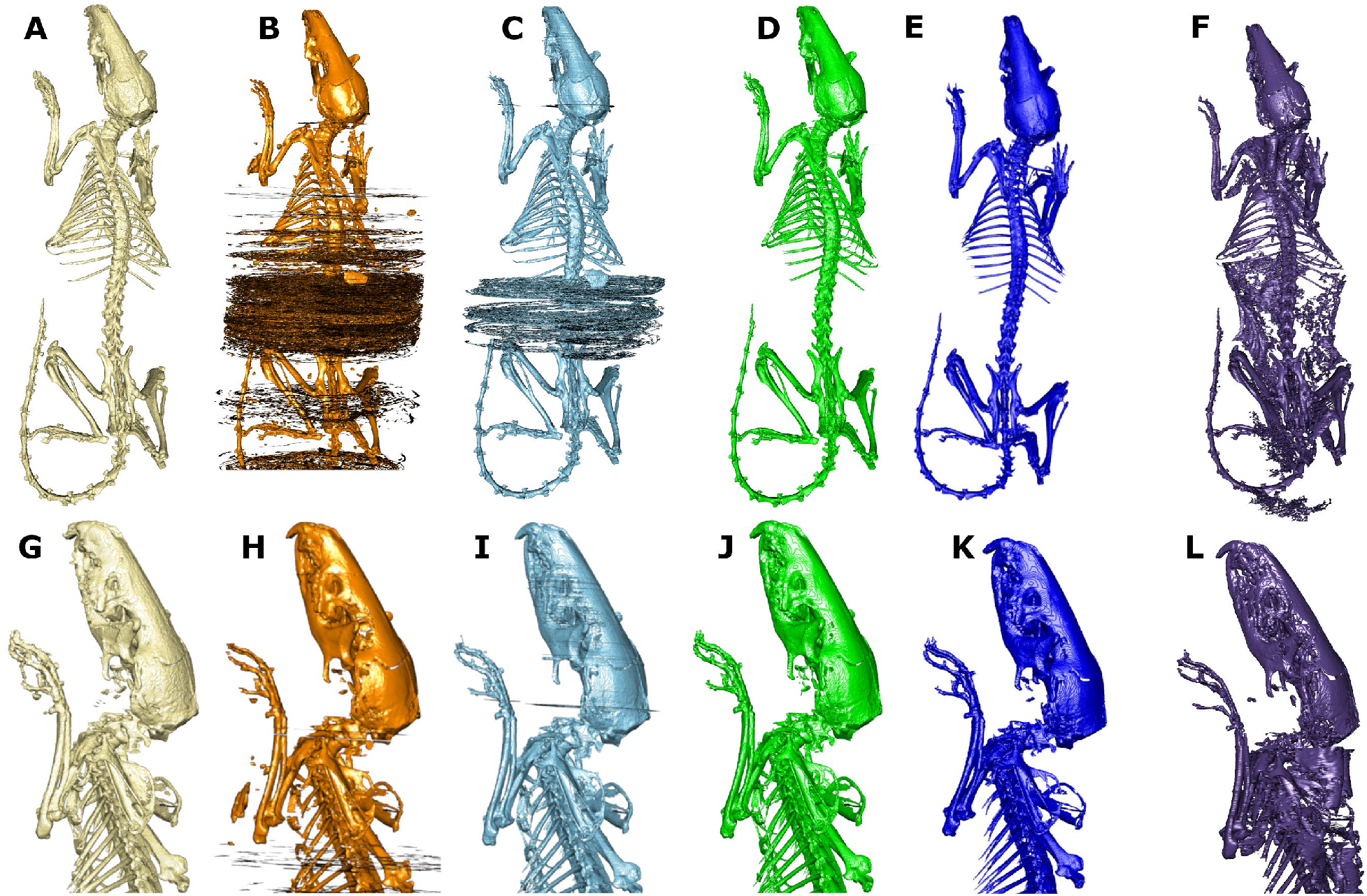
Comparison of animal mummy scan segmentations. (A) Weka Segmentation. (B) 2D local c-means. (C) 2D k-means. (D) Half-maximum height. (E) 3d k-means. (F) 3d c-means. (G-K) comparisons of above meshes with original data. Blue to red scale, Blue indicates values which are concave compared with Weka; Red indicates areas that are convex compared with Weka.

## Discussion

Firstly, we address the specific questions raised in our introduction.

1. How effective are supervised and unsupervised machine learning segmentation algorithms at segmenting different types of material? Are they actually an improvement over a simple greyscale thresholding using half maximum height of the stack histogram? Overall, the machine learning algorithms were effective at segmenting the brightest material in every dataset. Two-dimensional k-means and local c-means regularly flipped the order of the labels, suggesting that they are not particularly effective for the segmentation of whole image stacks. Three-dimensional local c-means struggled with artefacts in scans (e.g. ring artefacts in the Kiik-Koba 2 scan). These results suggest that overall, the machine-learning based algorithms are an improvement on greyscale thresholding for processing of data and should be preferred. Overall, the Weka segmentation algorithm generated improved results when compared with standard tools available in ImageJ and was especially good at tracking fine structures throughout the samples. All the algorithms tested apart from Weka and three dimensional local c-means seem to struggle to differentiate between materials when contrast is low and fine details approach the resolution of the scans (e.g. they struggle with details that are around 10 microns in size, when the scan is 5 microns in resolution). Although it did not perfectly segment out the bone in the rodent mummy sample, it is a noted improvement on current methods, and is extremely easy to implement. The segmentations produced compared favorably with three-dimensional c-means as applied in MIA-tools (Dunmore et al. 2018) and has the added advantage that ImageJ is widely available, easy to learn, platform independent and free to download and extend using a variety of common scripting languages. It also has the advantage over the MIA-clustering algorithm in that it can compensate for material interface artefacts in scans (Dunmore et al., 2018), especially ring artefacts. A suggestion for further processing of data from especially noisy scan data is that the user applies the ‘despeckle’ filter in imageJ, either on the original data, or the resulting segmentation. We would advise researchers if they are to use unsupervised clustering techniques to segment their data, then it is best to use three-dimensional approaches such as the tools provided in the MorphoLibJ toolkit (Argenda-Carreras et al., 2019, Leglang et al., 2016) or MIA-tools (Dunmore et al.).
2. Which machine learning algorithm is most effective for each type of scan? Three-dimensional local c-means was most effective for segmentation of a single-phase material in air (e.g. the wire phantom and the synthetic phantom). Weka was most effective for segmentation of multiple materials simultaneously. In term of absolute accuracy, the three-dimensional local c-means performed the best in the wire segmentation, and we were able to exactly replicate the results of Dunmore et al., (2018). The clustering results when simple objects that had only two material types yielded the most satisfactory segmentations. The ‘halo effect’ noted in two-dimensional k-means and both variants of local c-means clustering is however an issue when trying to classify a range of materials within one dataset (such as muscle, fat, and bone in a scan of a wet tissue sample) as this may be of equal interest to researchers. We surmise that the reason for this artefact is linked to the partial volume averaging effect at material boundaries (Goodenough et al., 1981). Essentially, at the boundary, the probability of the voxel belonging to either material is equal and therefore it assigns a null material as it has failed to classify the data The repeatability of the Weka segmentation between observers is also within acceptable margins (<0.2% difference between 5 observers) and intra-observer repeatability over a single complex sample is also acceptable.
3. How do the different algorithms effect the results of biomechanical analyses based upon these segmentations? The algorithm has a marked effect on results of biomechanical analyses, with a difference in mean trabecular thickness of ~15% and a difference in anisotropy of up to ~7%. Verdelis et al., (2011) previously cautioned that inter system microCT results at high resolutions may not necessarily be comparable. Our results with the trabecular ROI suggest that as well as differences in scanning system choice, segmentation algorithms (and the parameter applied within them) should also be carefully chosen and explicitly stated by researchers (including settings used), as this can have a large effect on results. It also appears to contradict the results of Christiansen (2016), who found that different watershed methods did not really affect results at high resolutions. This is an area which warrants more investigation.
4. How well do these algorithms cope with scan noise, both real and artificially introduced? Three-dimensional local-c-means struggles with scan noise, greyscale thresholding and two-dimensional c-means and k-means do not cope well with speckles in the data, when contrast is low and fine details approach the resolution of the scans as stated above. Weka copes very well with scan noise.
5. Are three dimensional approaches more effective than two dimensional ones for unsupervised clustering? In the case of the synthetic data, the thickness results obtained were in an order of magnitude more accurate than two-dimensional ones. For the wire artefact, three-dimensional local c-means produced by far the most accurate segmentation, but three-dimensional k-means did not appear to change much. For other datasets, three-dimensional approaches were much more effective as they did not suffer from ‘flipping’ of label order, thereby reducing post-processing significantly. Overall, three-dimensional clustering approaches appear to be more effective and as such, are recommended over two-dimensional ones.
6. How repeatable are segmentations performed using the Weka toolkit, both between users and within users? Weka segmentation shows a good repeatability both between users and within users. Inter-observer error on the synthetic dataset was less than 0.2%. Intra-observer repeatability over a single complex sample is also acceptable, being less than voxel size.

More generally, the Weka algorithms can be applied to a wide range of image types, as it was originally developed for microscopy (Arganda-Carreras et al., 2017). We have tested this on image datasets from 8 to 32-bit depth. Given workflow constraints, most images used in everyday analysis will be either 8 or 16-bit. DICOM data requires the conversion to TIFF or other standard image formats to processing with Weka. ImageJ/Weka segmentation is multi-platform and has a user-friendly GUI. This make it an ideal toolbox to teach researchers (who may be unfamiliar with the subtleties of image processing) a fast and free way to process their CT data. Key parameters to observe are to use a fast CPU with multiple cores, which will enable users to fully leverage multi-threading; as well as the use of fast hard drives (preferably solid-state drives) if working on a desktop. Training the segmentation using fine structures will also improve delineation of edge features. Finally, the use of a graphics tablet is also recommended. A major disadvantage currently is that the Weka algorithms are extremely CPU intensive but in ImageJ, do not utilise the GPU. K-means and fuzzy c-means algorithms are also extremely CPU intensive, regardless of if they are written in ImageJ or Matlab. Three-dimensional local c-means clustering, as implemented in the MIA package, is very RAM intensive (personal observation). Implementation of the Weka algorithm, either through virtualised clusters (e.g. FIJI archipelago) or through GPU optimisation (either through CLIJ (Haase et al., 2019) or Matlab bridging) may work to ameliorate bottlenecks in processing speed. The yield in minimising user time in segmentation (as once trained, segmentation can process independently of the user) does however make the current implementation an ideal approach for the first pass segmentation of structures in archaeological and evolutionary studies.

Two-dimensional localised c-means segmentation in both ImageJ and Matlab also tend to flip the order of labels in some images, which then necessitates further steps of interleaving different stacks to obtain one segmentation. This is probably straightforward to address by a forcing of order of labels in the algorithm but is beyond the scope of this paper. Our suggestion therefore is if k-means or c-means clustering is used for segmentation, a three-dimensional approach should be taken.

## Conclusions

Here, we have presented the implementation of the Weka Machine learning library to archaeological and palaeontological material and compared it with conventional thresholding and other well-known machine-learning clustering techniques. For many of the examples presented, it yields results that are the equal of leading edge-based methods and superior to conventional threshold-based segmentations. It is also able to cope well with the introduction of imaging artefacts (whether real or simulated). For absolute accuracy with high contrast scans where only one material is of interest, a fully automated approach using three-dimensional local c-means (Dunmore et al., 2018) performs the best. For the simultaneous segmentation of multiple materials of equal interest, (especially where scans are sub-optimal in quality) we recommend using Weka based segmentation, as less manual intervention to correct the segmentation is required afterwards (though three dimensional local c-means does segment multiple materials simultaneously).

The implementation of Weka segmentation is fast, with no software cost to the end user and it enables an easy introduction to both image segmentation and machine learning for the inexperienced user. Future work will seek to apply this algorithm to larger and more varied samples and increasing the speed of computation, either through GPU based acceleration or use of virtual clusters.

## Supporting information

S1. Raw data and segmentations of synthetic dataset

S2. Weka_segmentation_simple_guide

## Acknowledgements

We would like to thank the curators of the collections from which scan material was used for this paper, including Campell Price and Sam Sportun (Manchester Museum-acces to mummifieid samples); Dr.V.Khartanovich from Kunstkamera, Russian Academy of Sciences (access to Kiik Koba 2) Professor N.Potrakhov (Electrotechnical University, St-Petersbourg) for assistance scanning Kiik Koba 2. The Duke Lemur Center provided access to the *Otolemur garnetti* data, originally appearing in Yapuncich et al. (2019), the collection of which was funded by NSF BCS 1540421 to Gabriel S. Yapuncich and Doug M. Boyer. The files were downloaded from www.MorphoSource.org, Duke University. The animal mummy was scanned at the Henry Moseley X-ray Imaging Facility, University of Manchester. We are grateful to our reviewers, Dr Elsa Panciroli and Chanin Nantasenamat and an anonymous reviewer for constructive feedback on our originally submitted version and to our academic editor, Dr Walter de Azevedo Jr. for his careful handling of our manuscript. William Harcourt-Smith made the very helpful suggestion of including the anisotropy analyses in our comparisons. We are also grateful to Christopher Dunmore and Charlotte Brassey for detailed feedback on earlier versions of this article. Finally, we are grateful to the School of Life Sciences, Anglia Ruskin University for computing resources.

