## Supplementary figures and images for "A machine learning based approach to the segmentation of micro CT data in archaeological and evolutionary sciences"

### 000.png

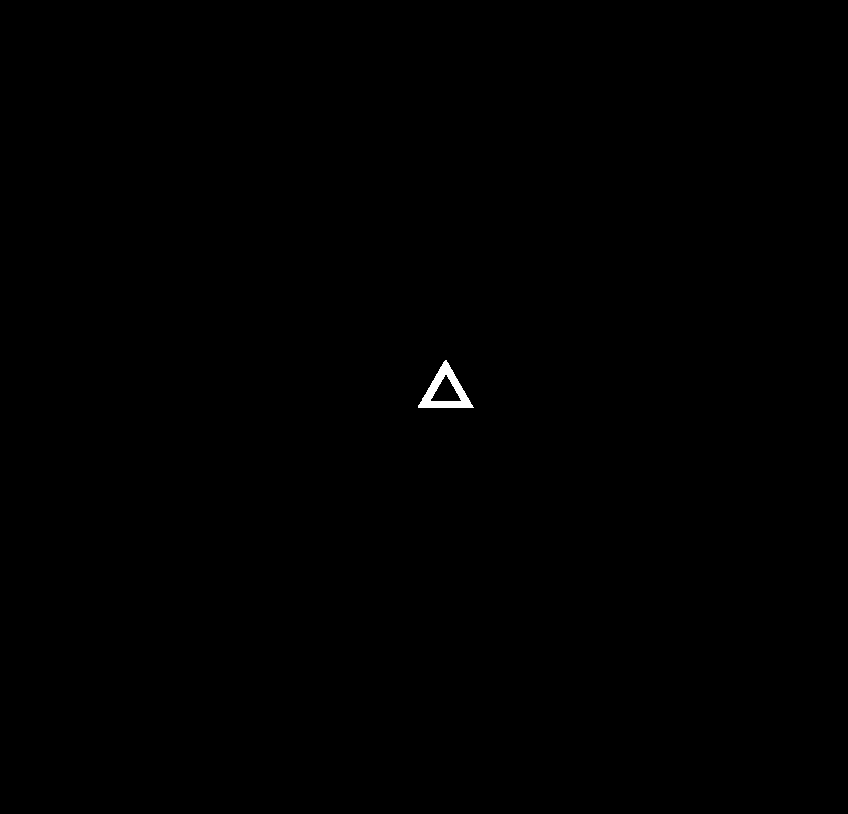

### 001.png

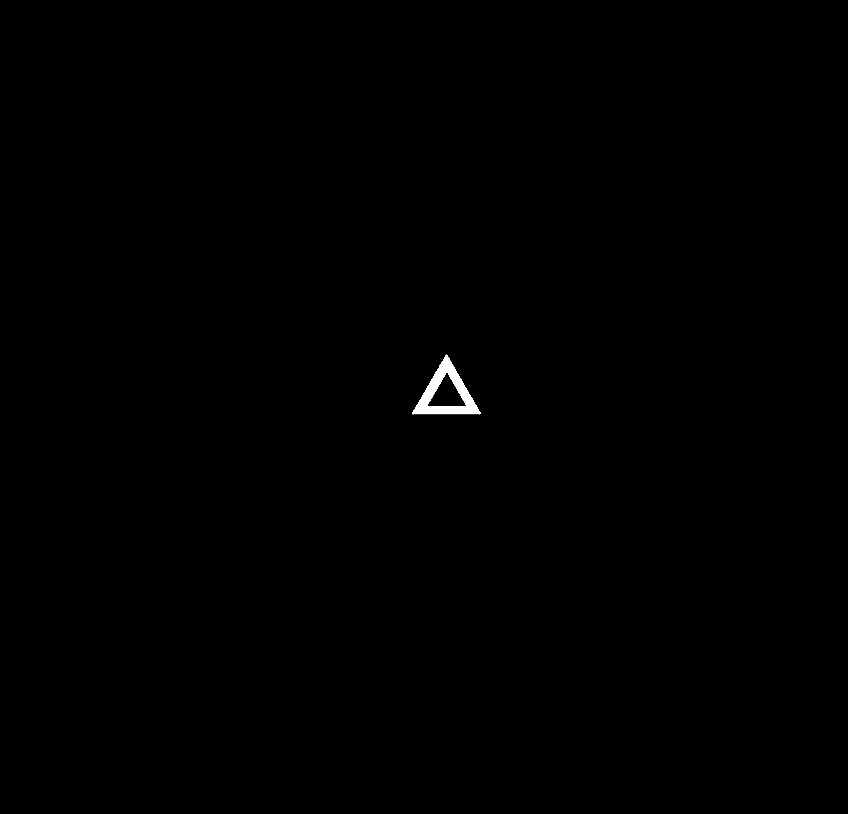

### 002.png

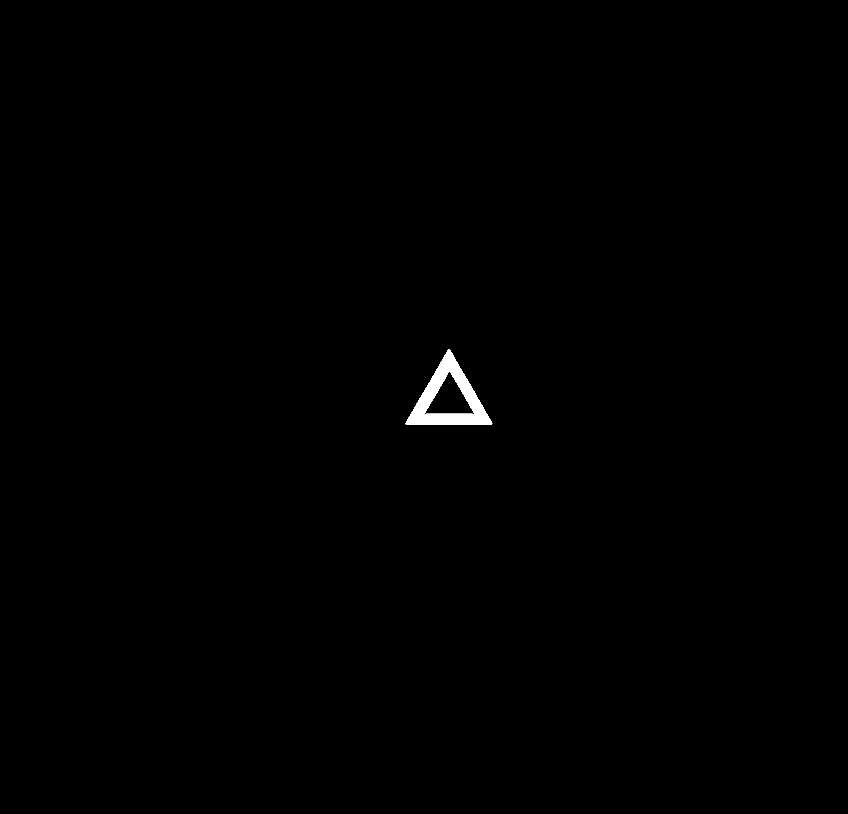

### 003.png

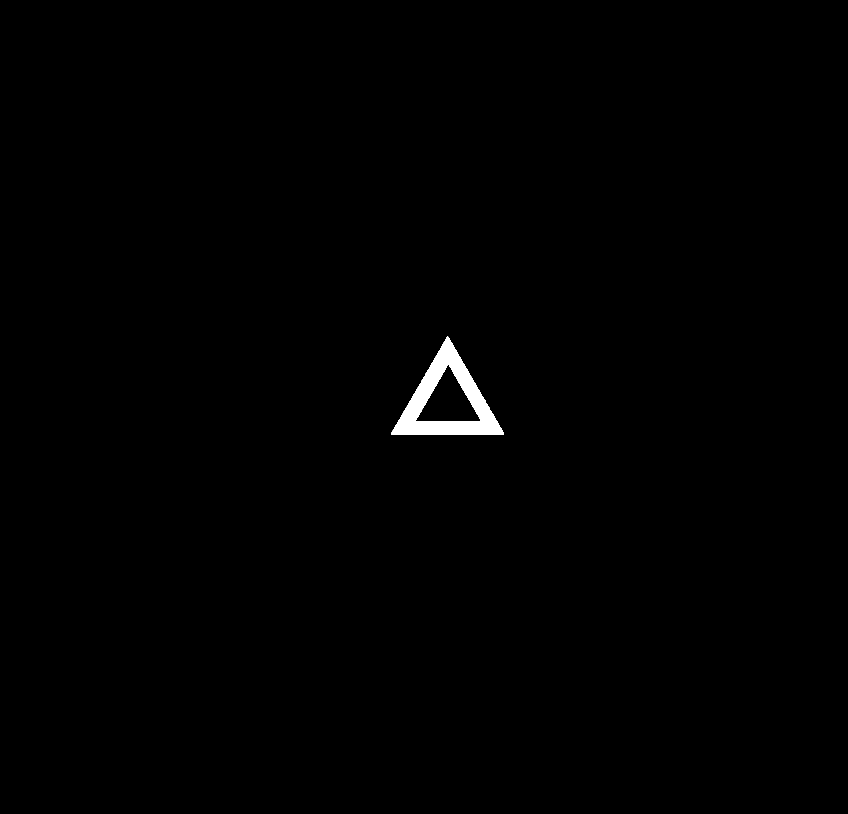

### 004.png

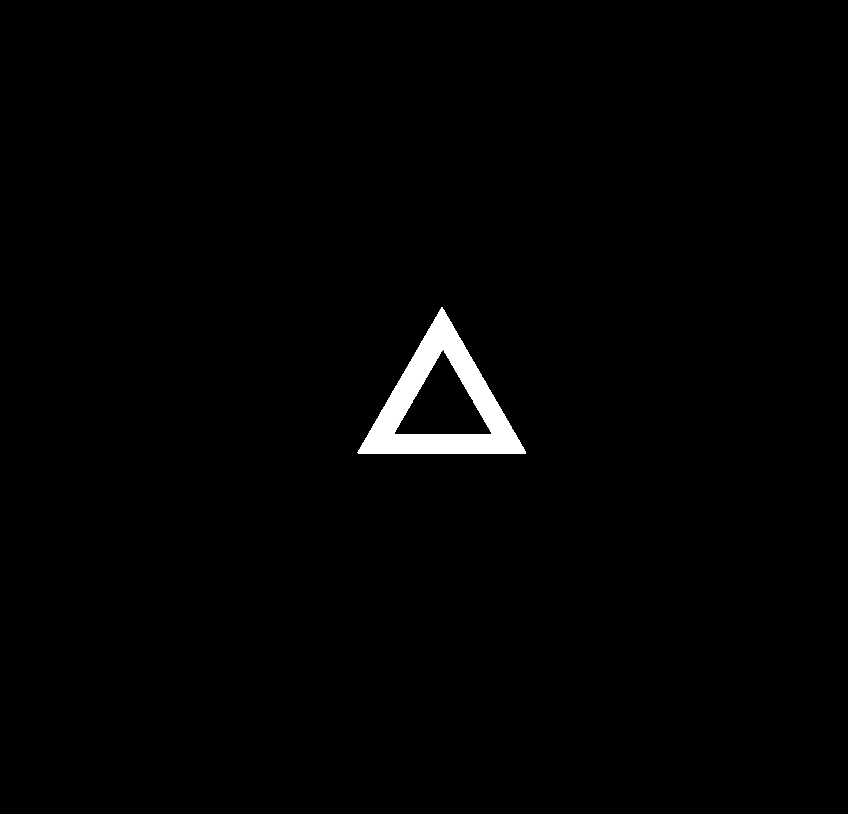

### 005.png

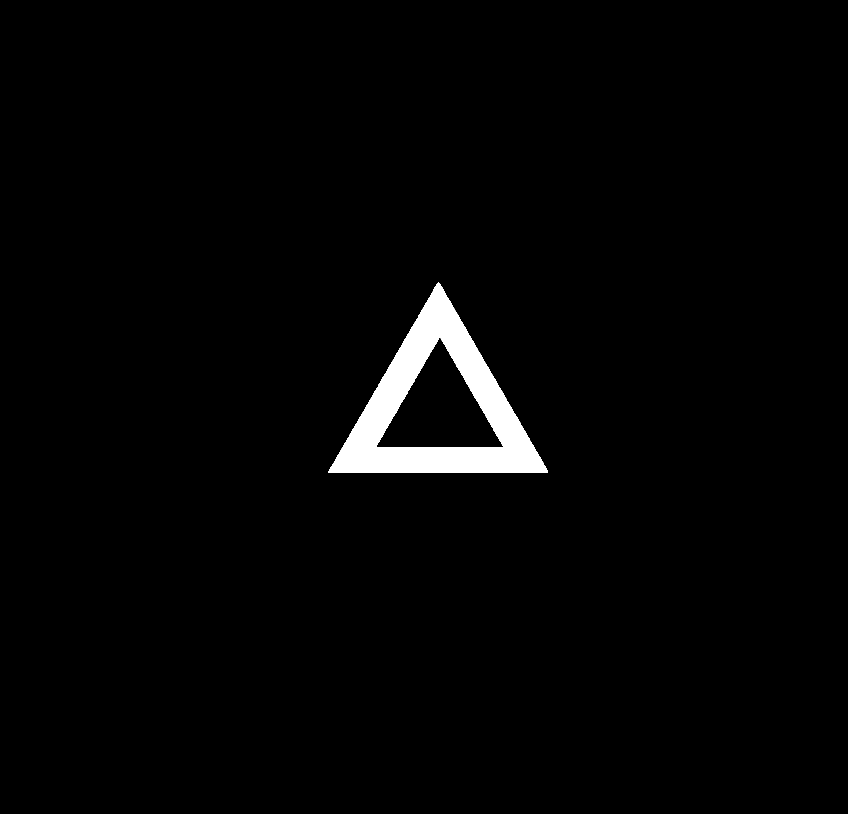

### 006.png

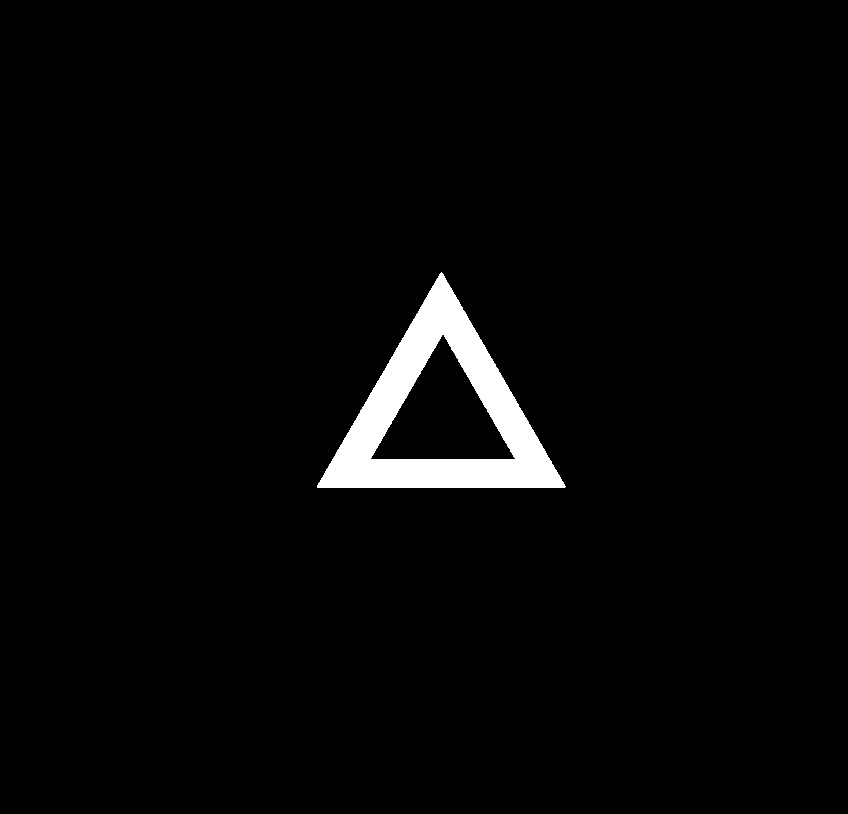

### 007.png

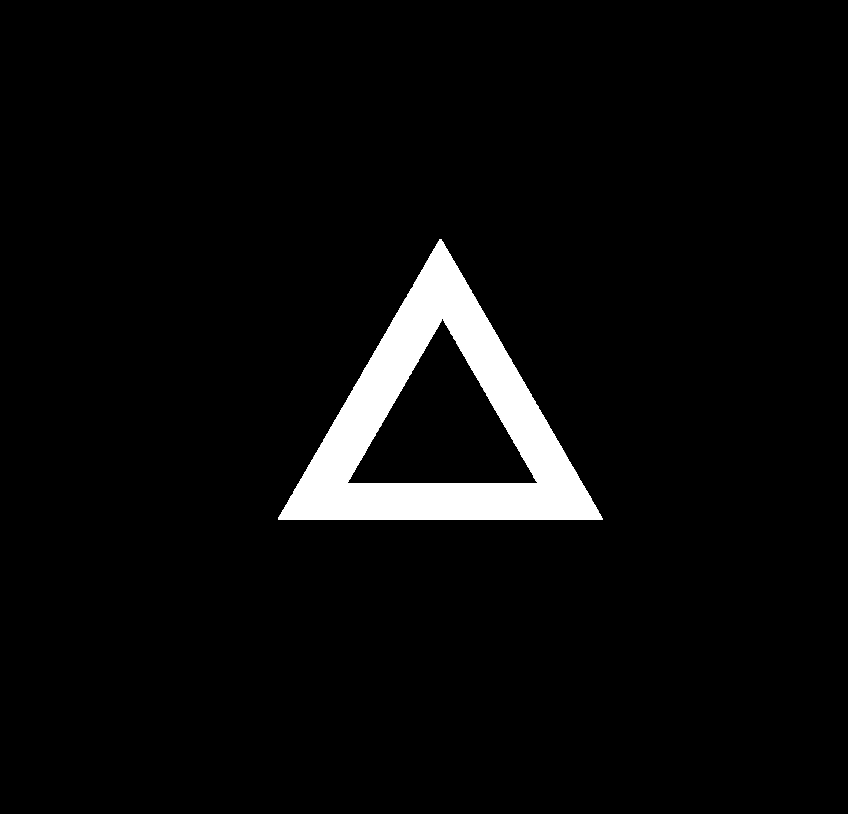

### 008.png

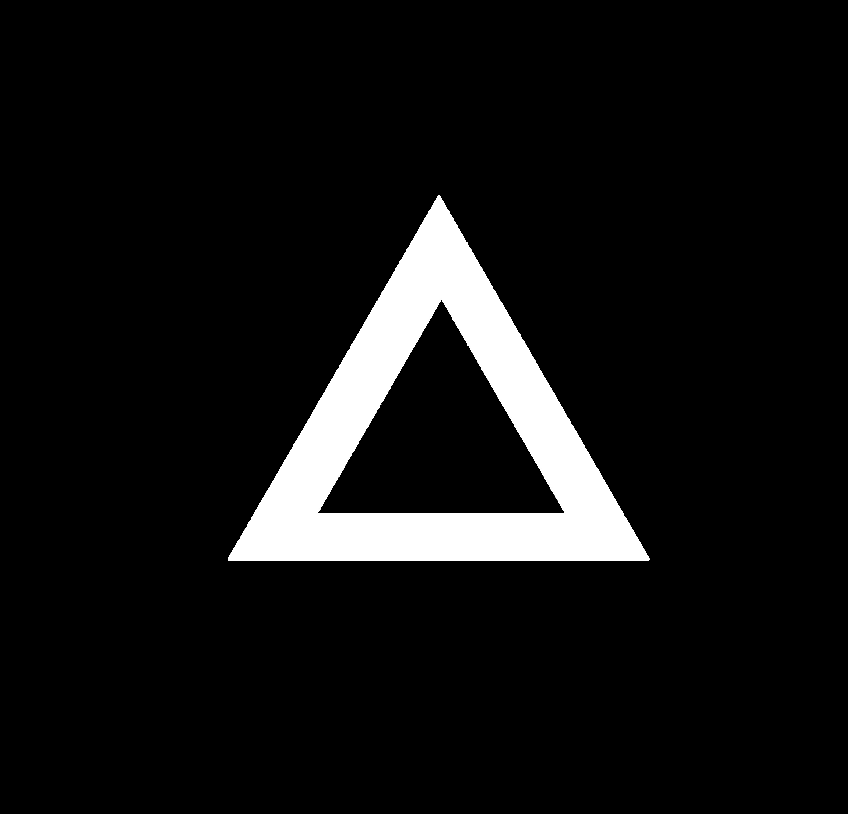

### 009.png

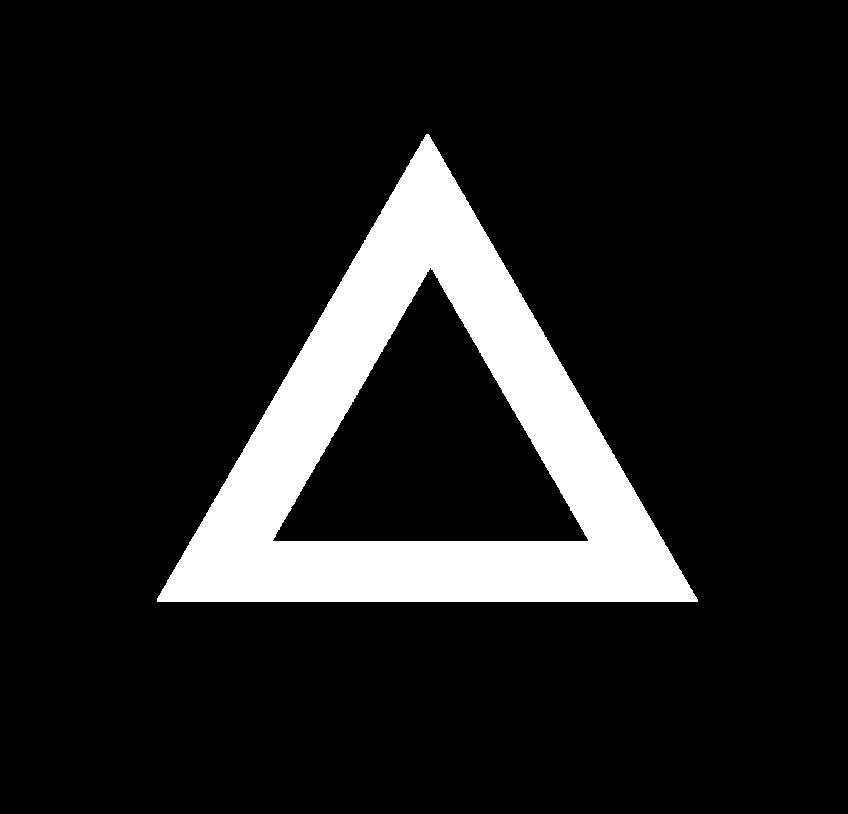

### 010.png

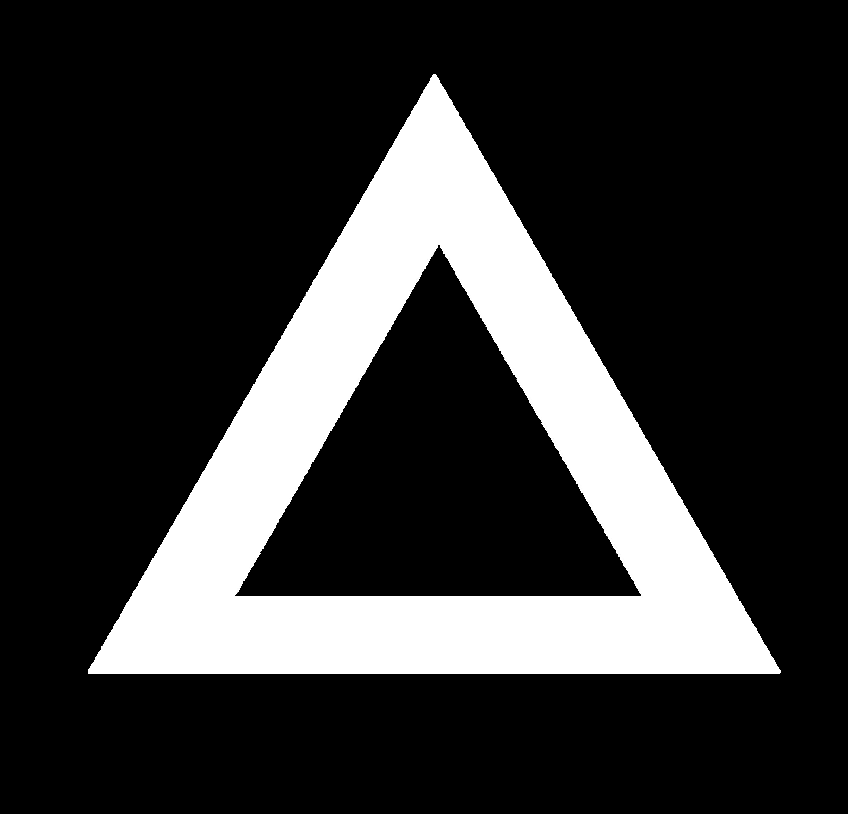

### 011.png

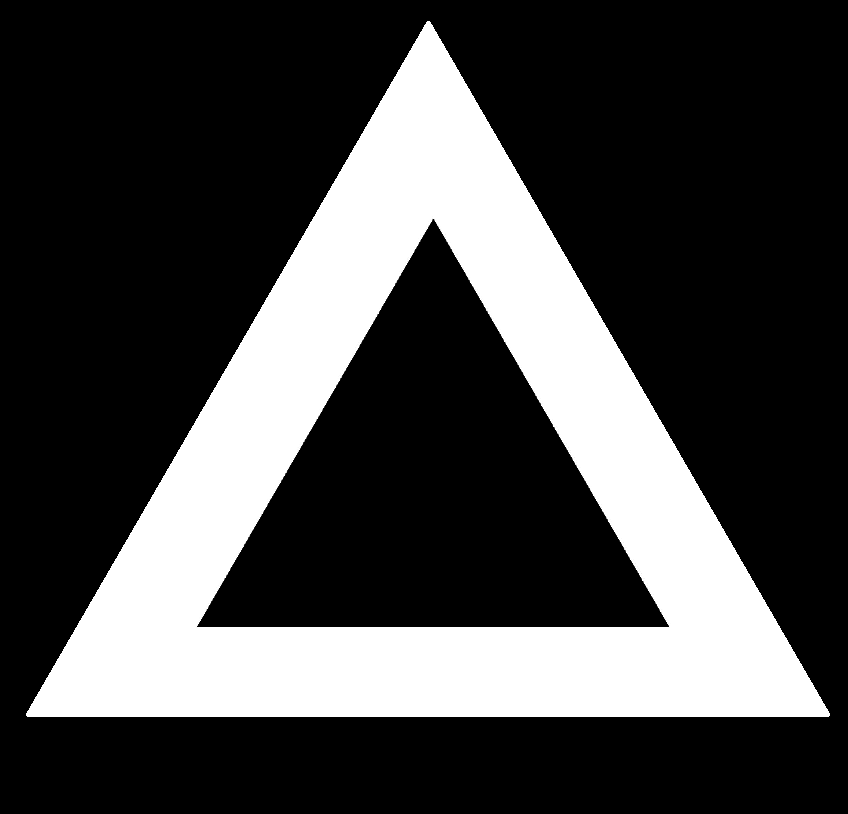

### Clusters0000.tif

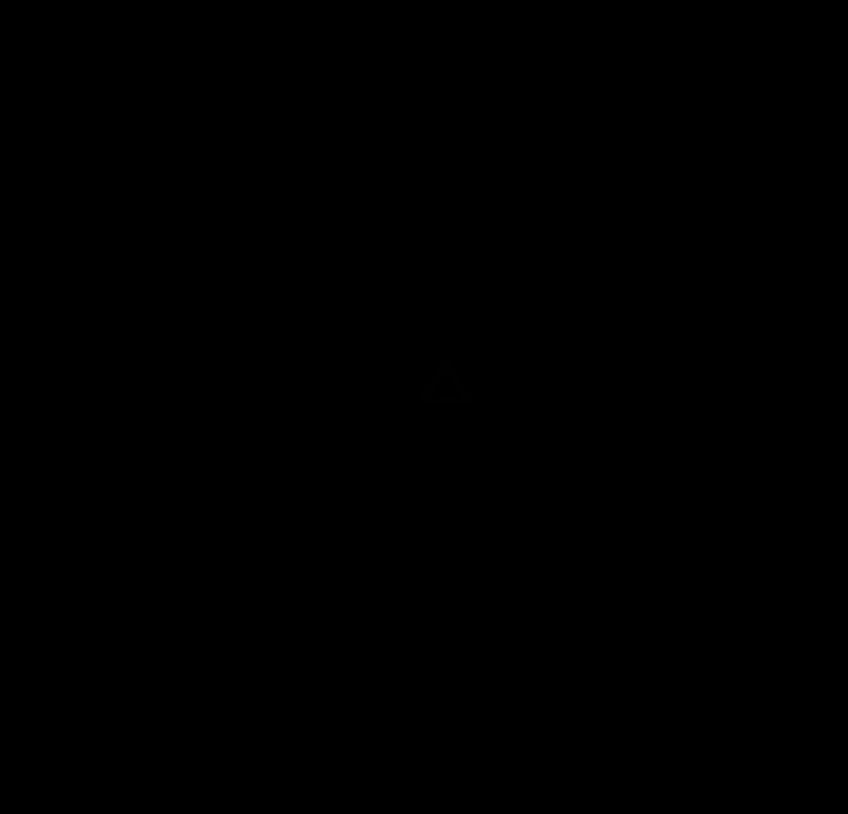

### Clusters0001.tif

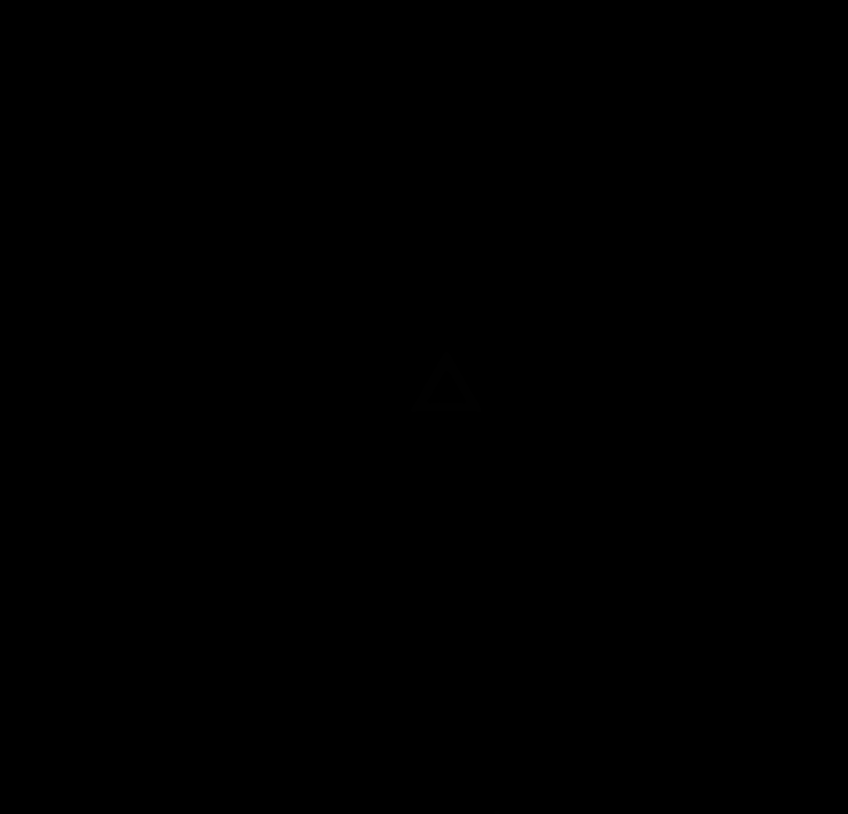

### Clusters0002.tif

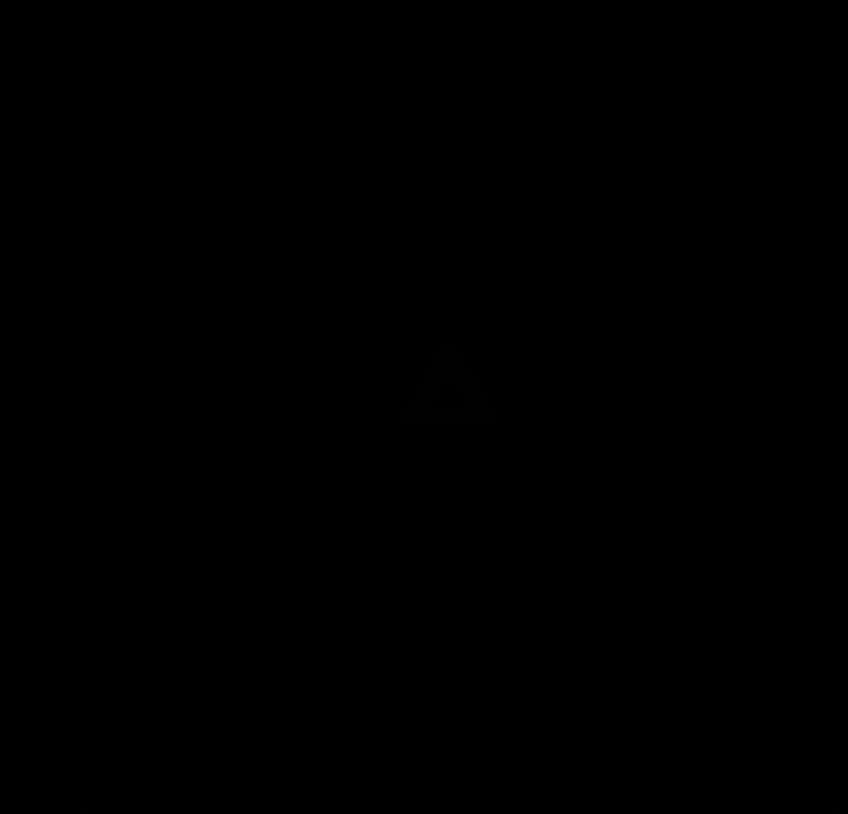

### Clusters0003.tif

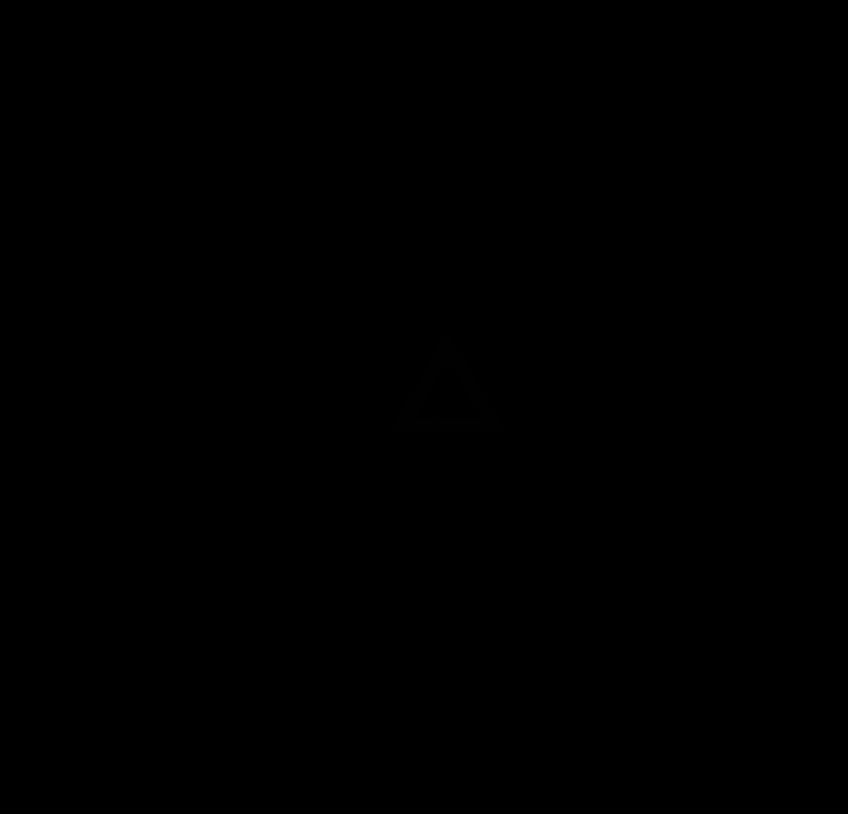

### Clusters0004.tif

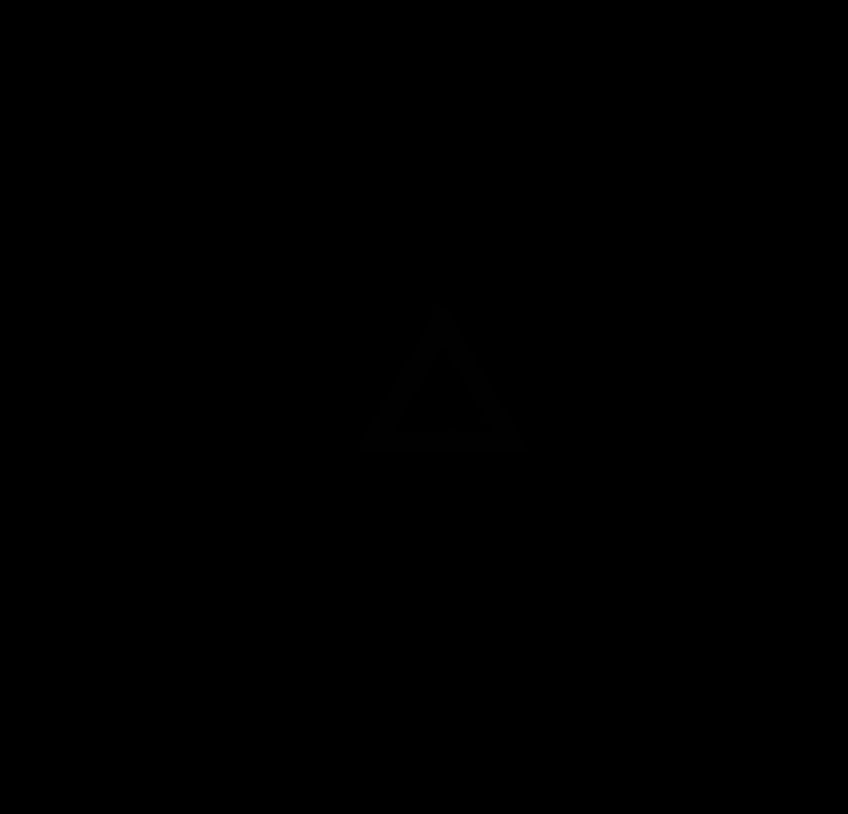

### Clusters0005.tif

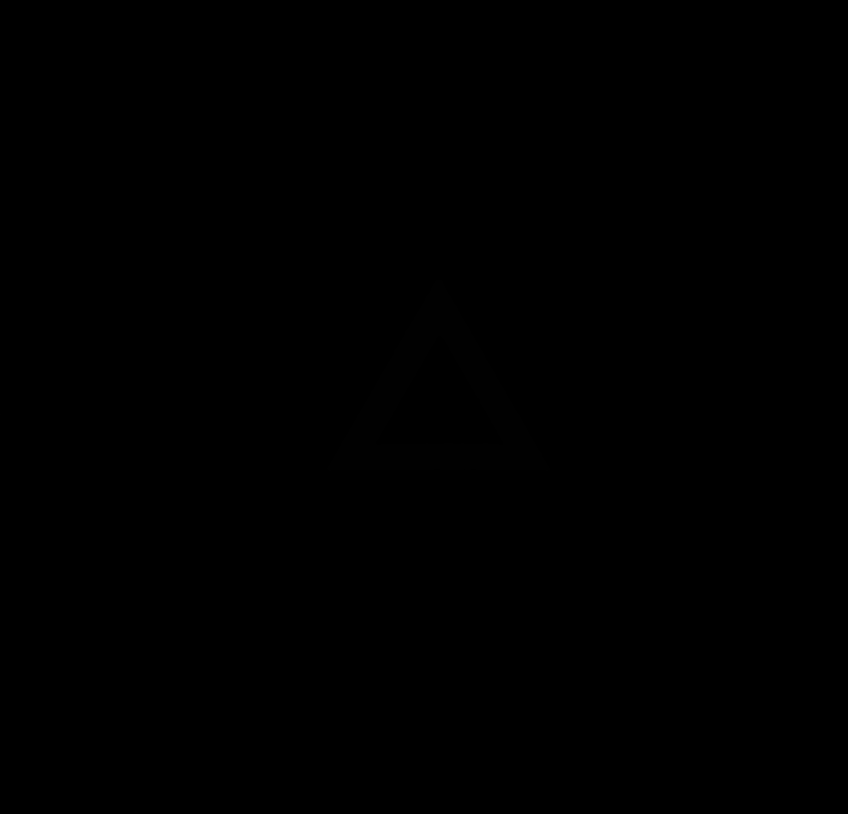

### Clusters0006.tif

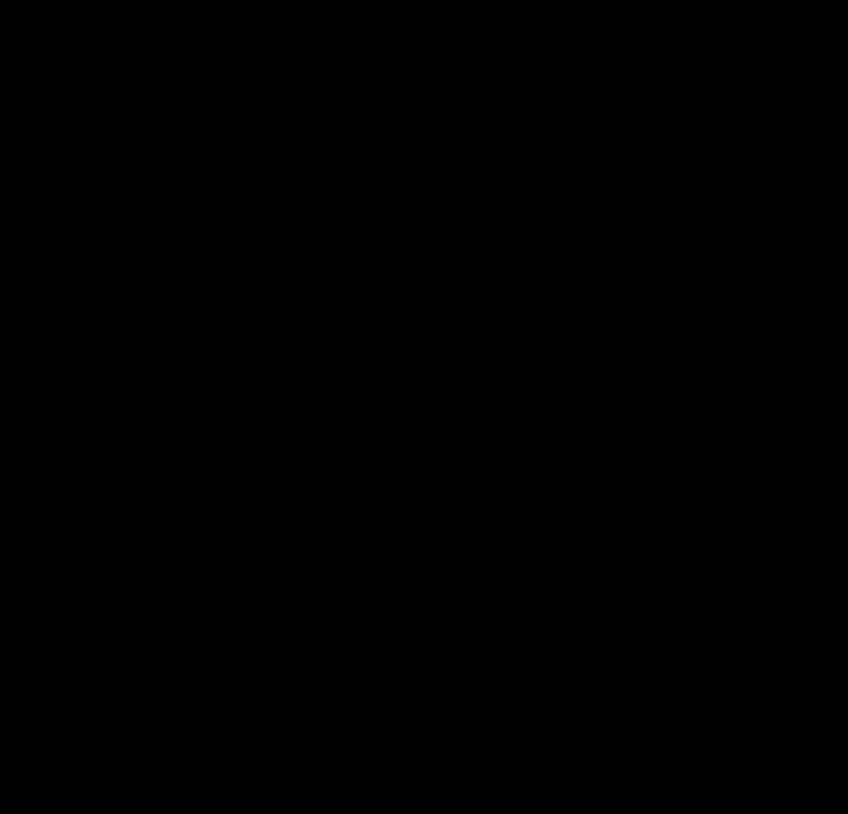

### Clusters0007.tif

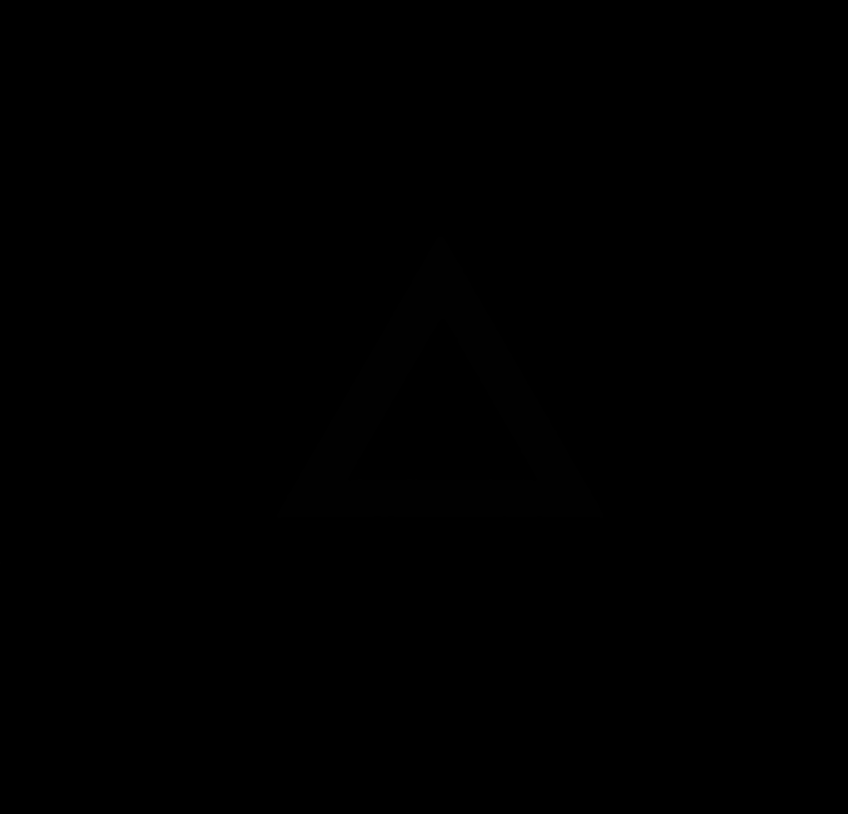

### Clusters0008.tif

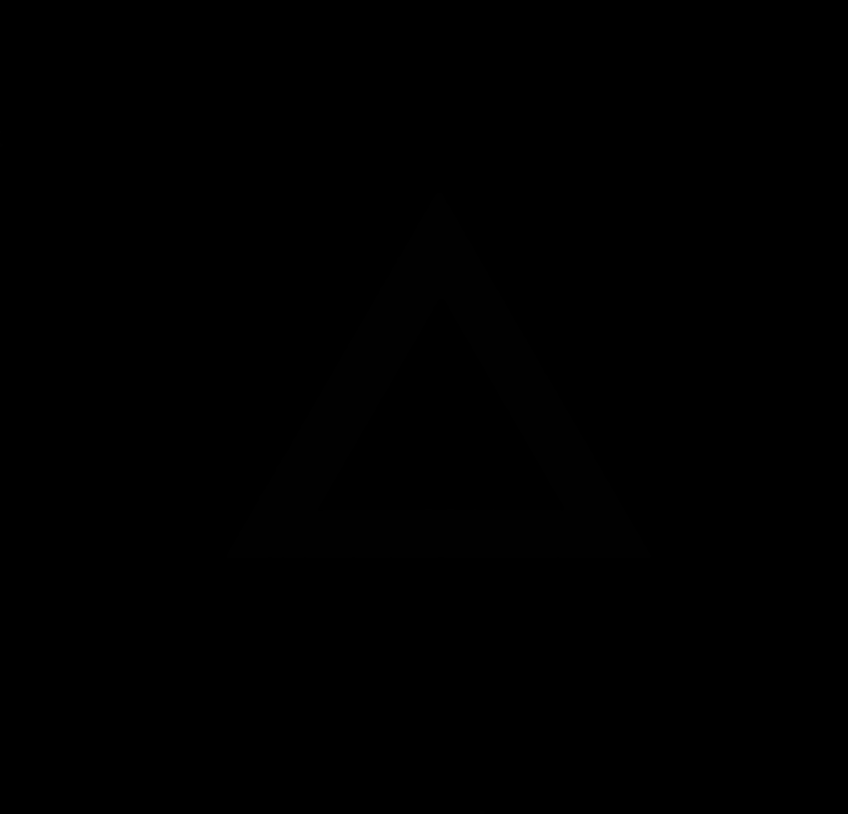

### Clusters0009.tif

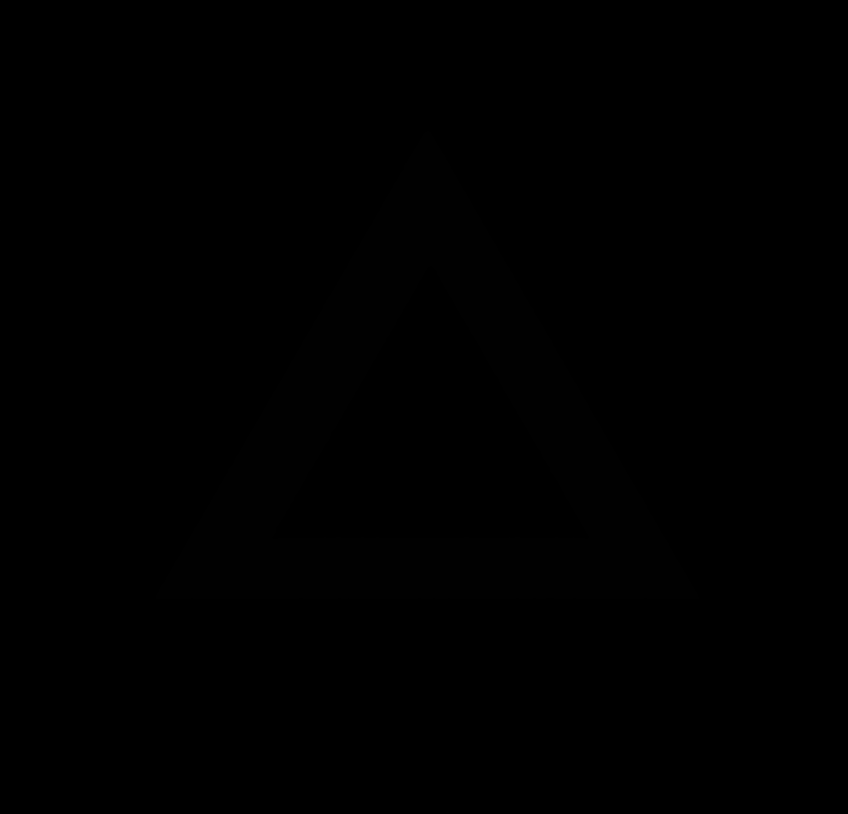

### Clusters0010.tif

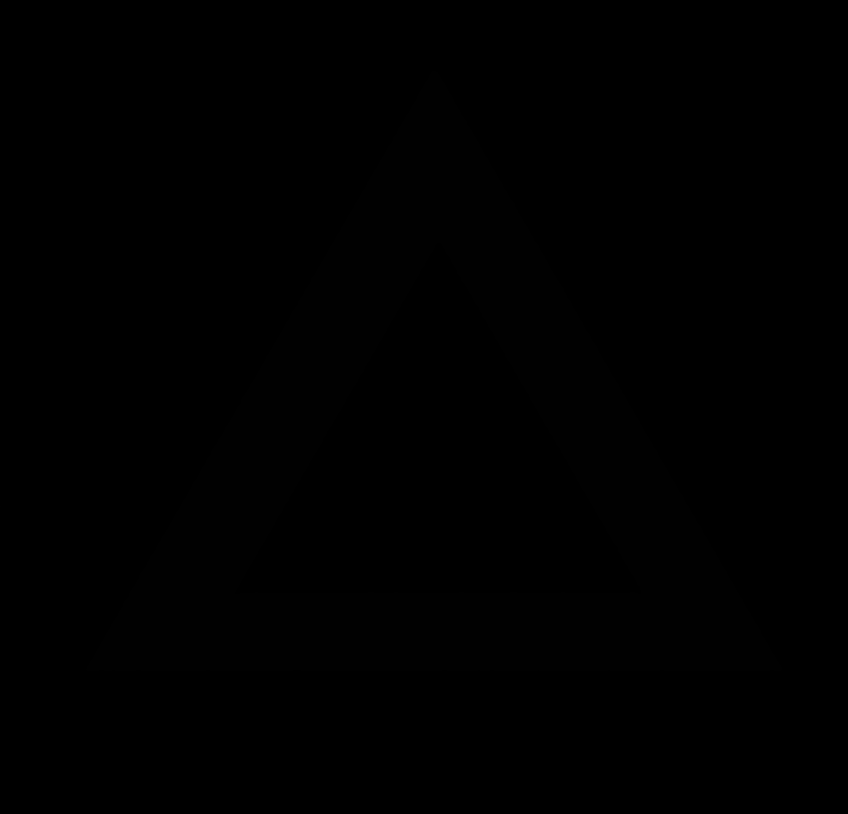

### Clusters0011.tif

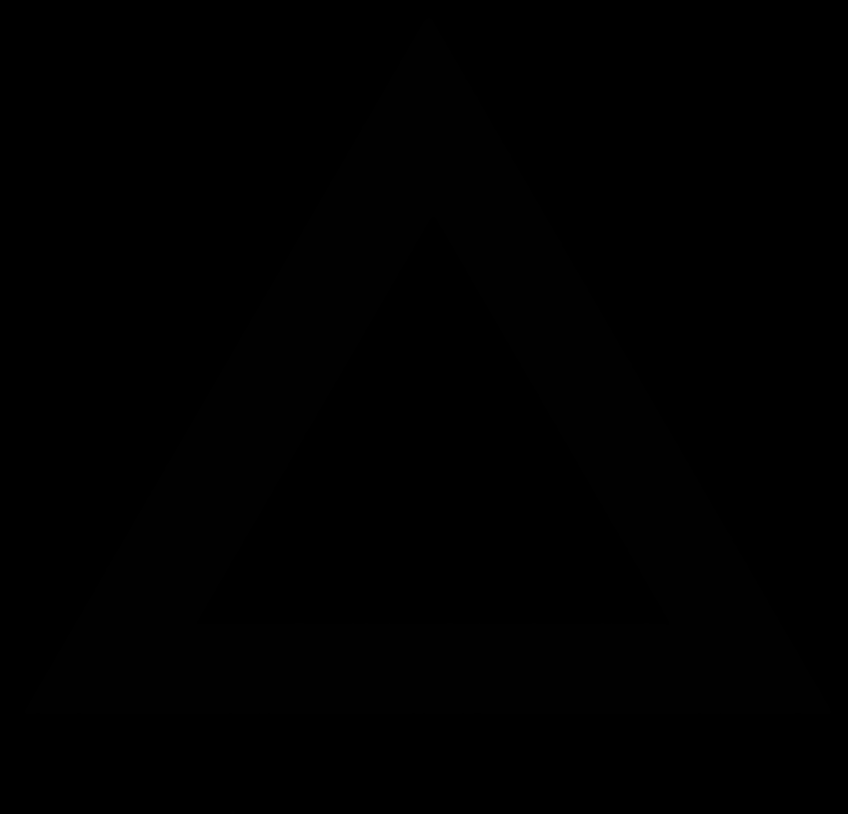

### Original synthetic data00.tif

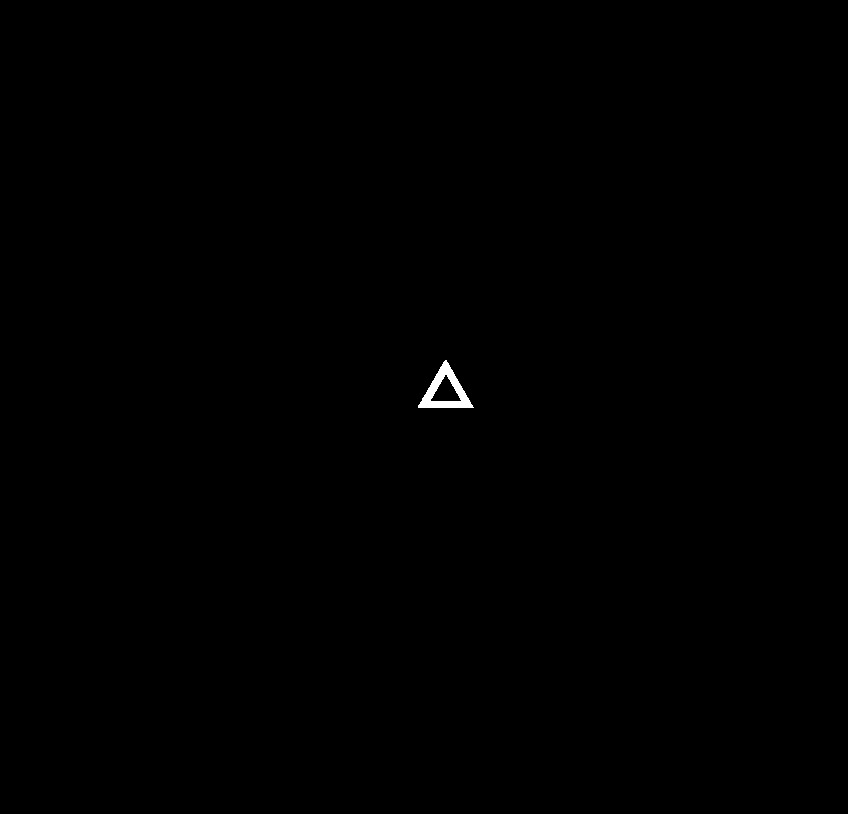

### Original synthetic data01.tif

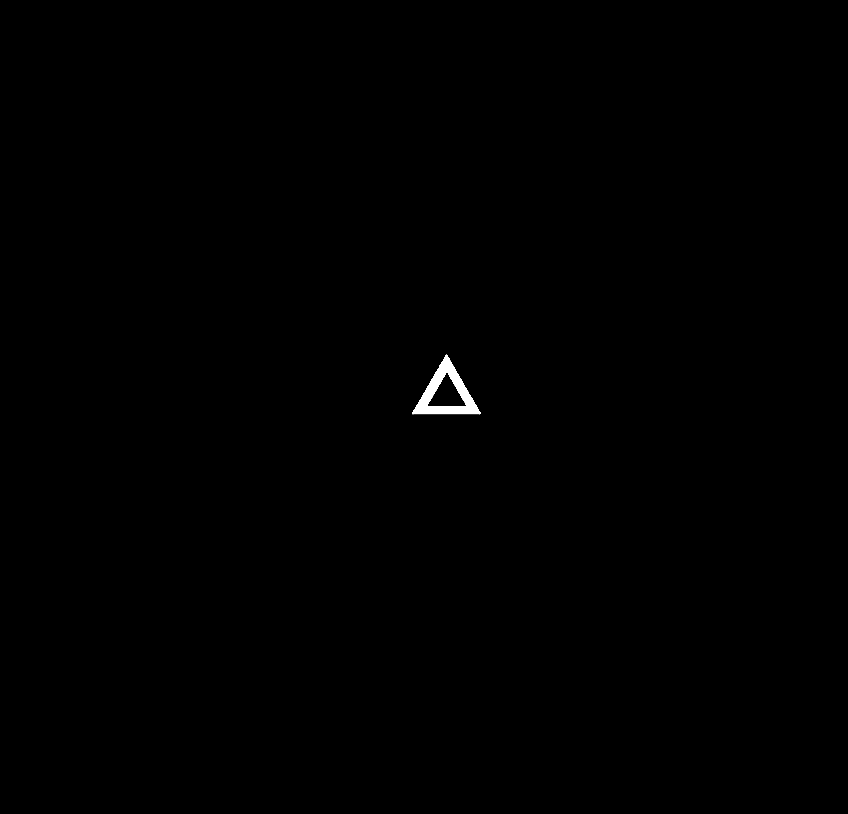

### Original synthetic data02.tif

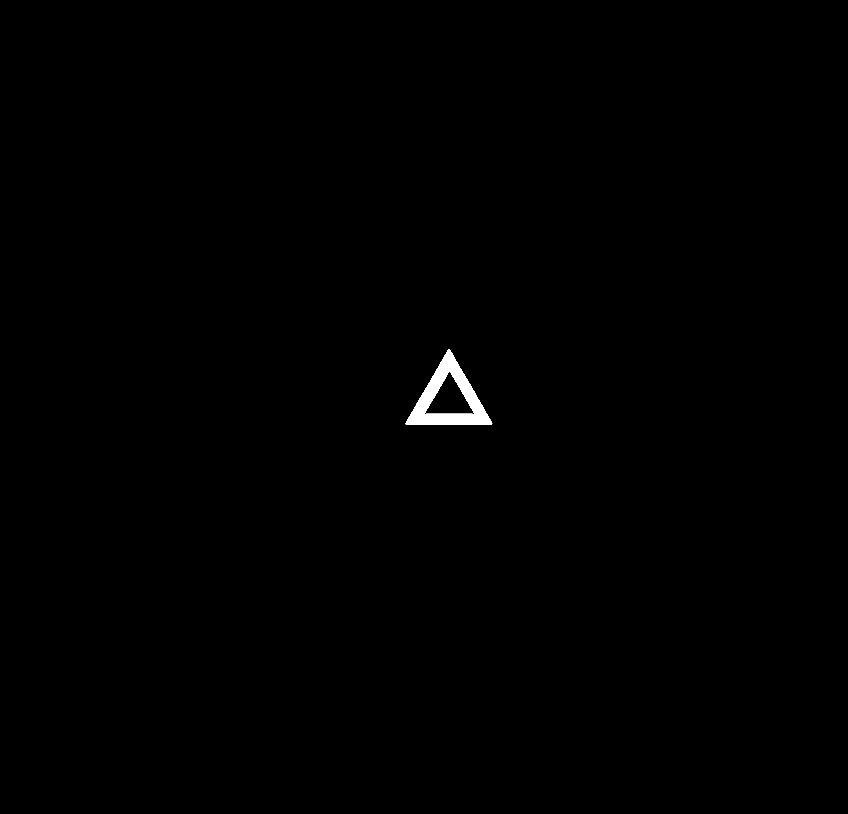

### Original synthetic data03.tif

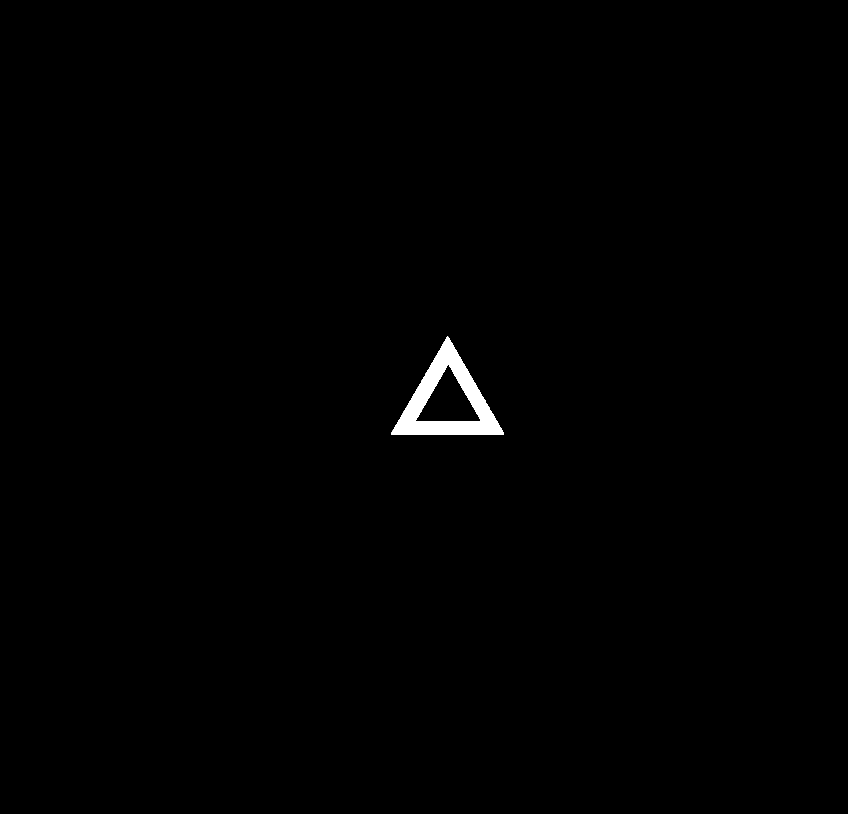

### Original synthetic data04.tif

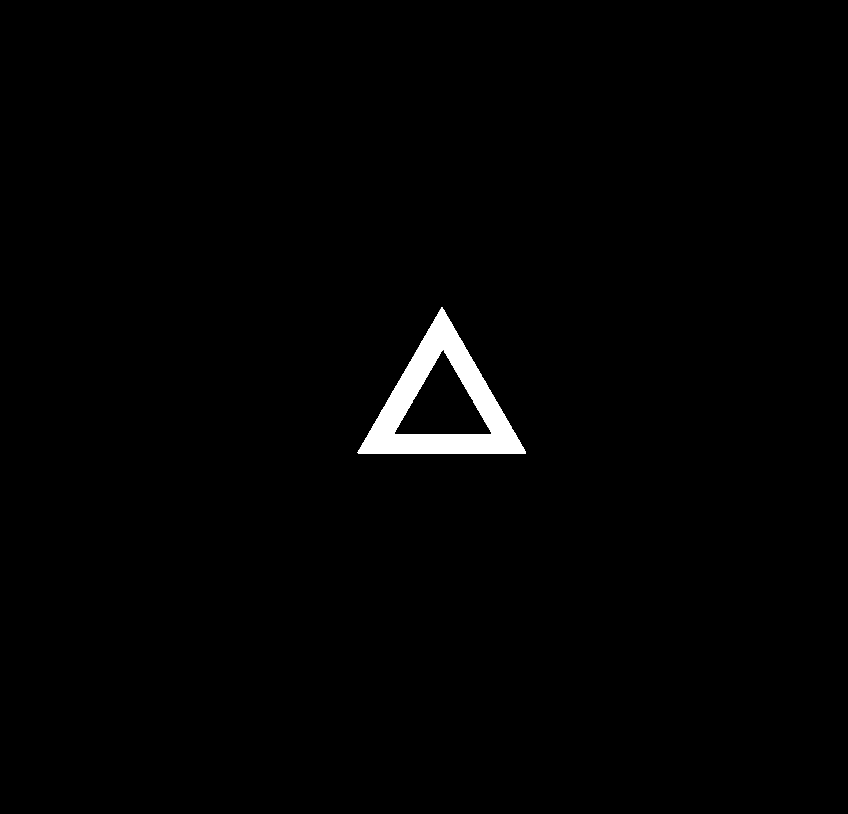

### Original synthetic data05.tif

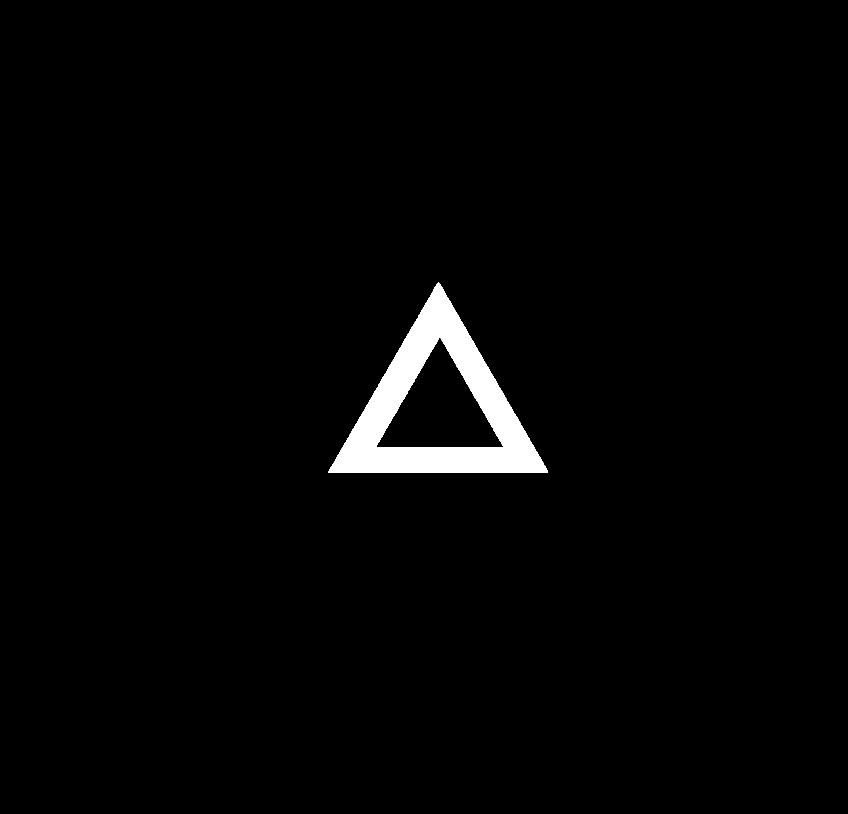

### Original synthetic data06.tif

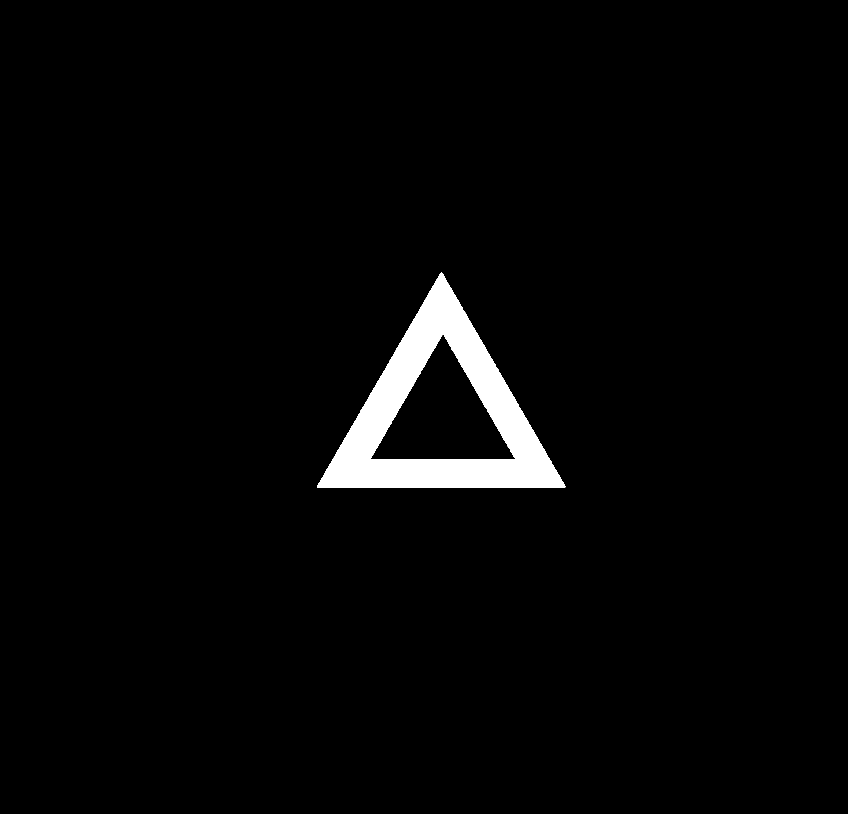

### Original synthetic data07.tif

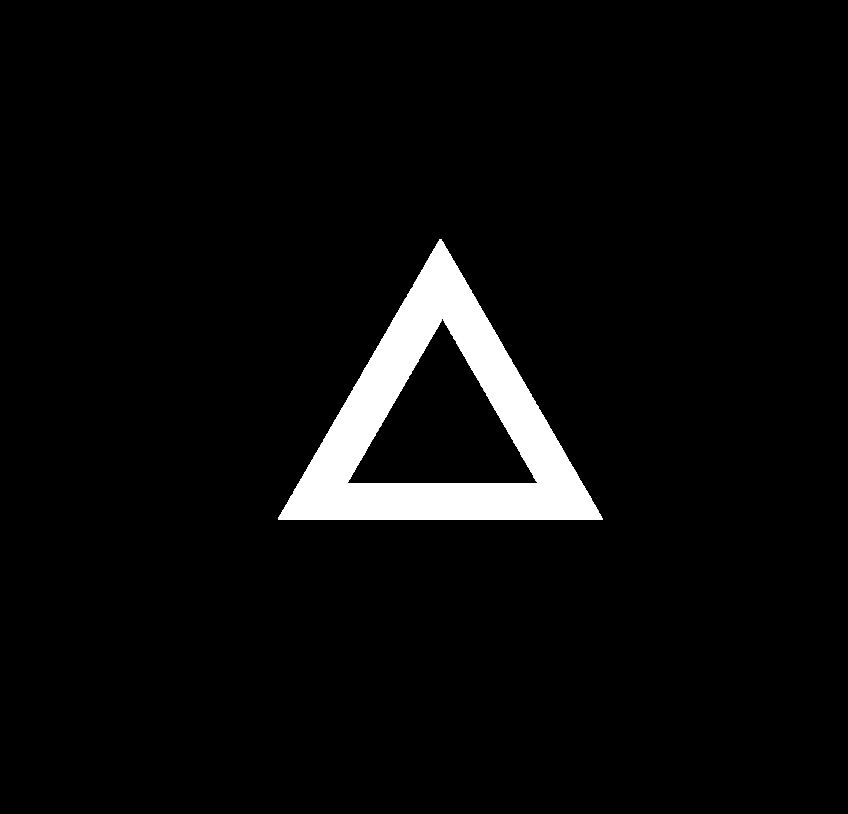

### Original synthetic data08.tif

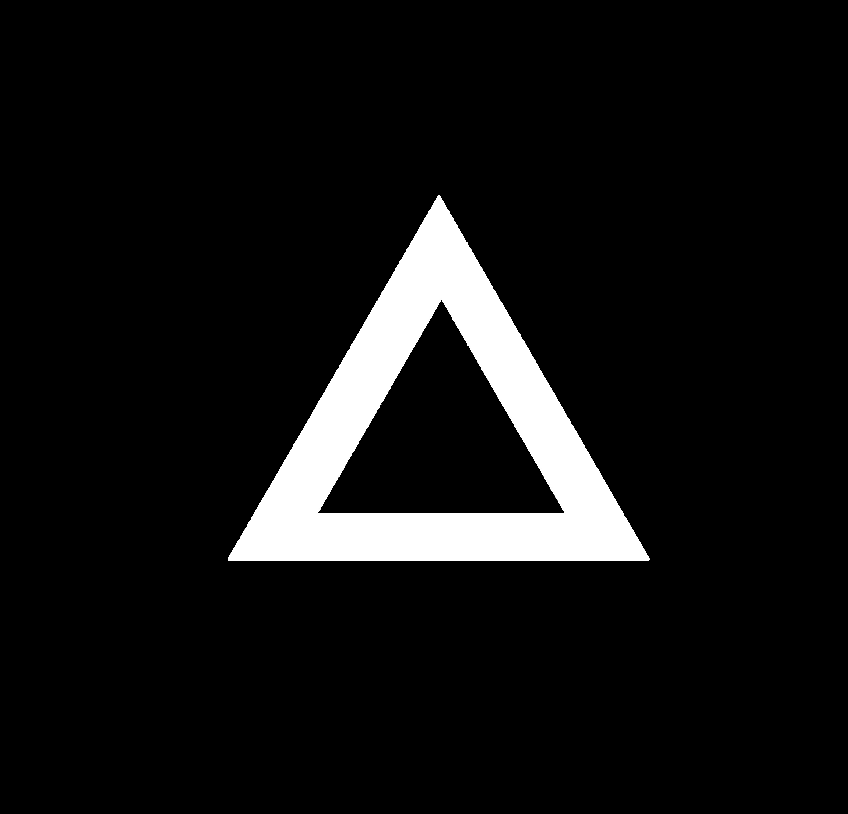

### Original synthetic data09.tif

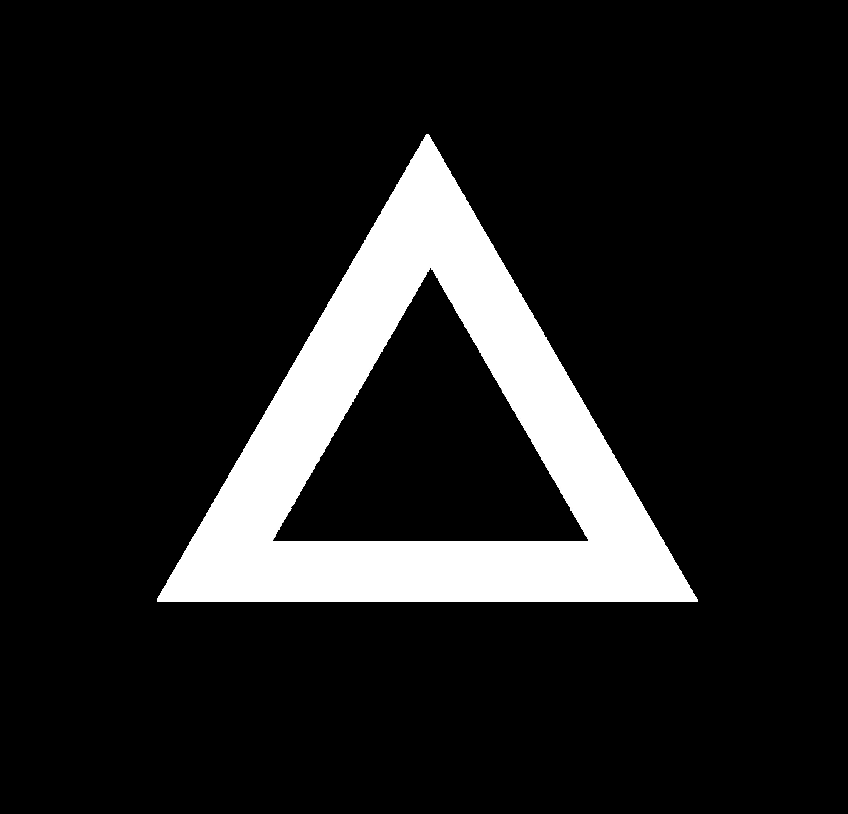

### Original synthetic data10.tif

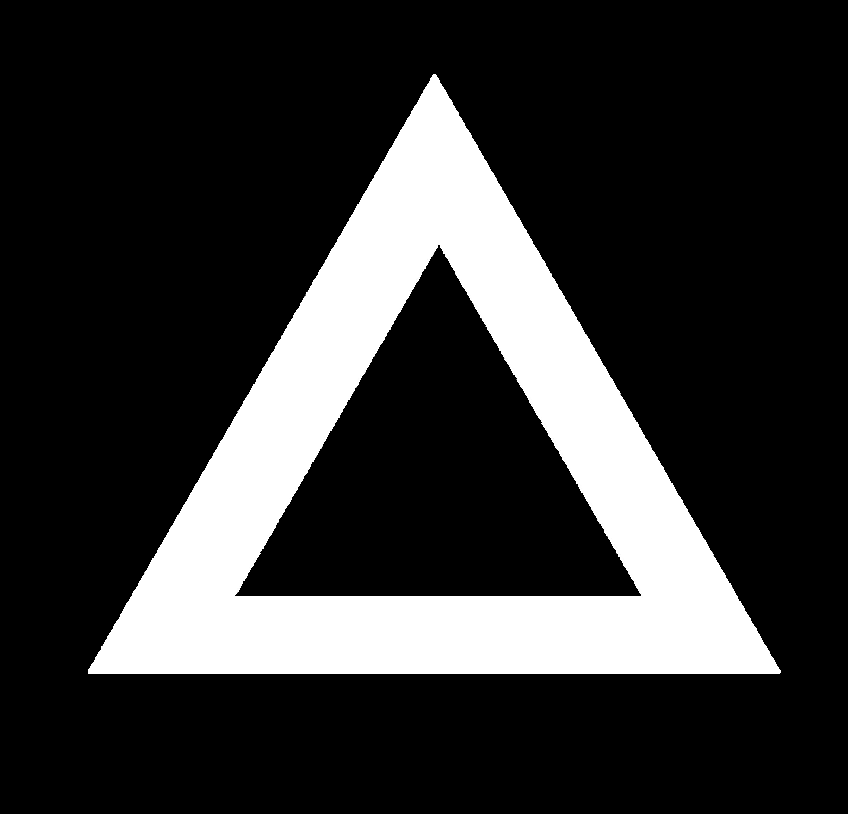

### Original synthetic data11.tif

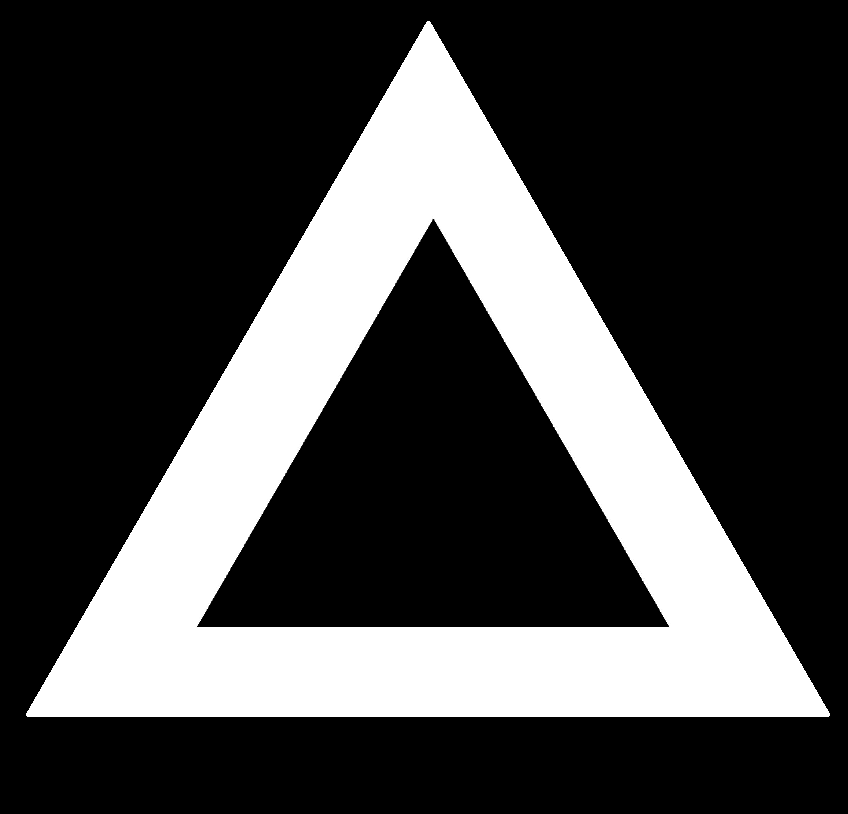

### Original synthetic data-10000.tif

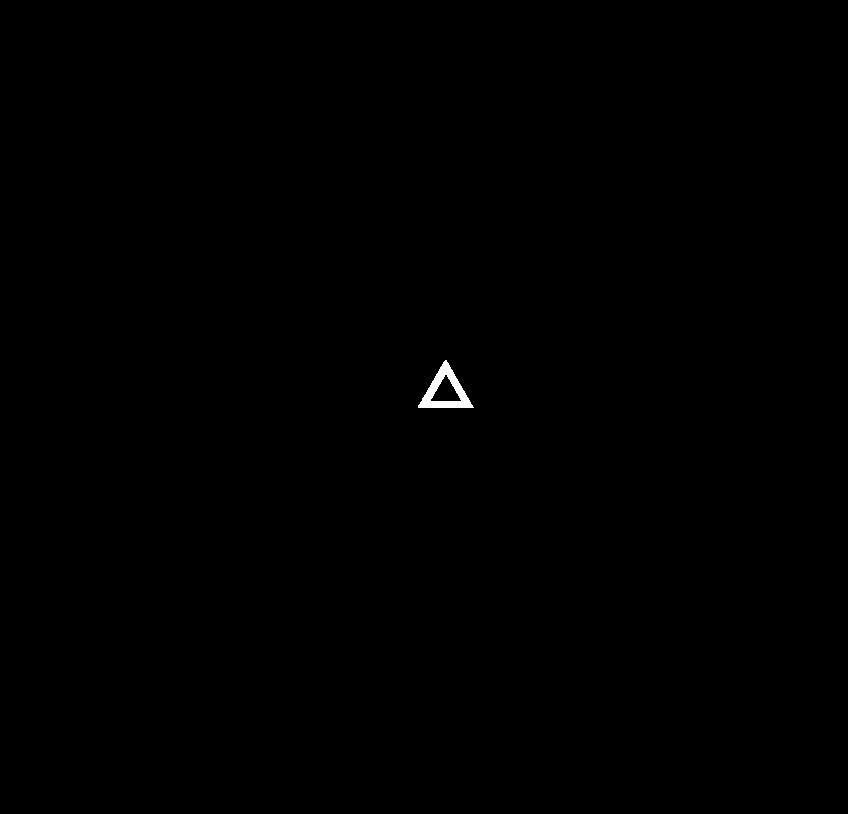

### Original synthetic data-10001.tif

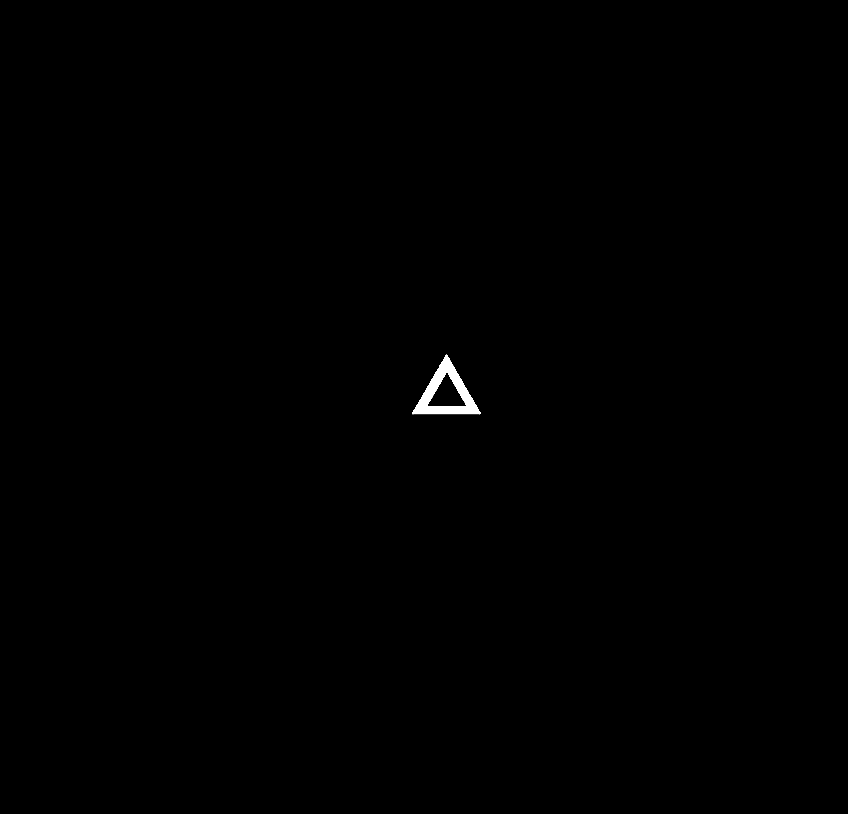

### Original synthetic data-10002.tif

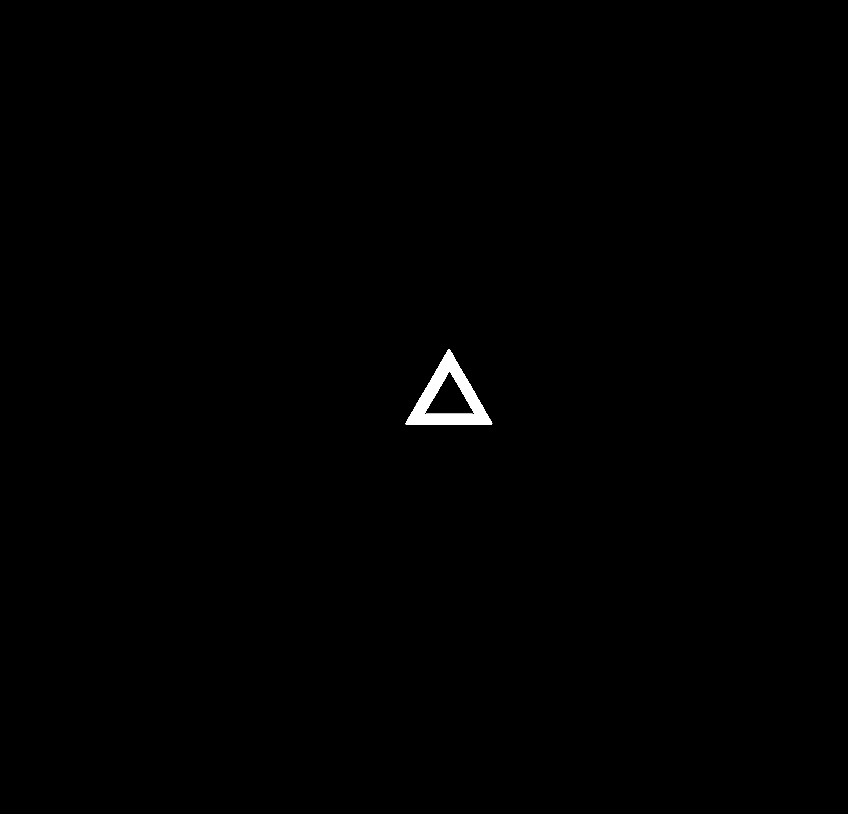

### Original synthetic data-10003.tif

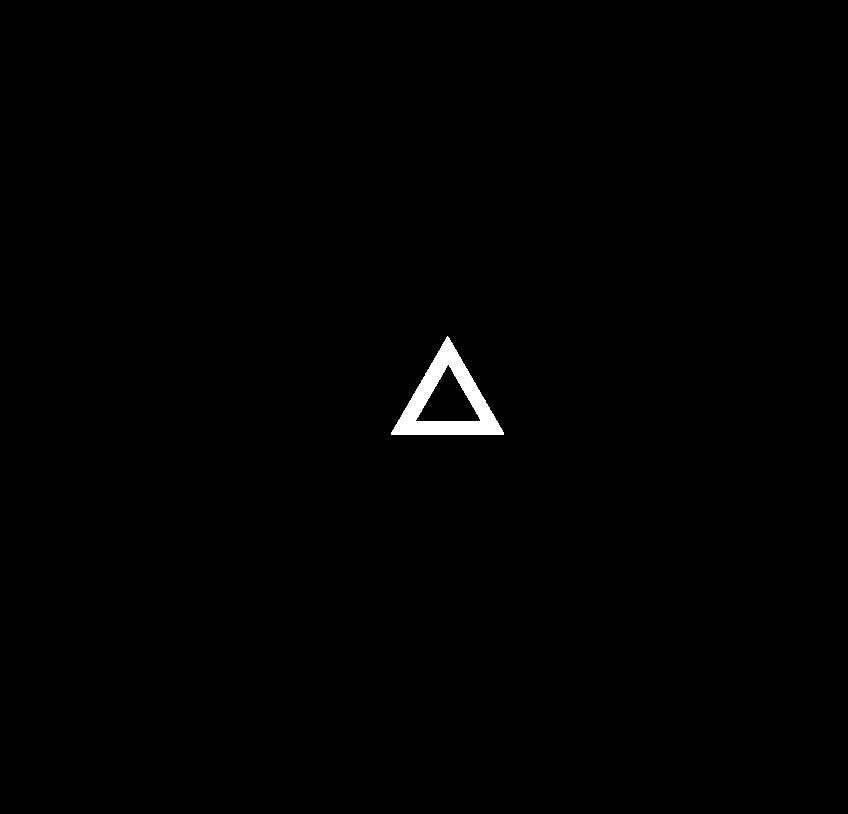

### Original synthetic data-10004.tif

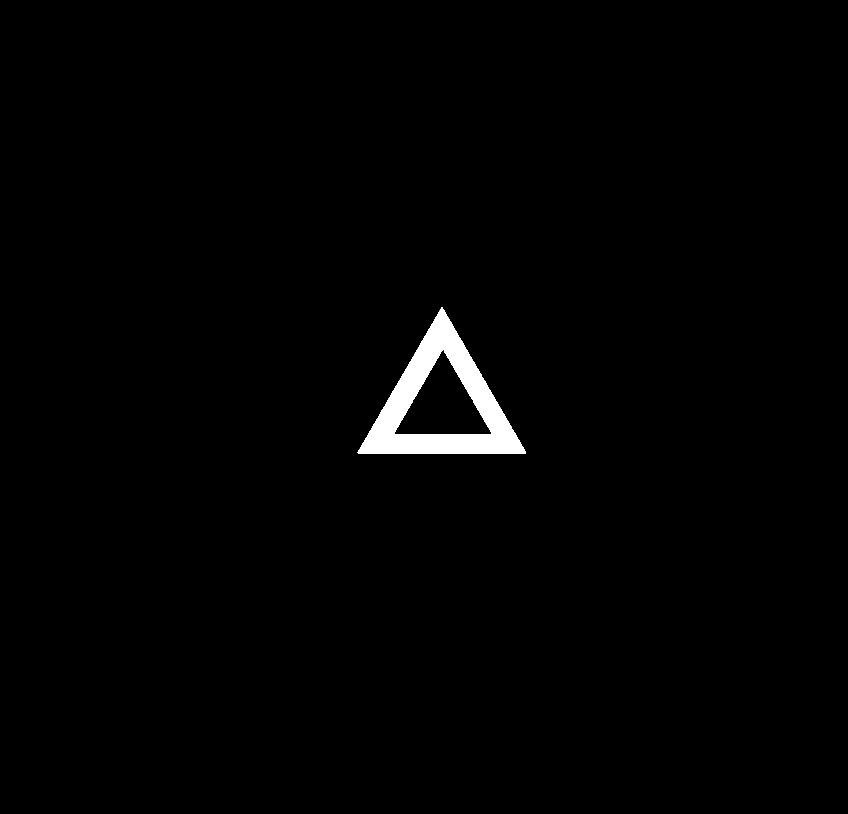

### Original synthetic data-10005.tif

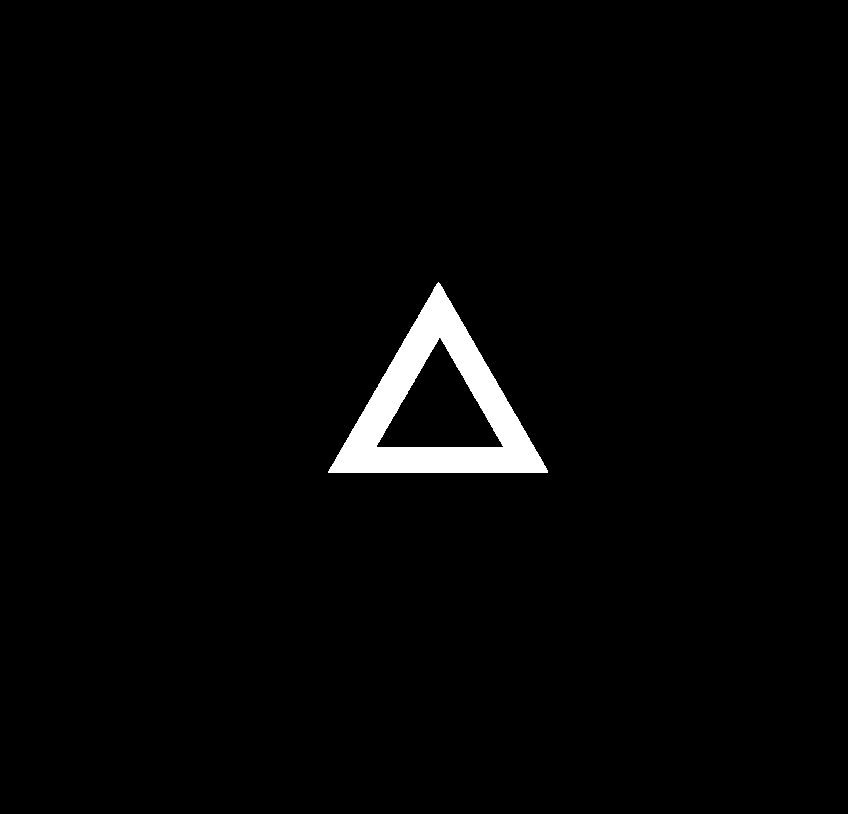

### Original synthetic data-10006.tif

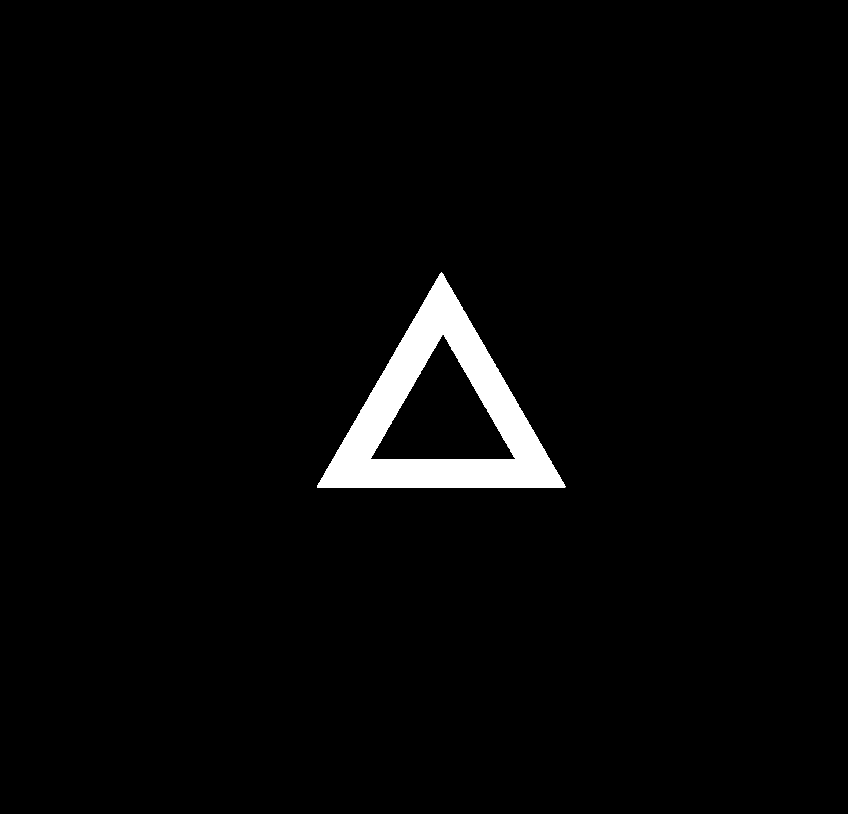

### Original synthetic data-10007.tif

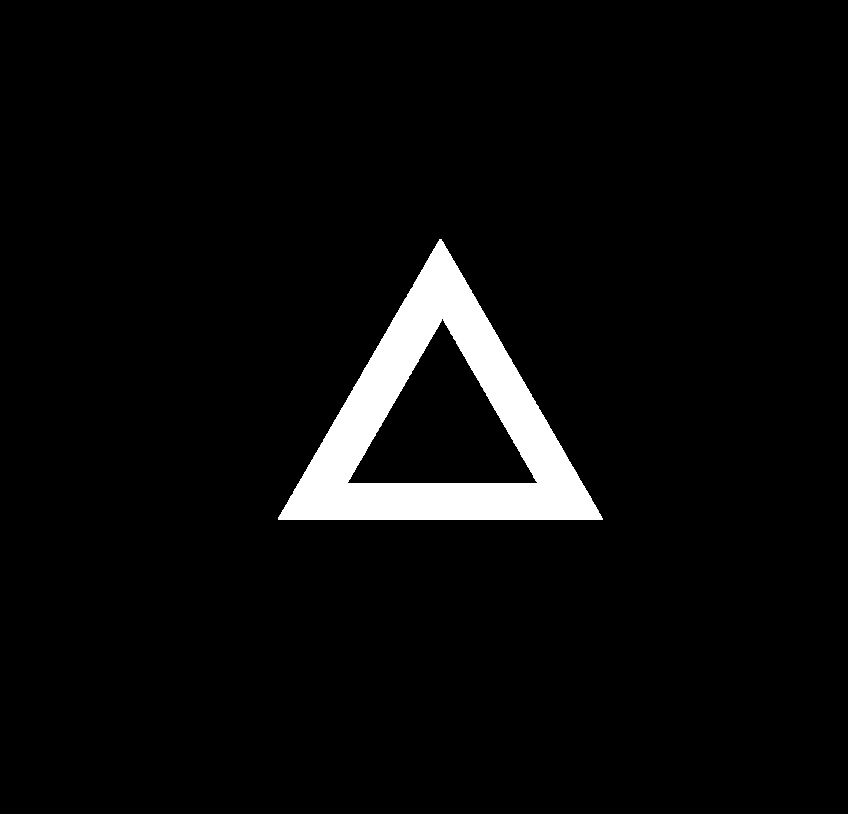

### Original synthetic data-10008.tif

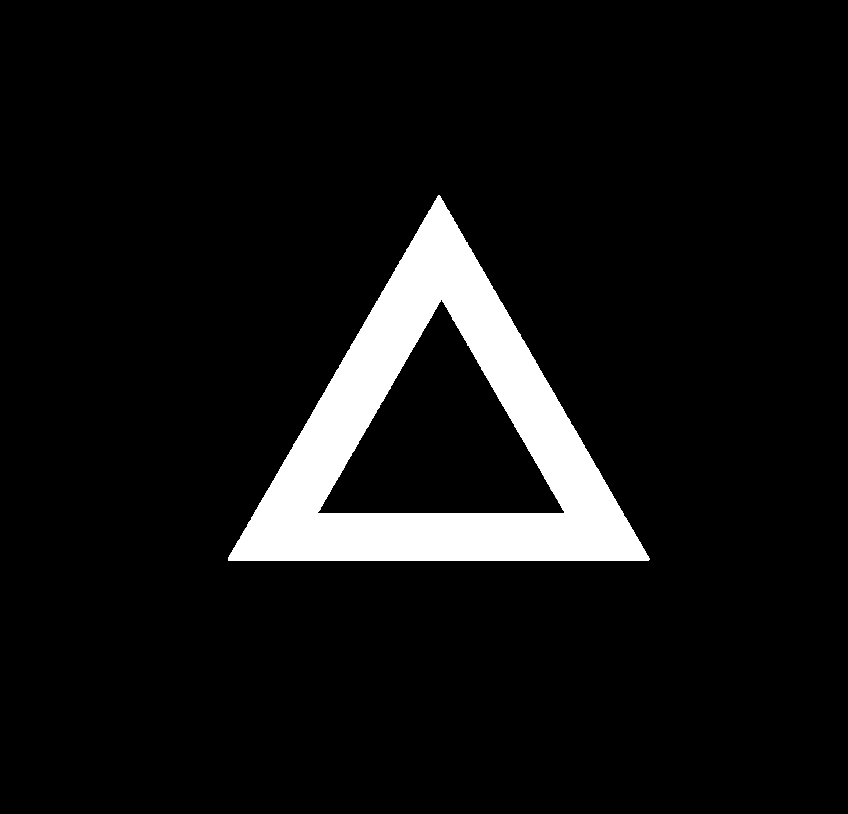

### Original synthetic data-10009.tif

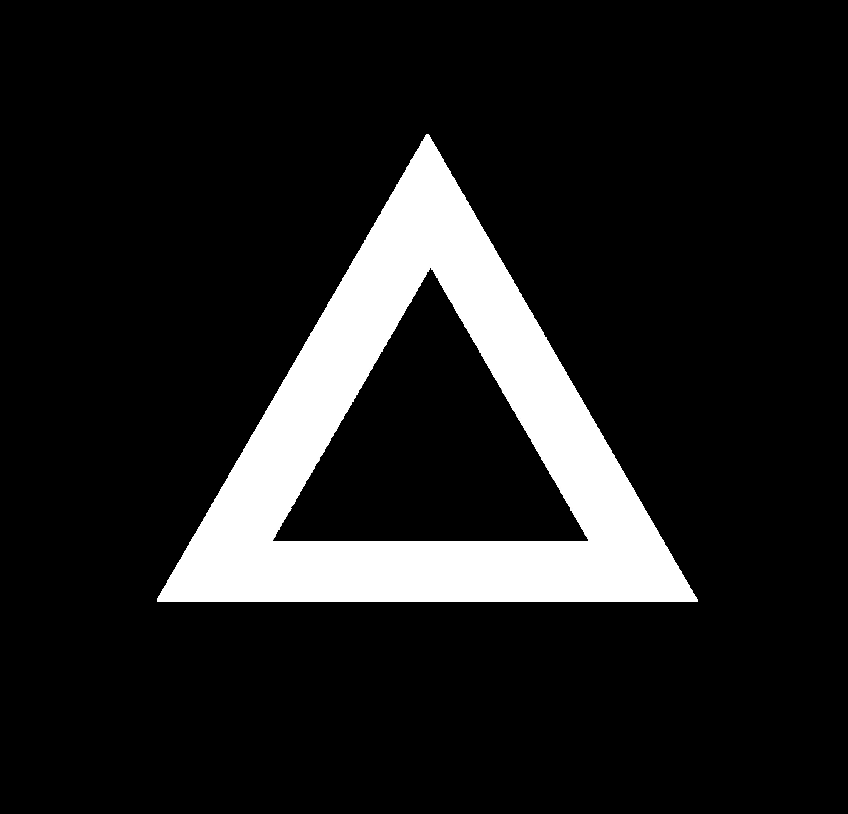

### Original synthetic data-10010.tif

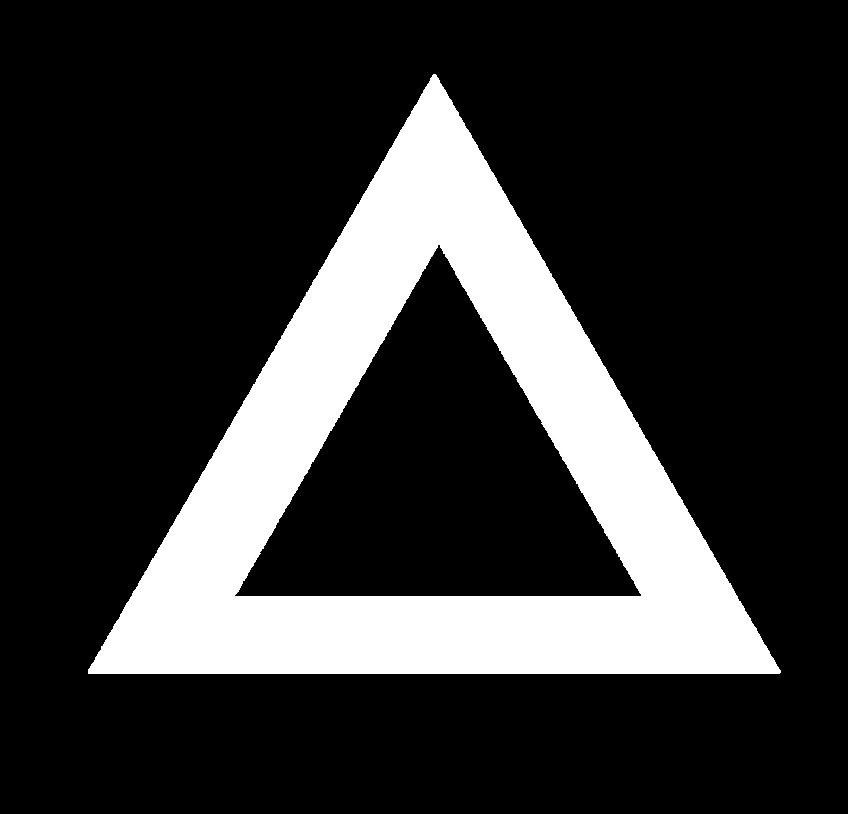

### Original synthetic data-10011.tif

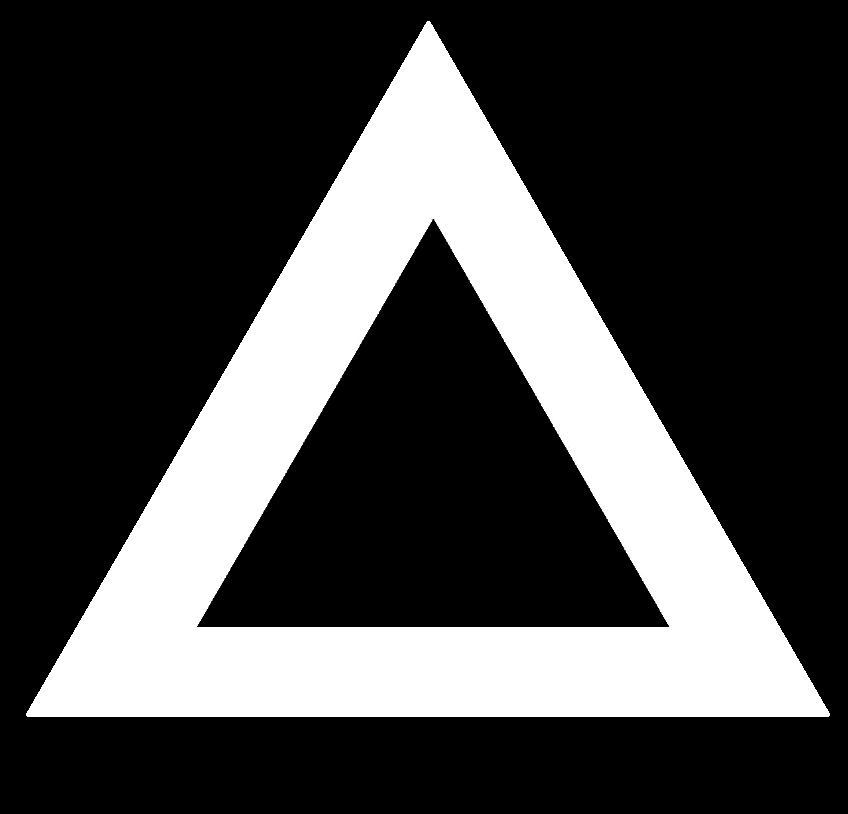

### Speckled blurred south shadow0000.tif

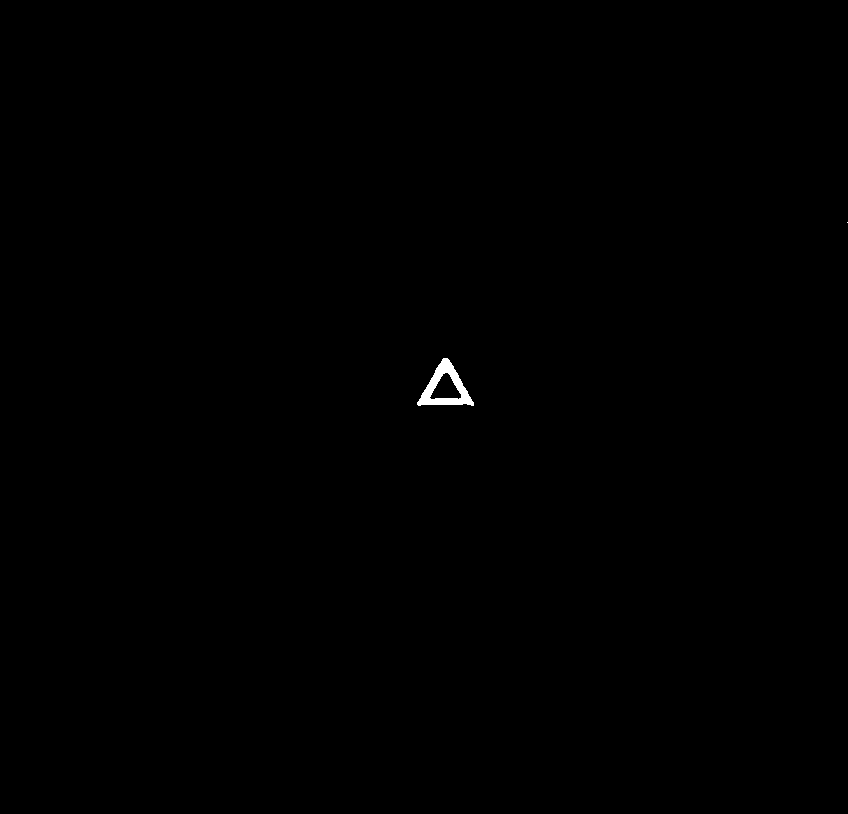

### Speckled blurred south shadow0001.tif

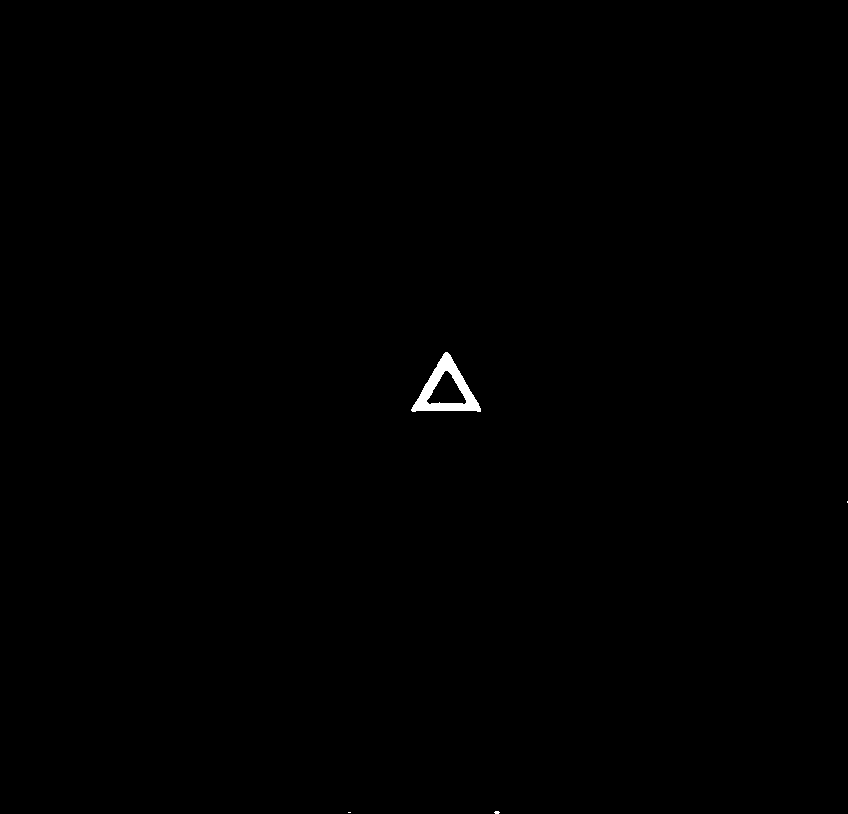

### Speckled blurred south shadow0002.tif

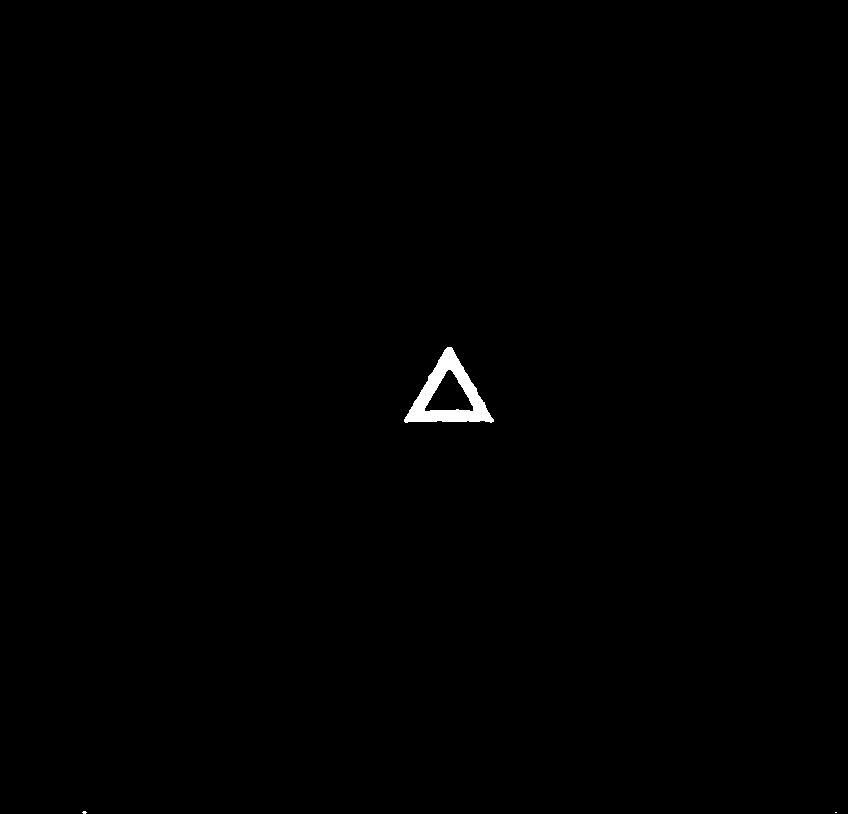

### Speckled blurred south shadow0003.tif

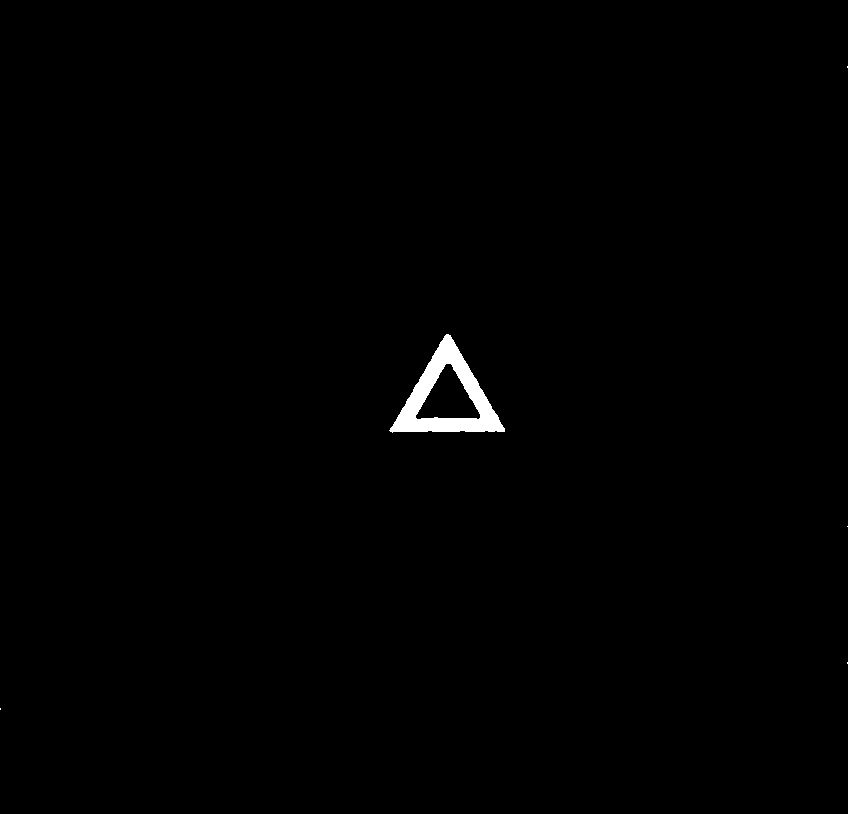

### Speckled blurred south shadow0004.tif

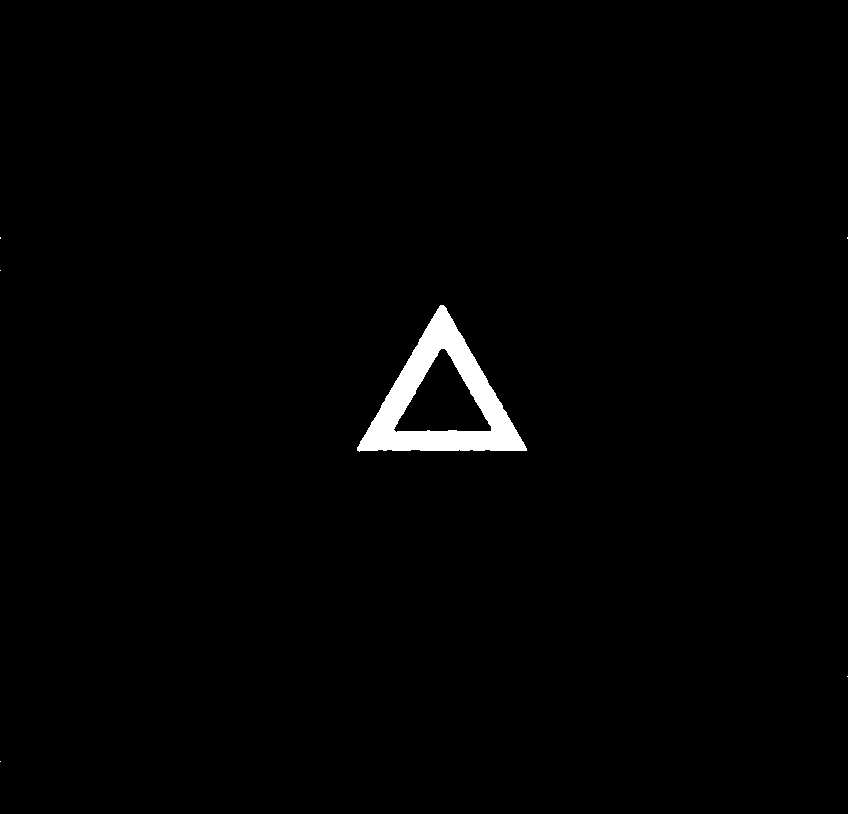

### Speckled blurred south shadow0005.tif

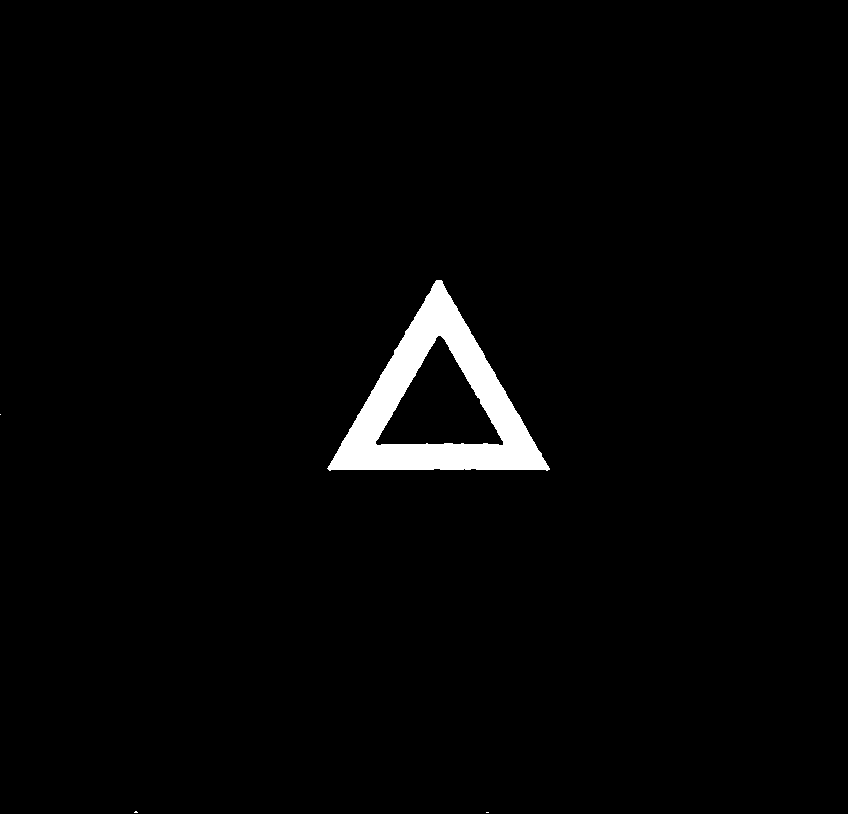

### Speckled blurred south shadow0006.tif

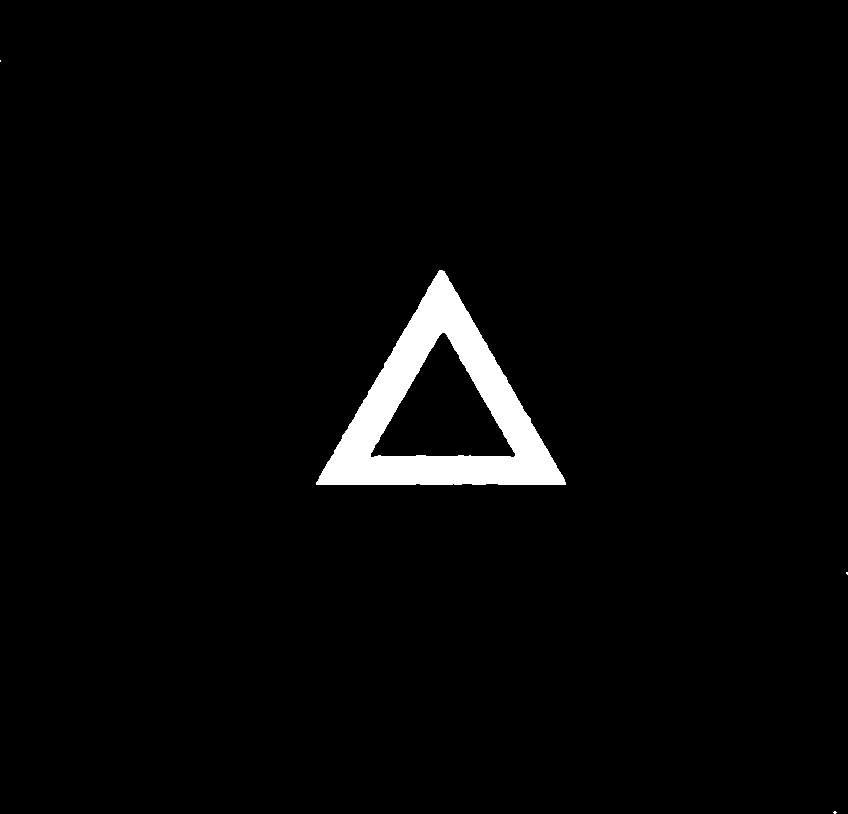

### Speckled blurred south shadow0007.tif

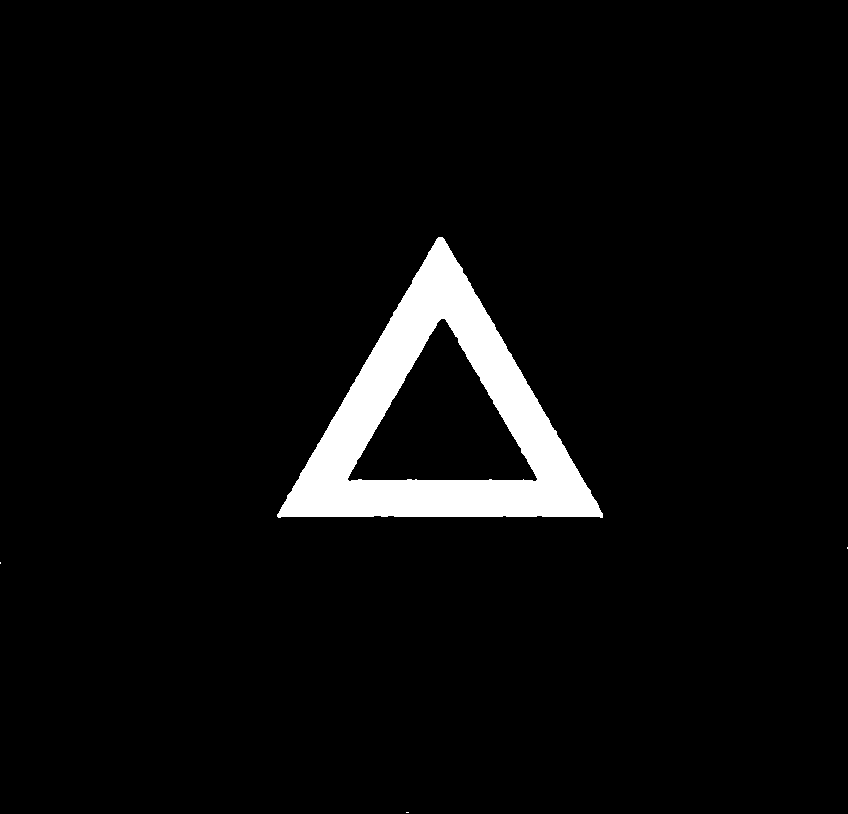

### Speckled blurred south shadow0008.tif

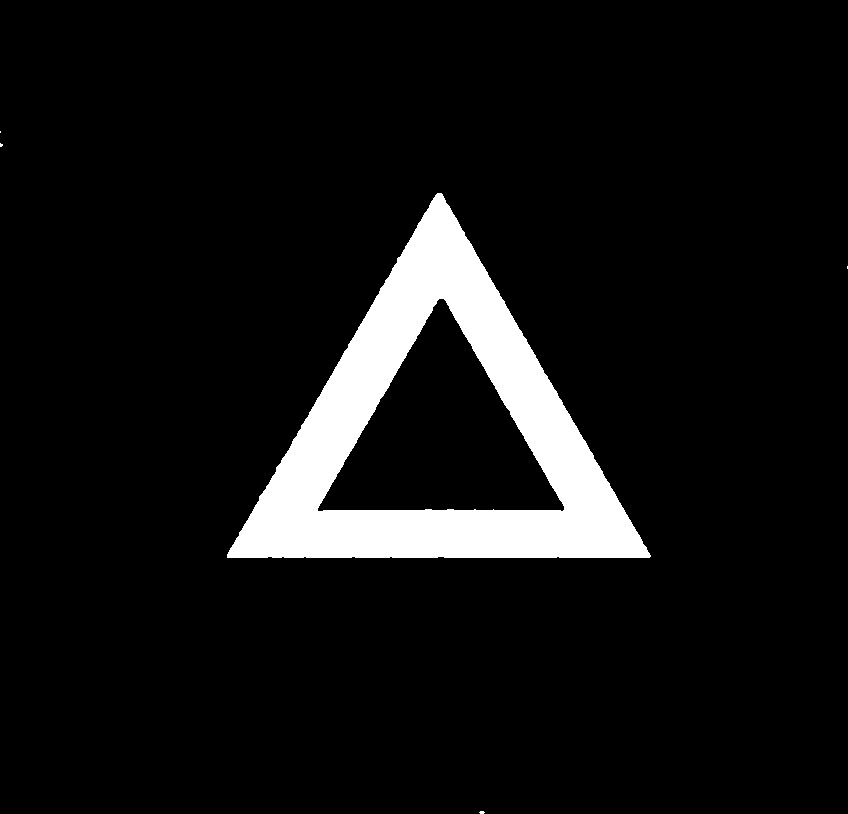

### Speckled blurred south shadow0009.tif

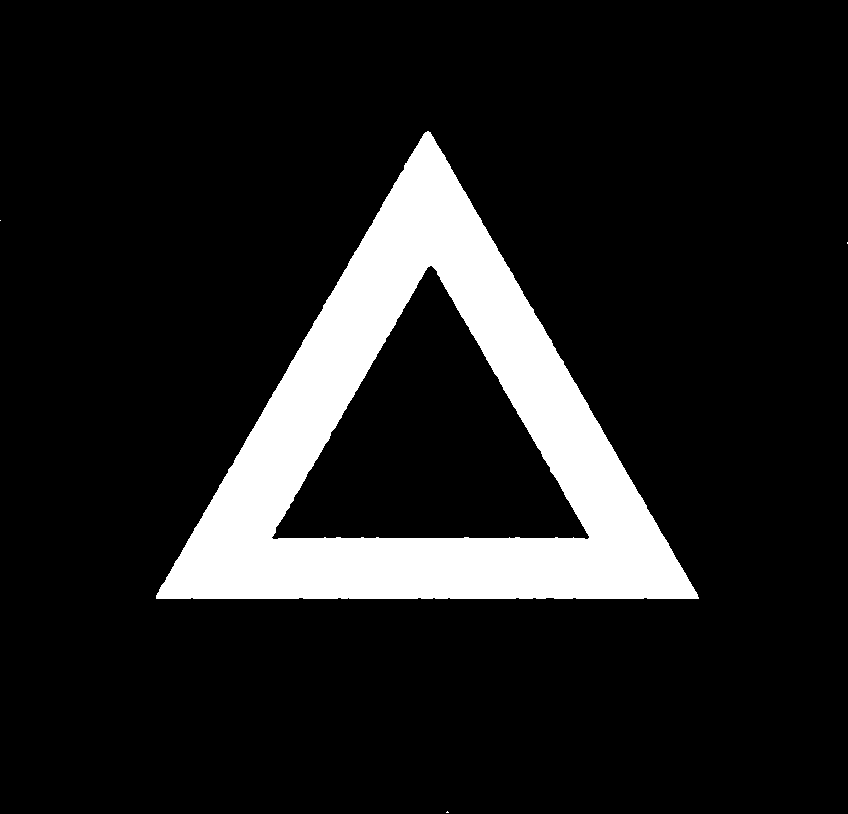

### Speckled blurred south shadow0010.tif

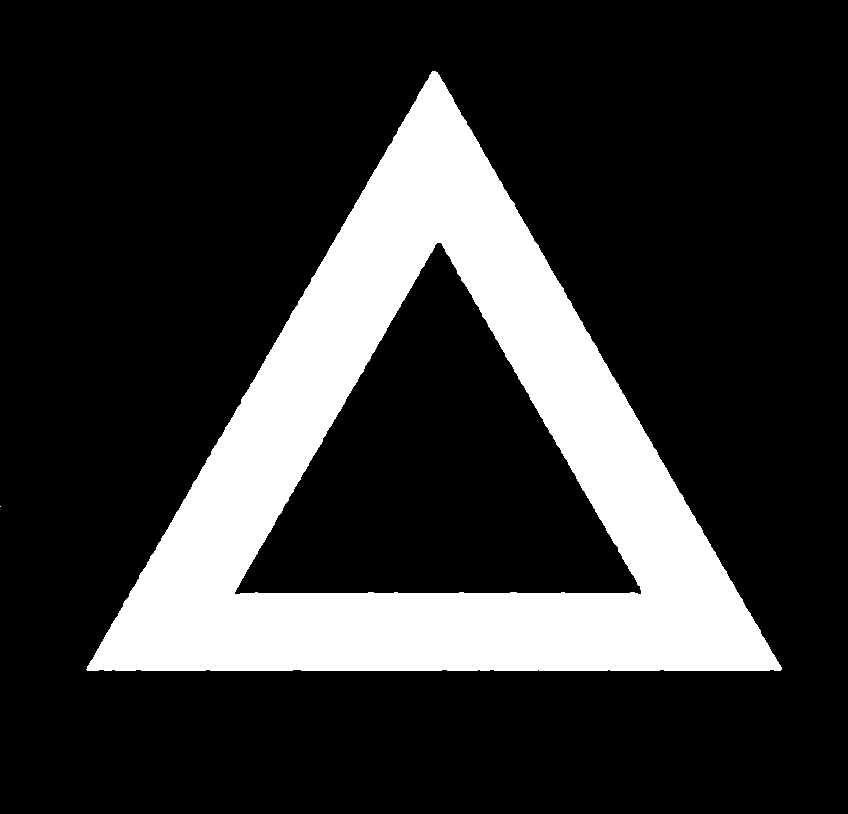

### Speckled blurred south shadow0011.tif

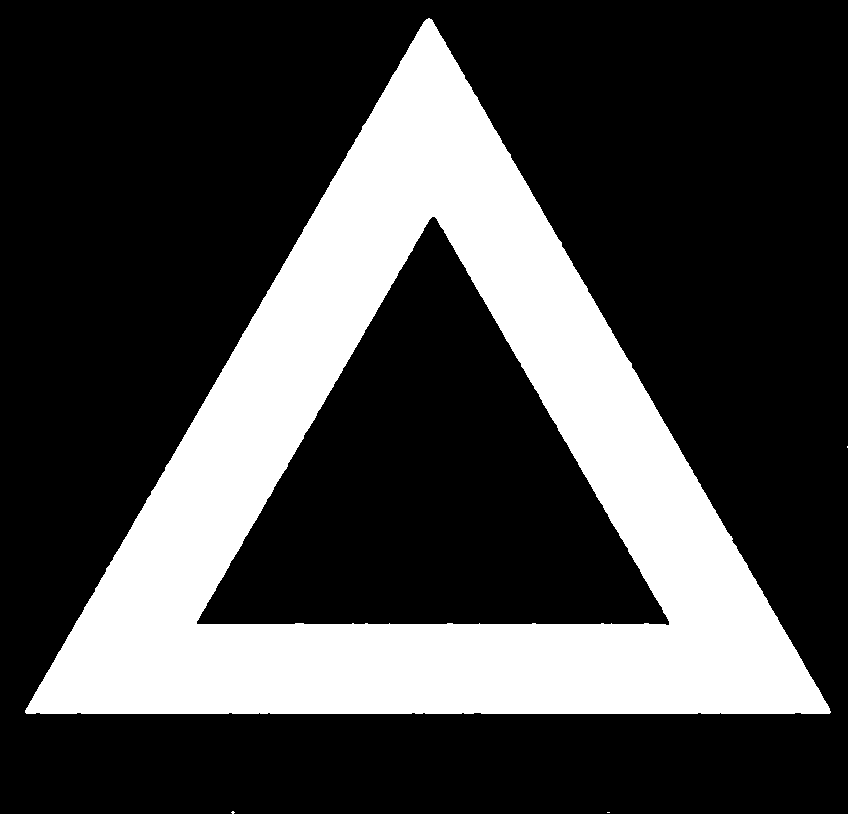

### Speckled,blurred,south shadow0000.tif

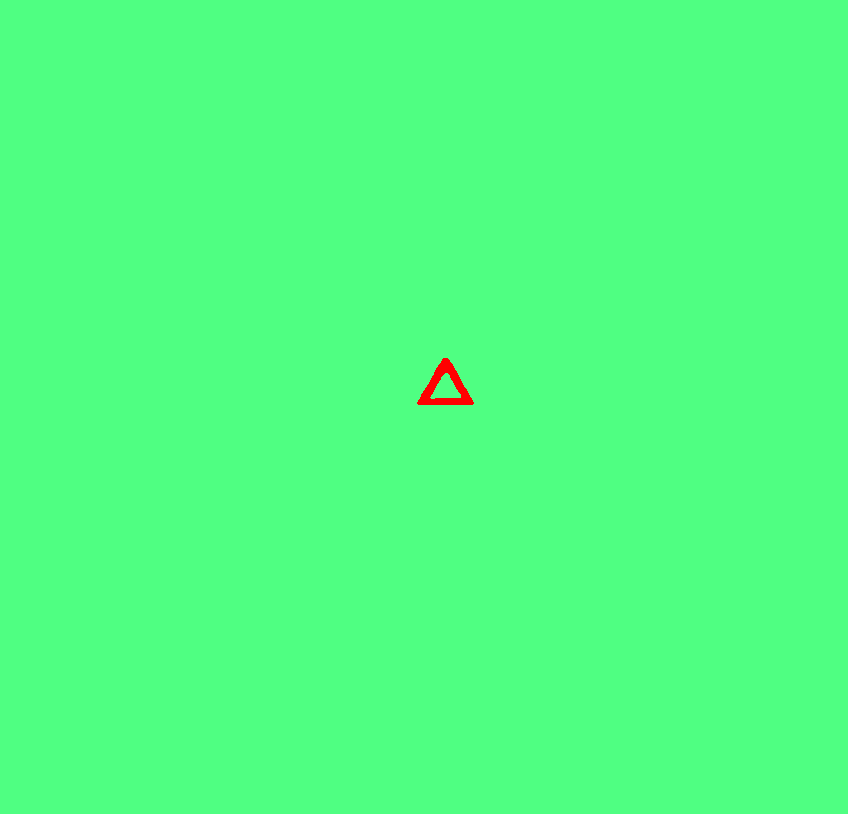

### Speckled,blurred,south shadow0000.tif

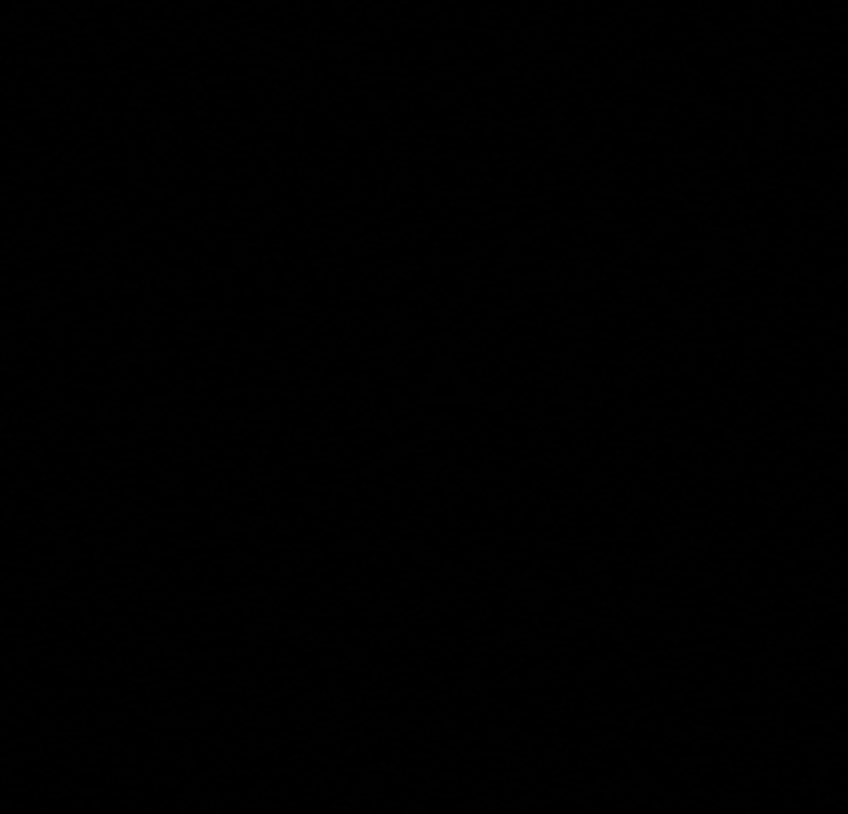

### Speckled,blurred,south shadow0001.tif

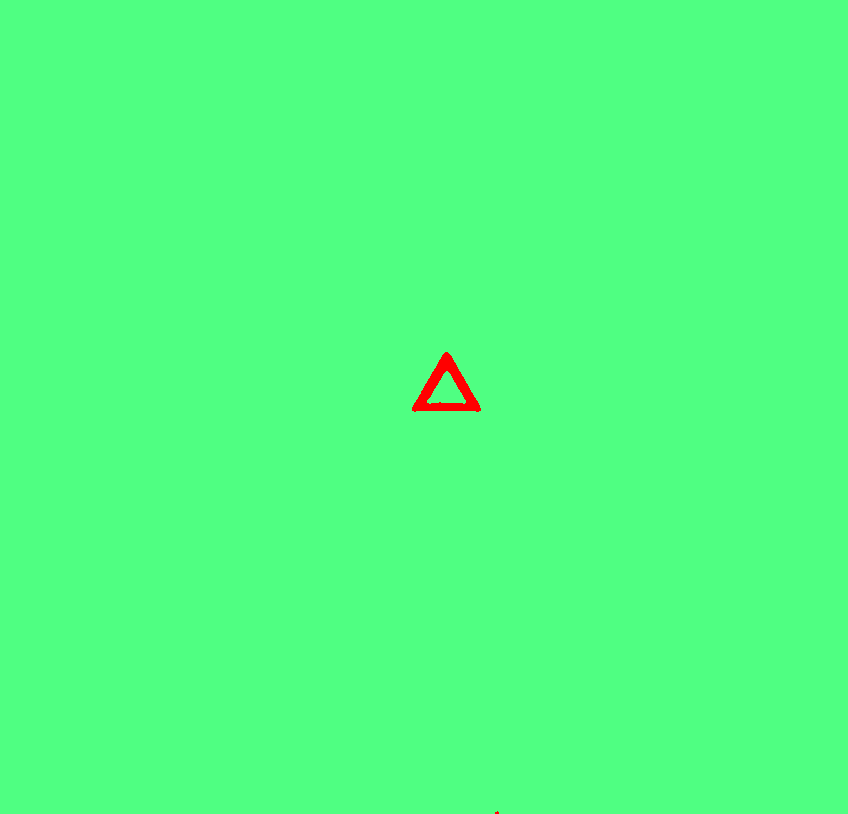

### Speckled,blurred,south shadow0001.tif

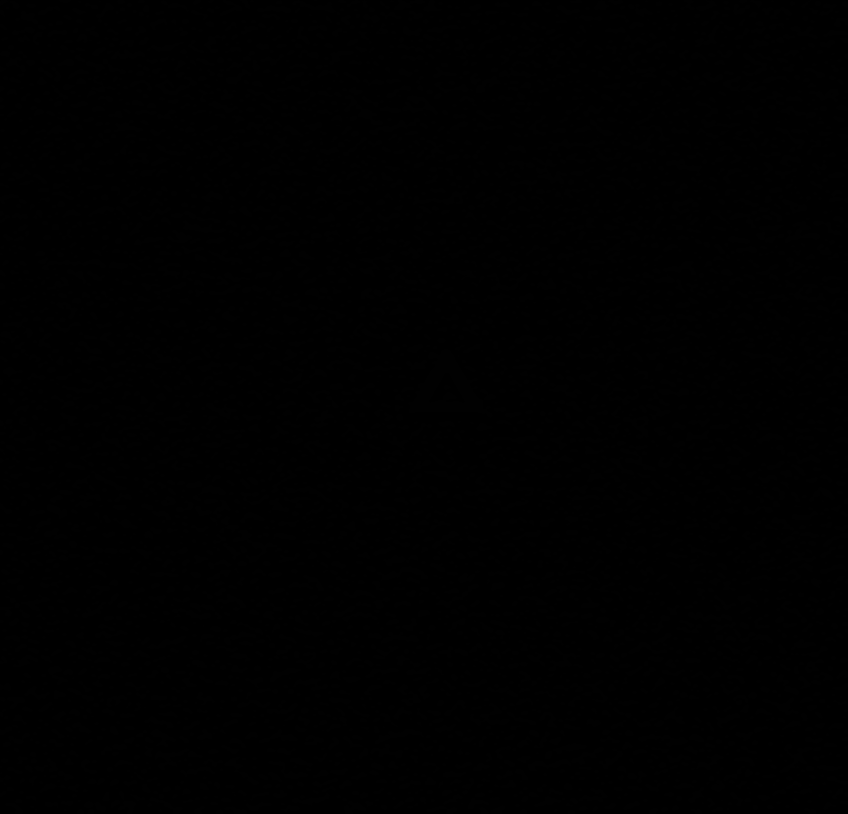

### Speckled,blurred,south shadow0002.tif

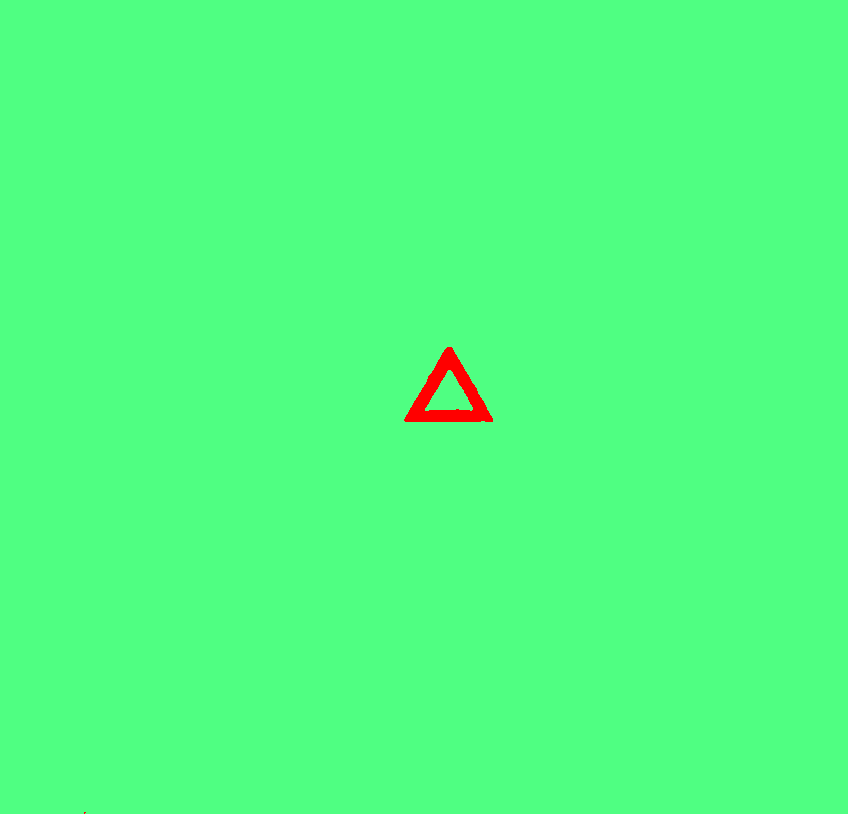

### Speckled,blurred,south shadow0002.tif

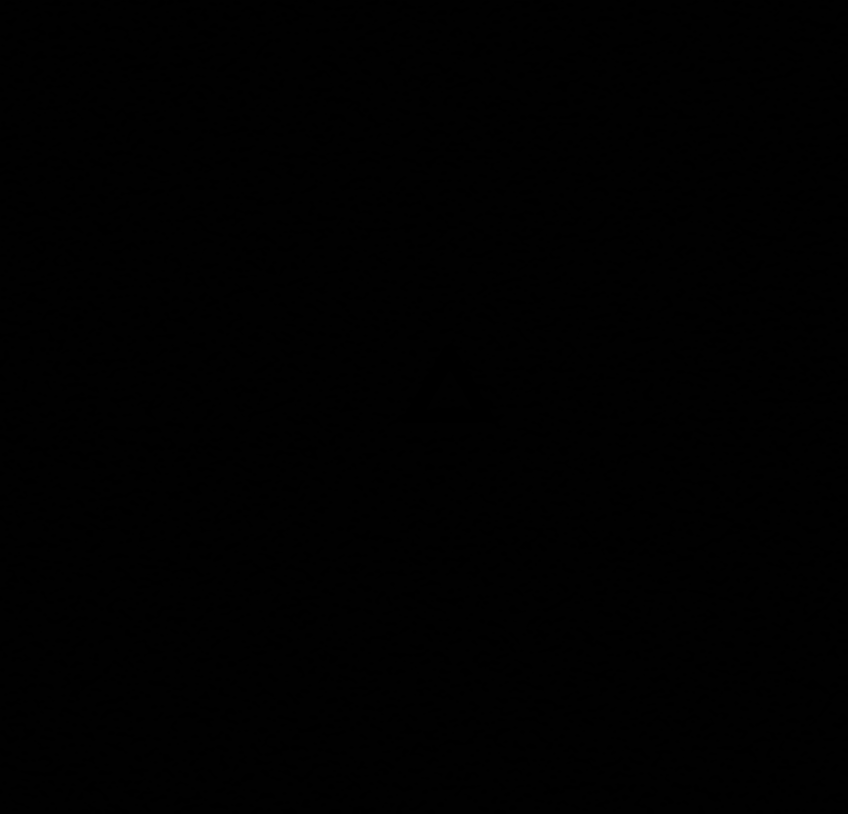

### Speckled,blurred,south shadow0003.tif

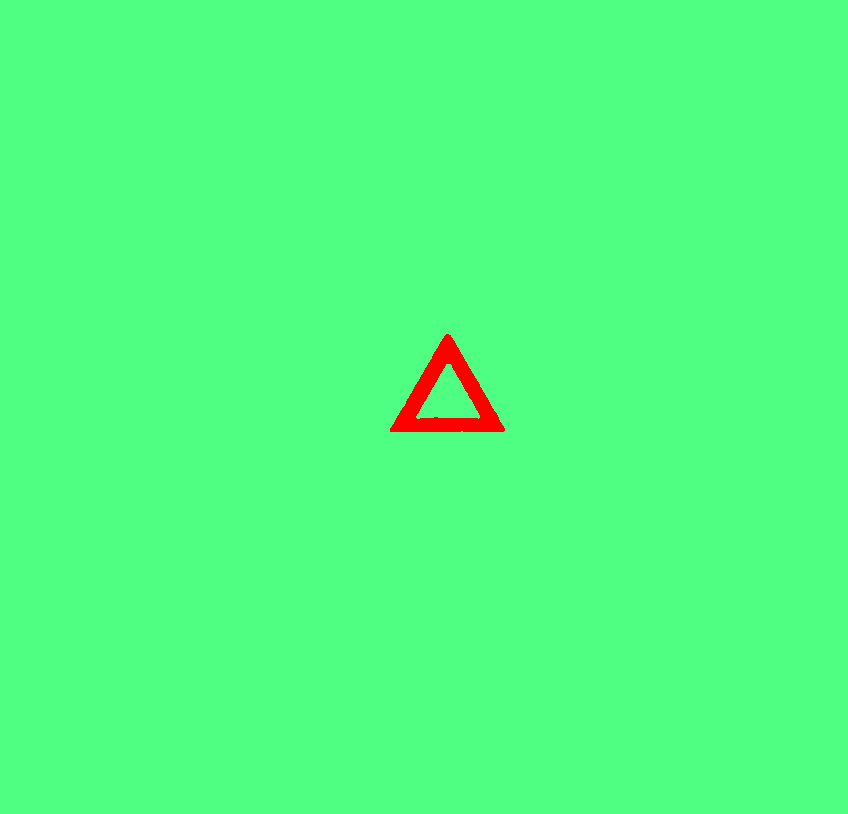

### Speckled,blurred,south shadow0003.tif

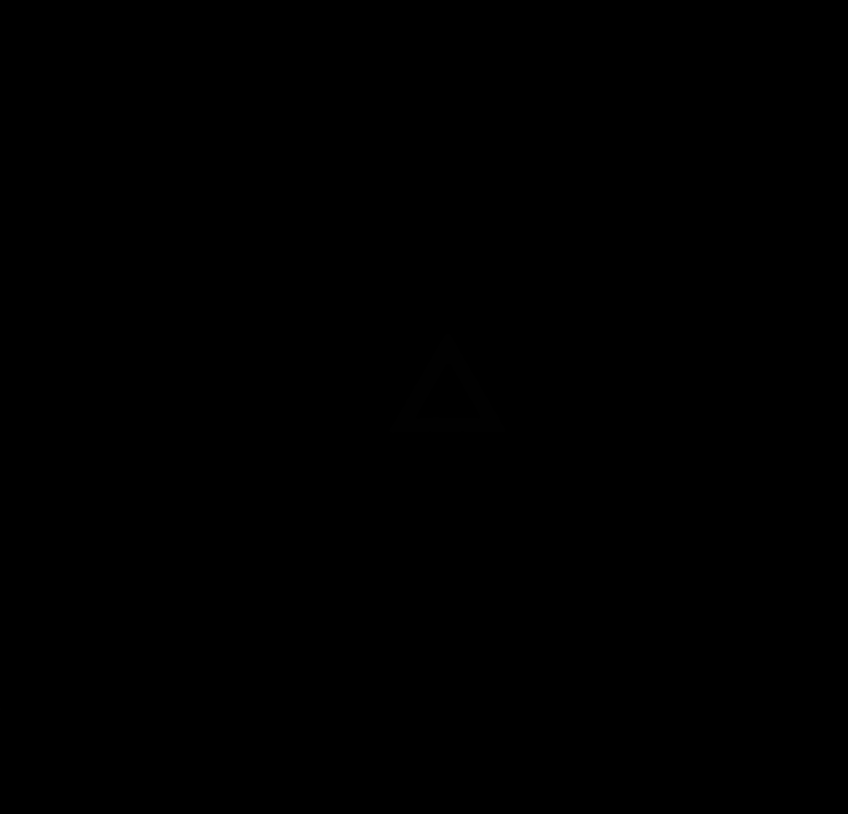

### Speckled,blurred,south shadow0004.tif

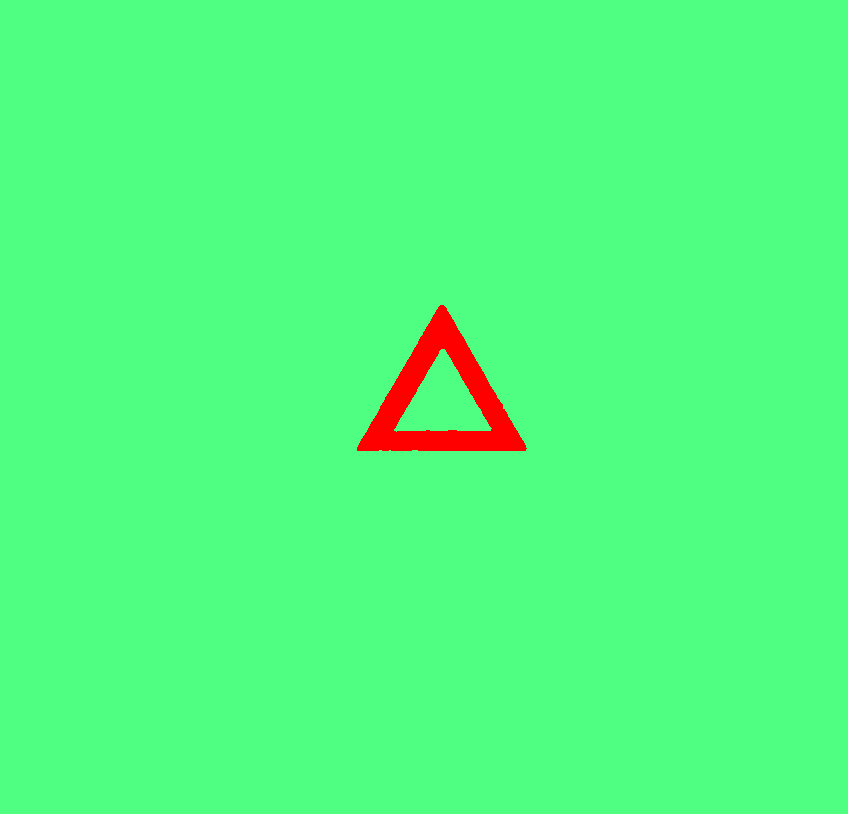

### Speckled,blurred,south shadow0004.tif

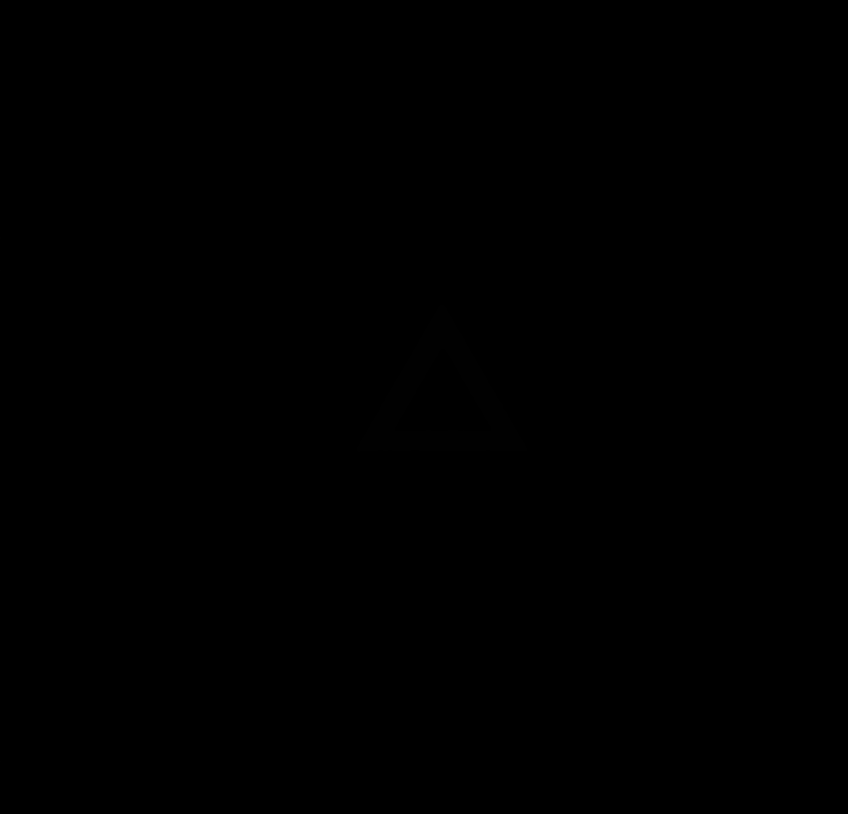

### Speckled,blurred,south shadow0005.tif

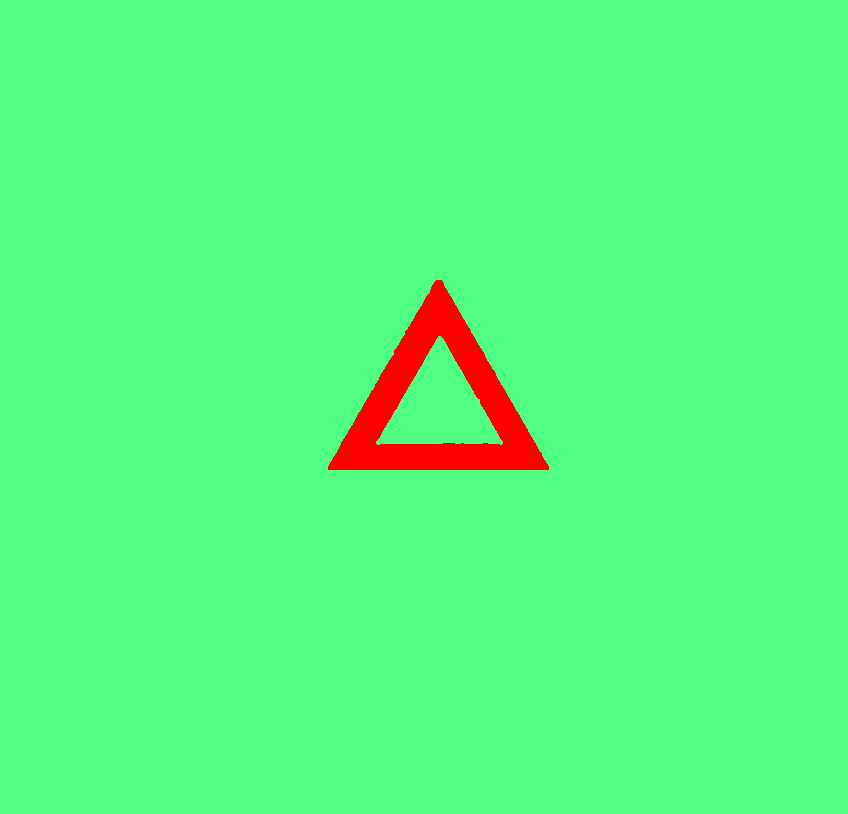

### Speckled,blurred,south shadow0005.tif

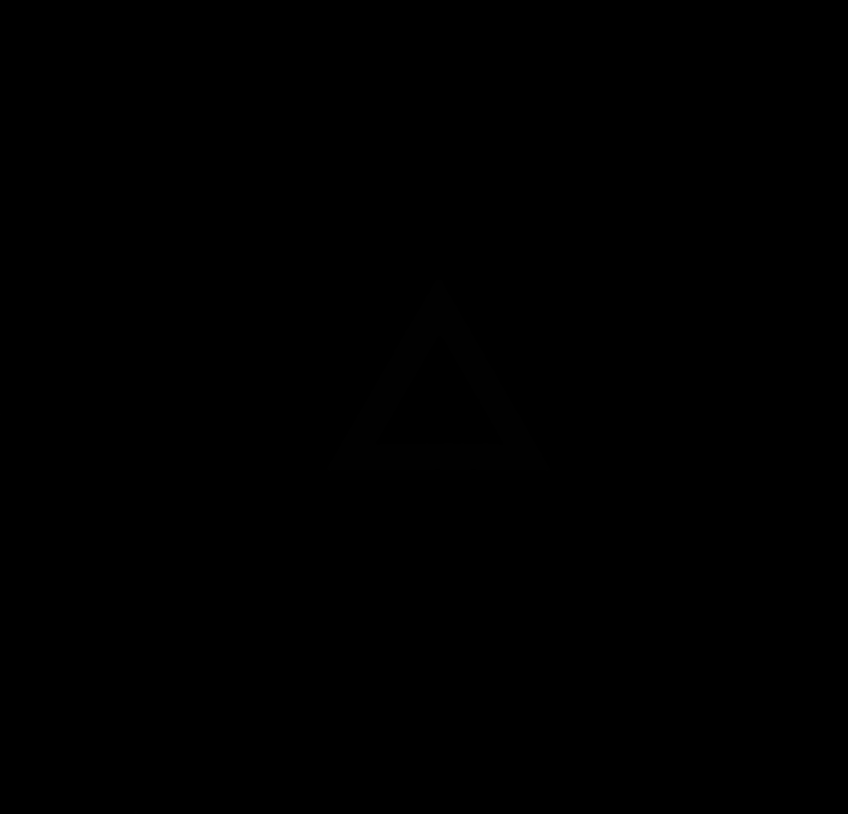

### Speckled,blurred,south shadow0006.tif

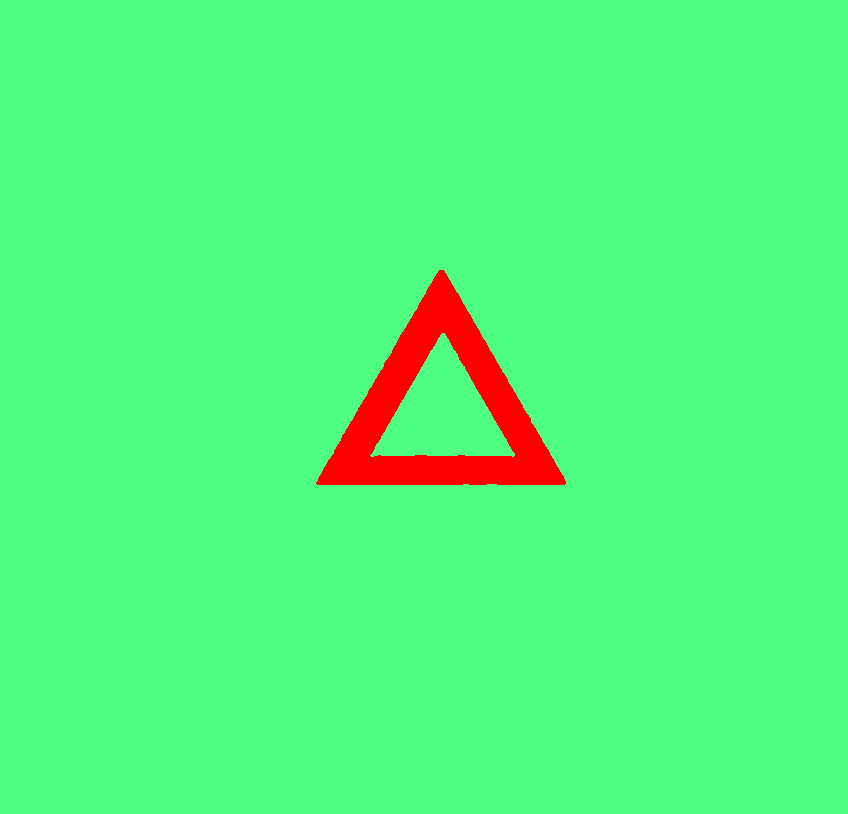

### Speckled,blurred,south shadow0006.tif

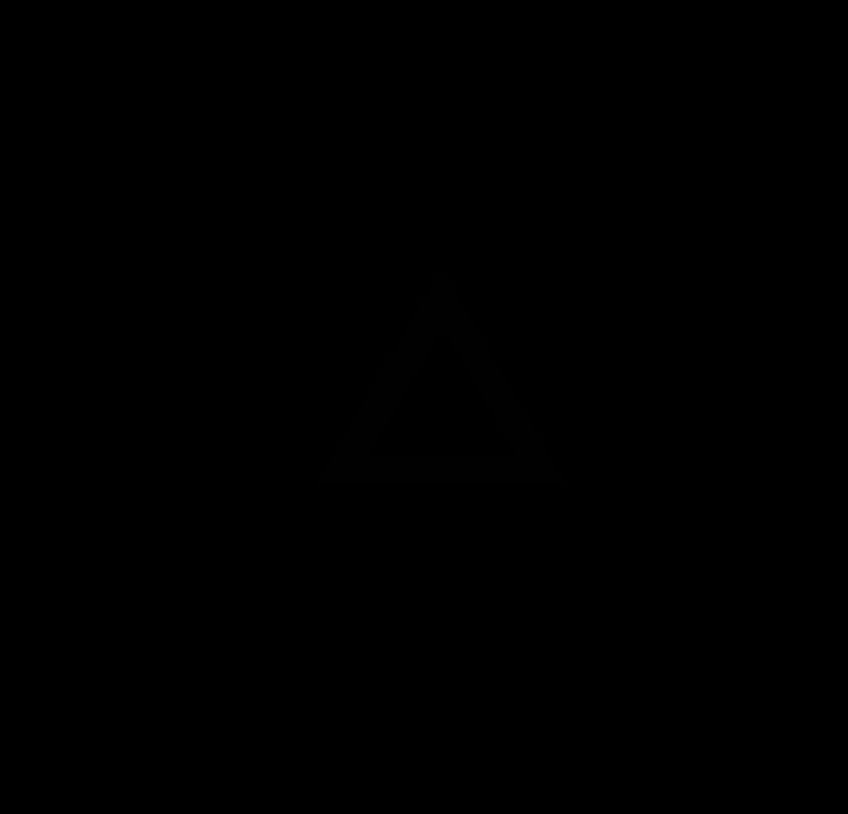

### Speckled,blurred,south shadow0007.tif

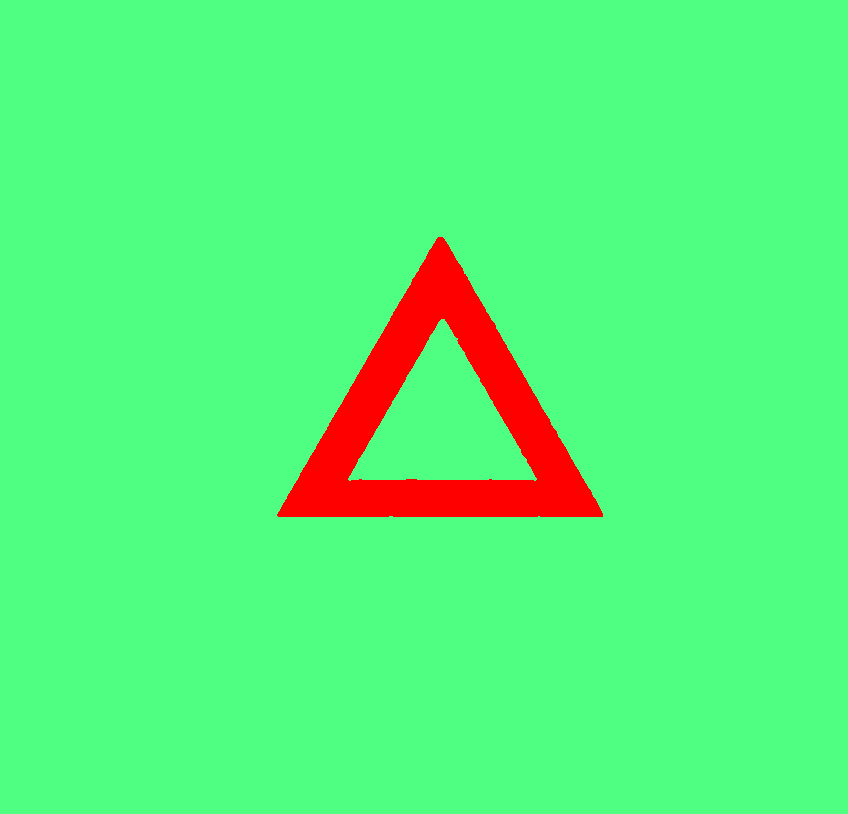

### Speckled,blurred,south shadow0007.tif

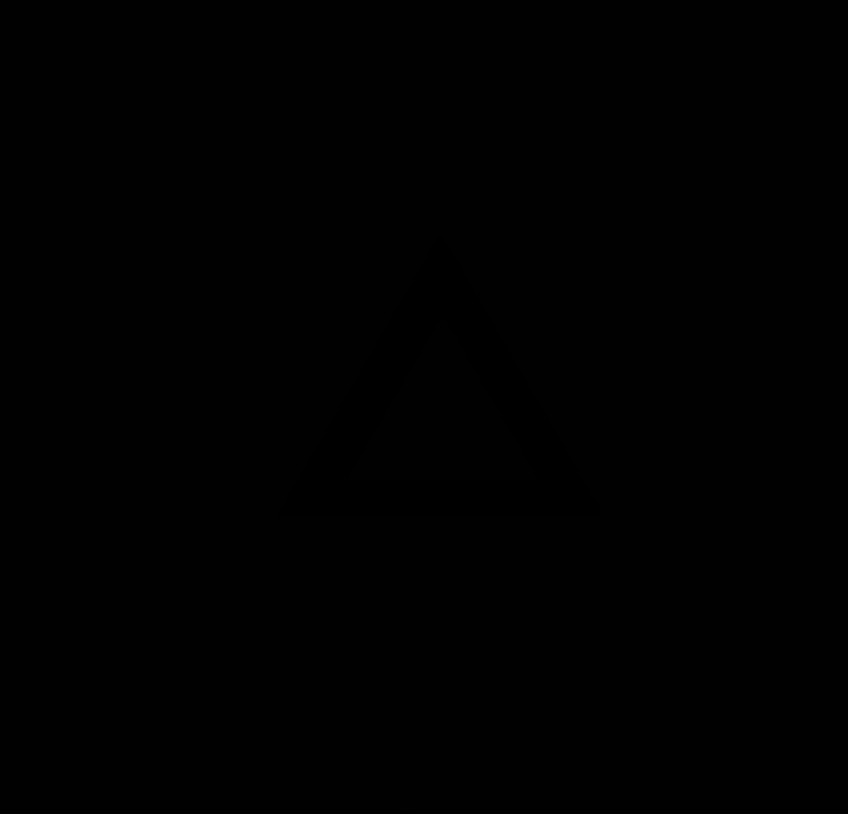

### Speckled,blurred,south shadow0008.tif

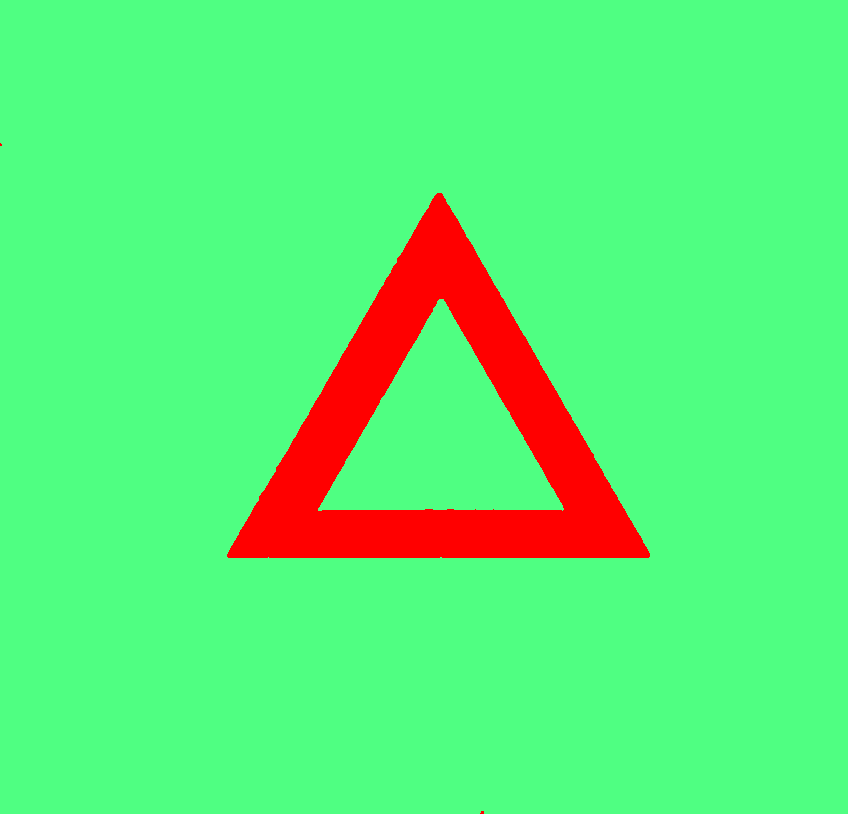

### Speckled,blurred,south shadow0008.tif

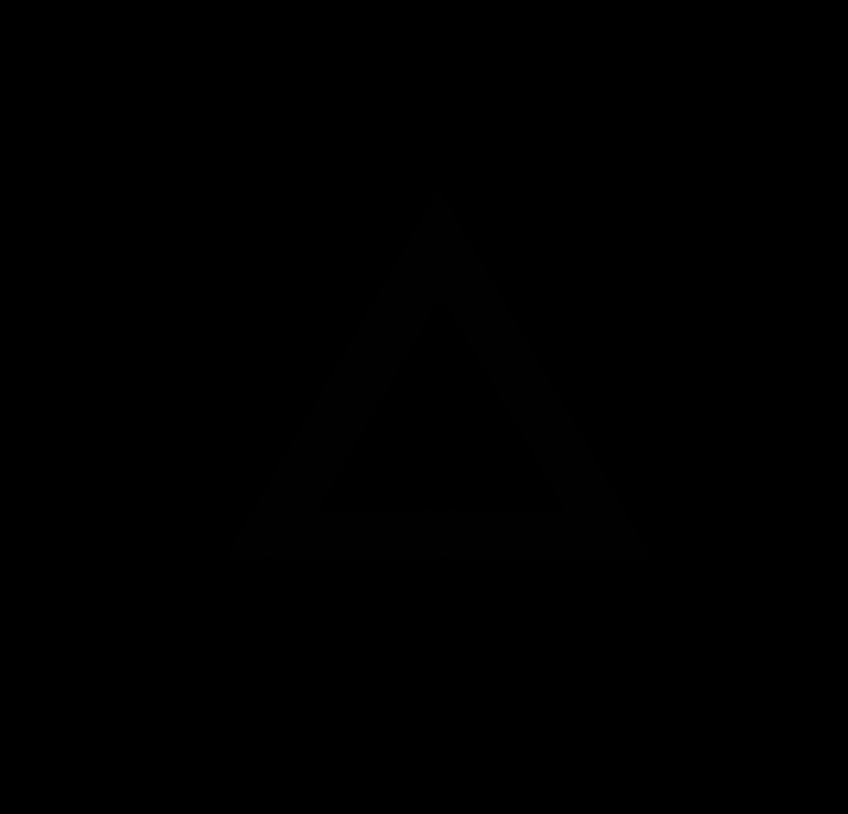

### Speckled,blurred,south shadow0009.tif

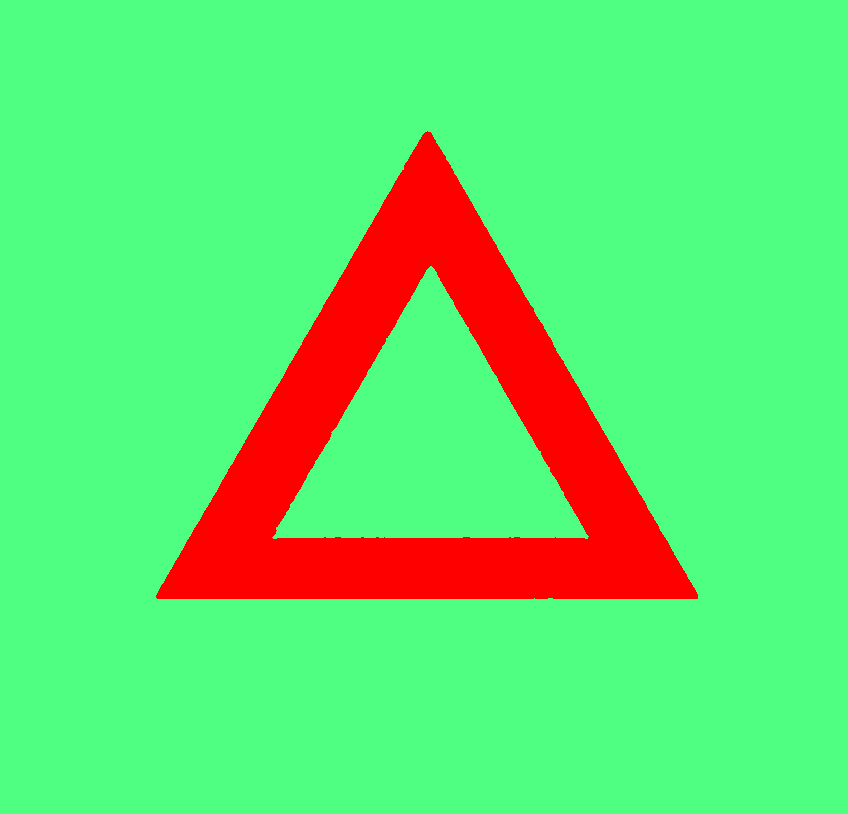

### Speckled,blurred,south shadow0009.tif

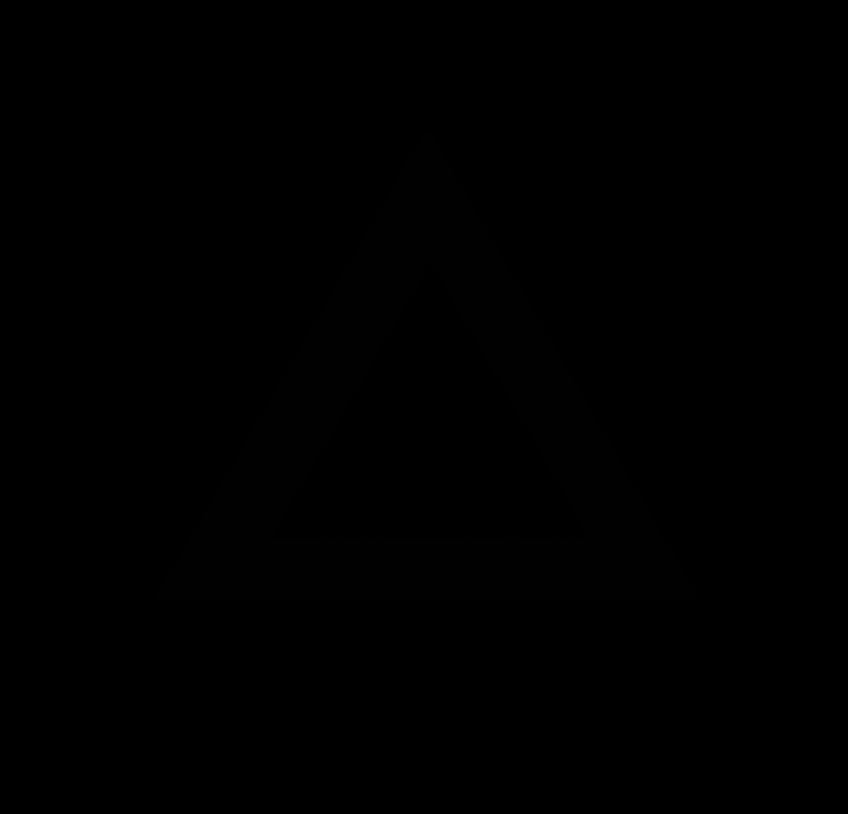

### Speckled,blurred,south shadow0010.tif

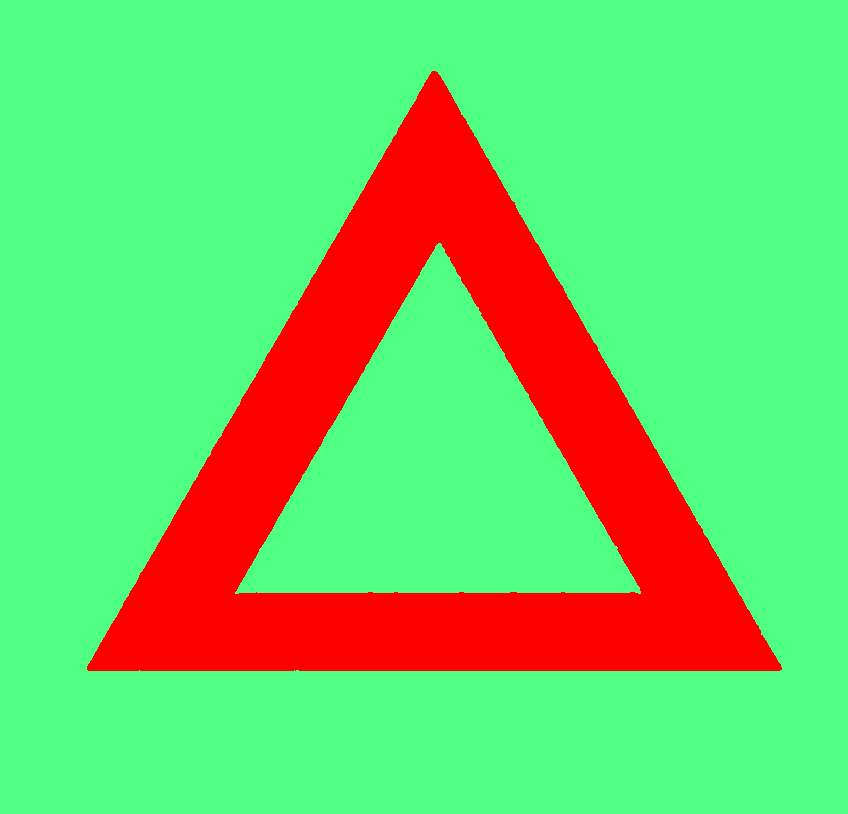

### Speckled,blurred,south shadow0010.tif

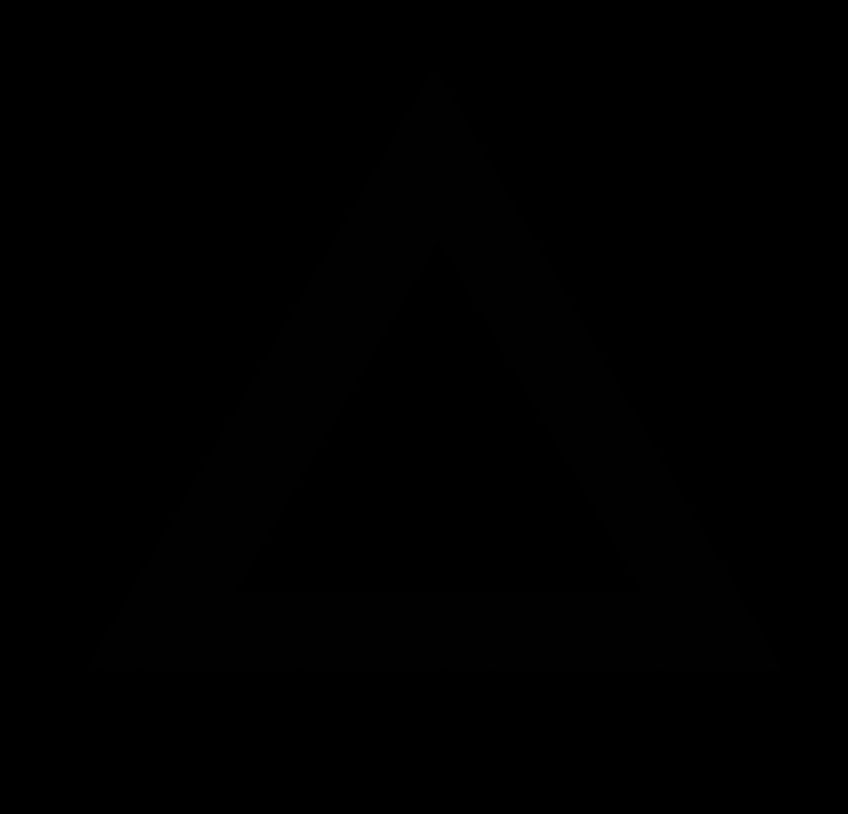

### Speckled,blurred,south shadow0011.tif

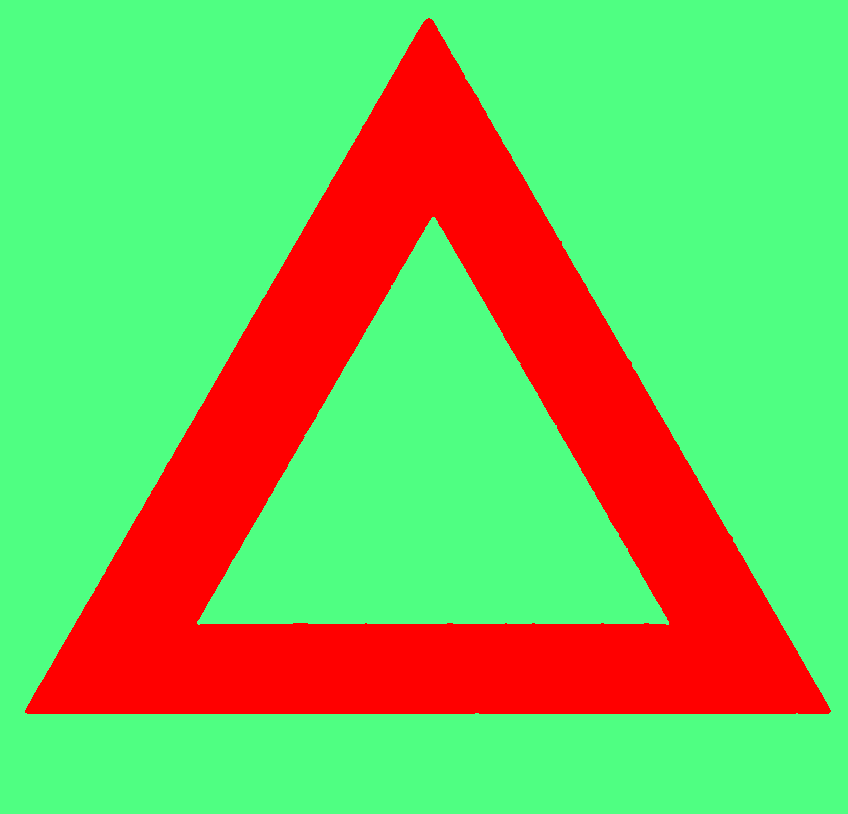

### Speckled,blurred,south shadow0011.tif

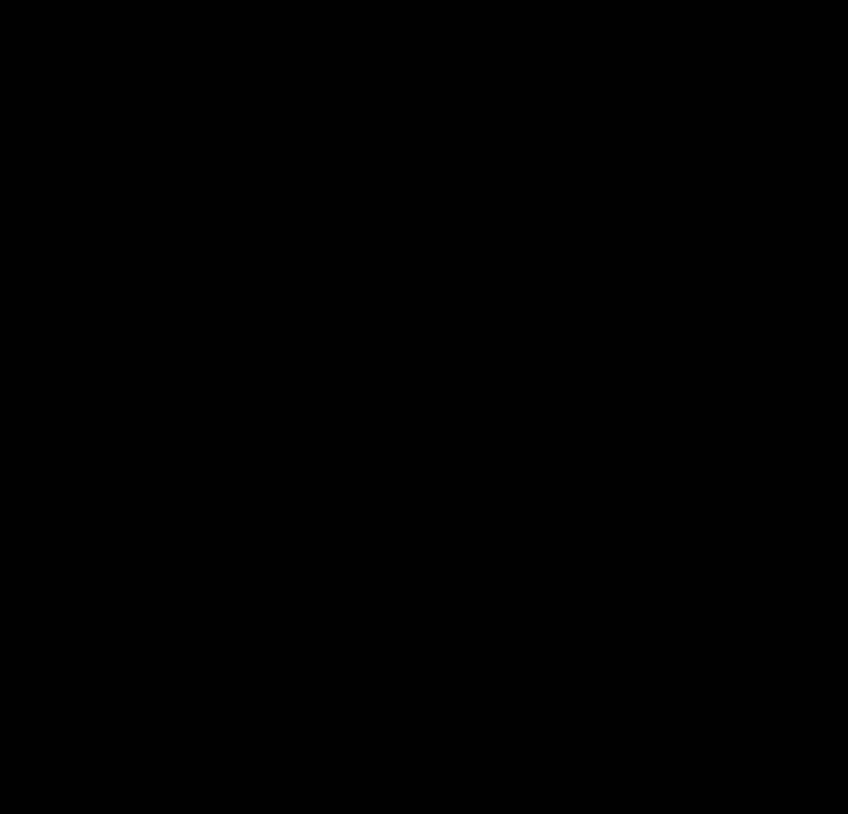

### syntheticlabelsgs12-10000.tif

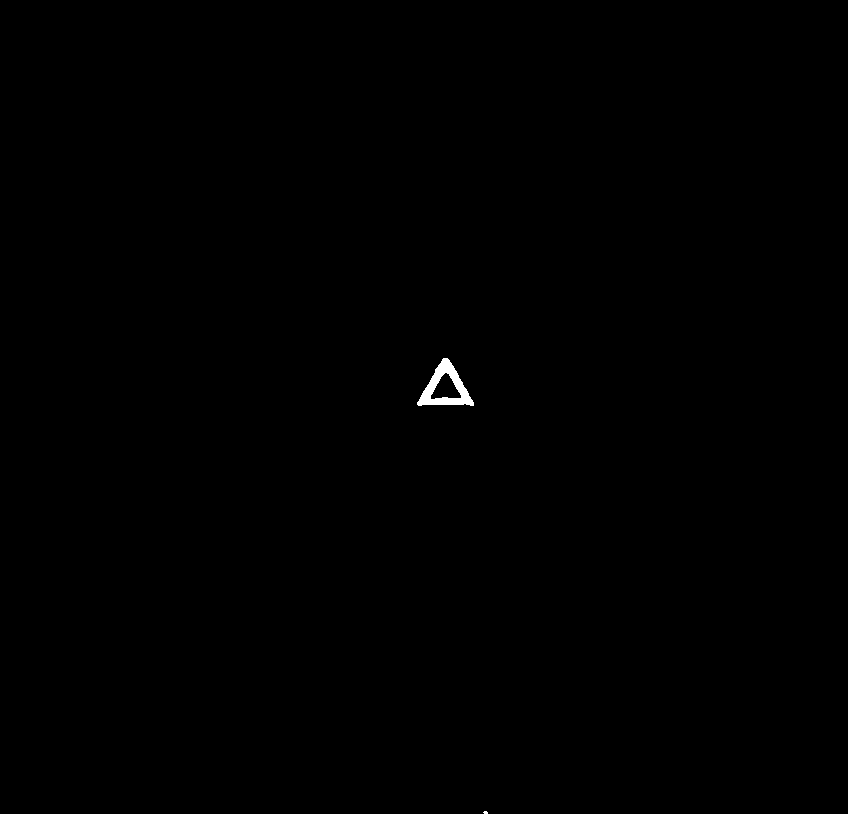

### syntheticlabelsgs12-10001.tif

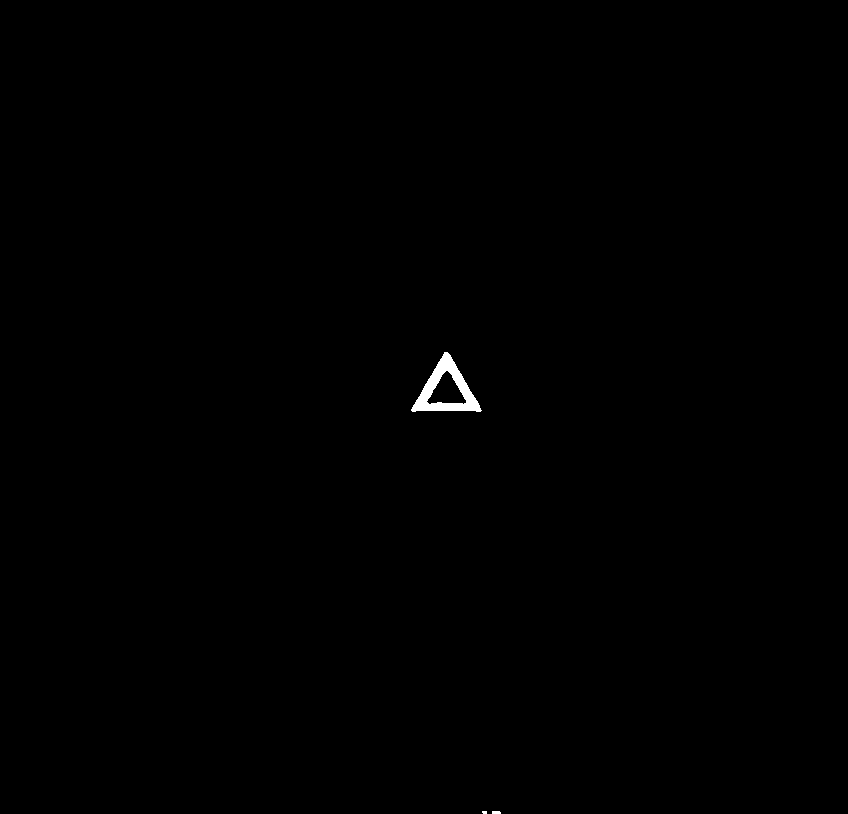

### syntheticlabelsgs12-10002.tif

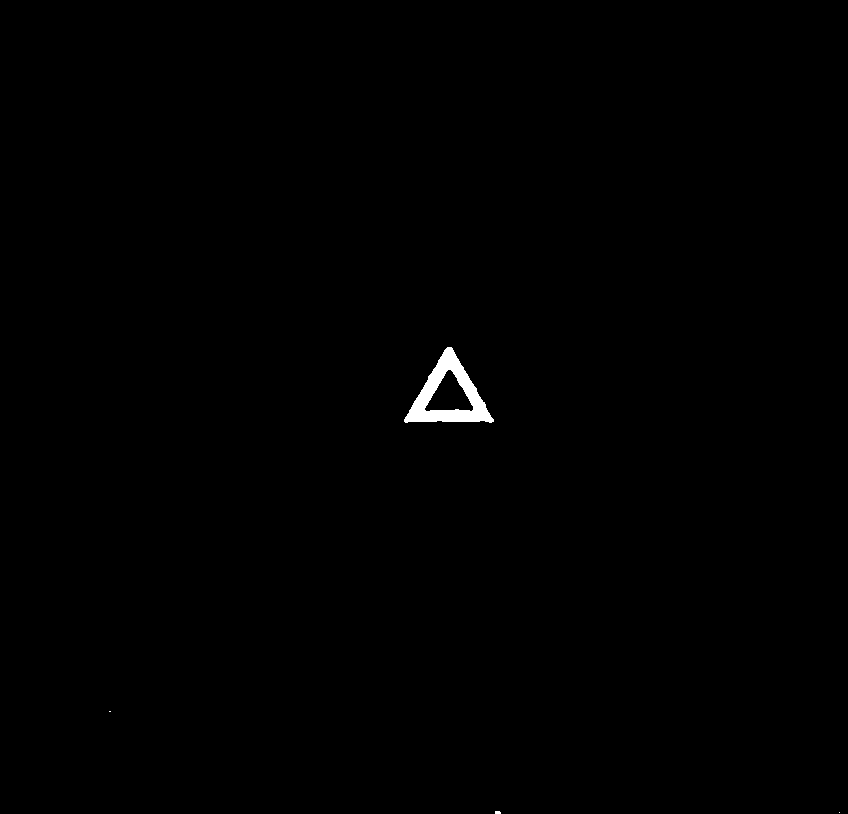

### syntheticlabelsgs12-10003.tif

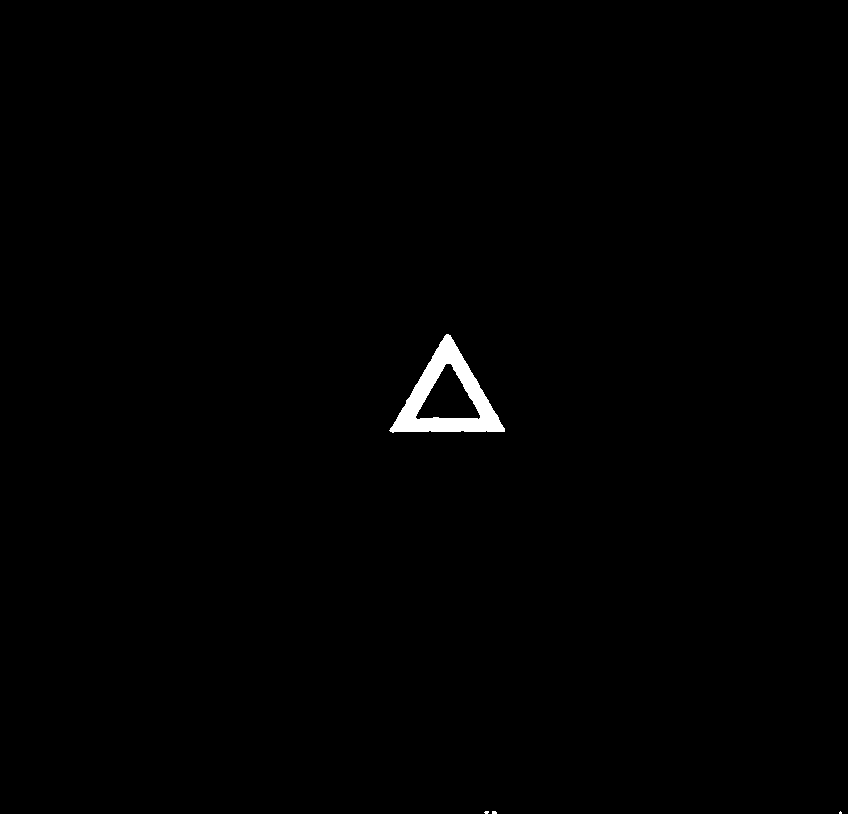

### syntheticlabelsgs12-10004.tif

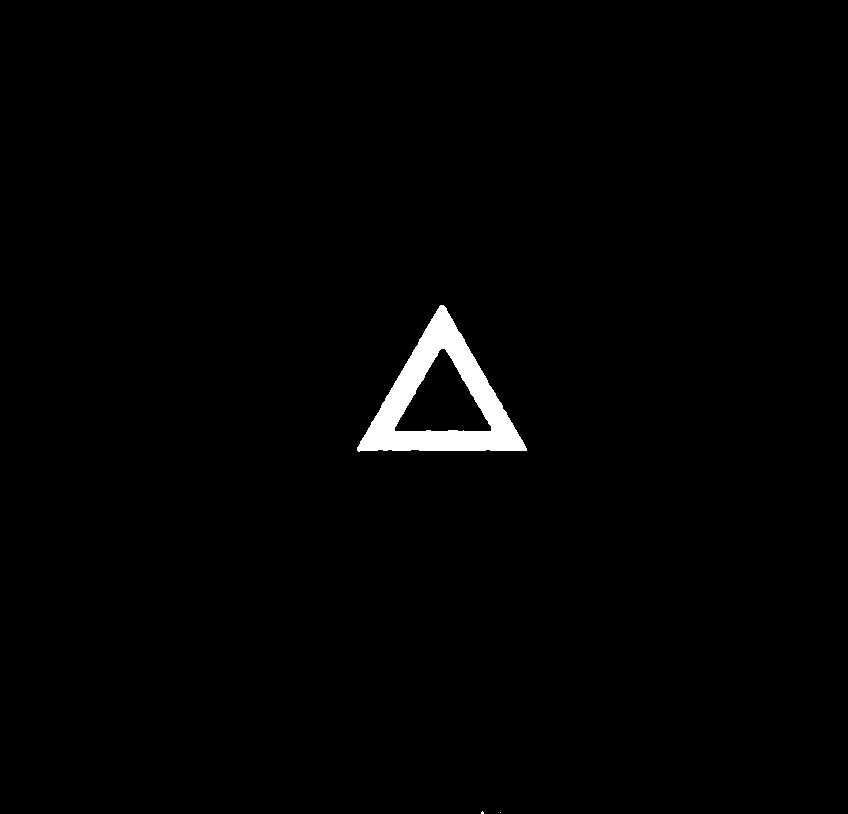

### syntheticlabelsgs12-10005.tif

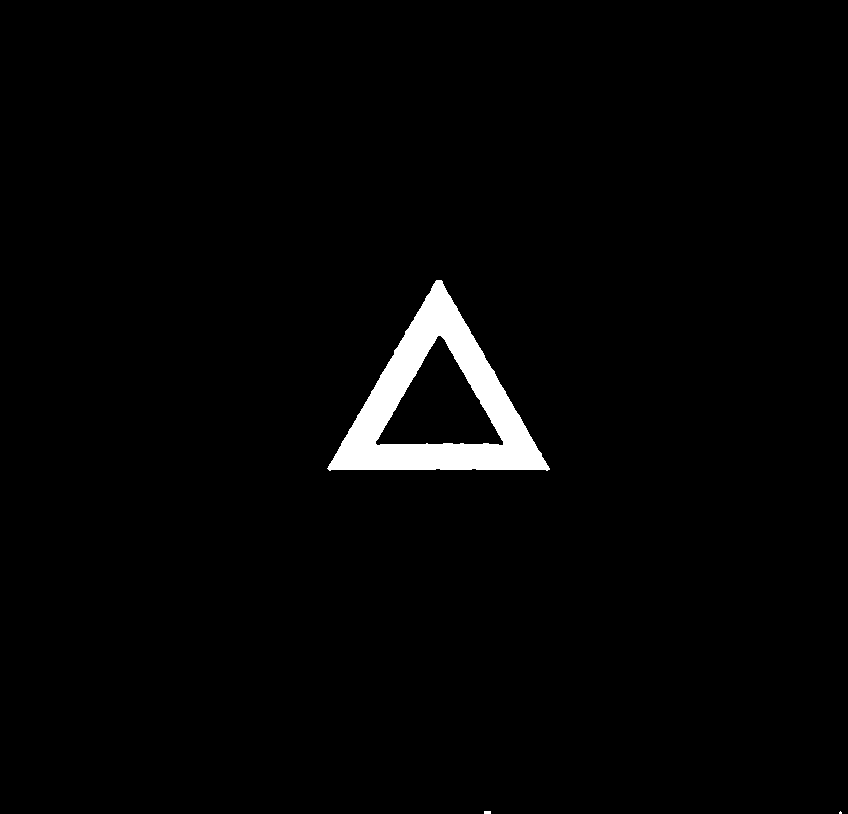

### syntheticlabelsgs12-10006.tif

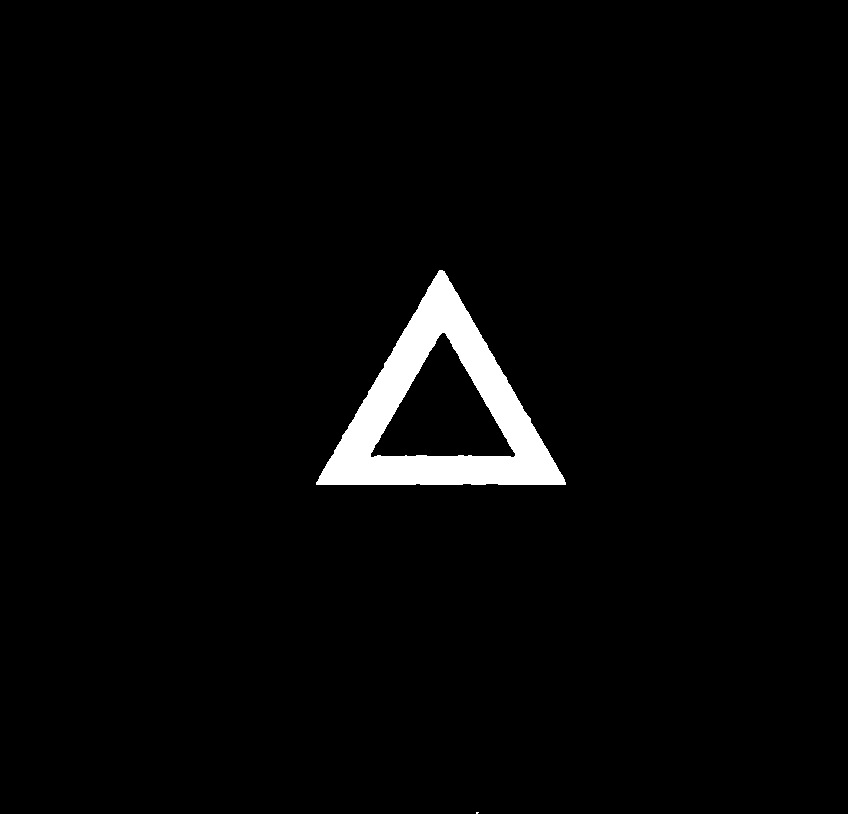

### syntheticlabelsgs12-10007.tif

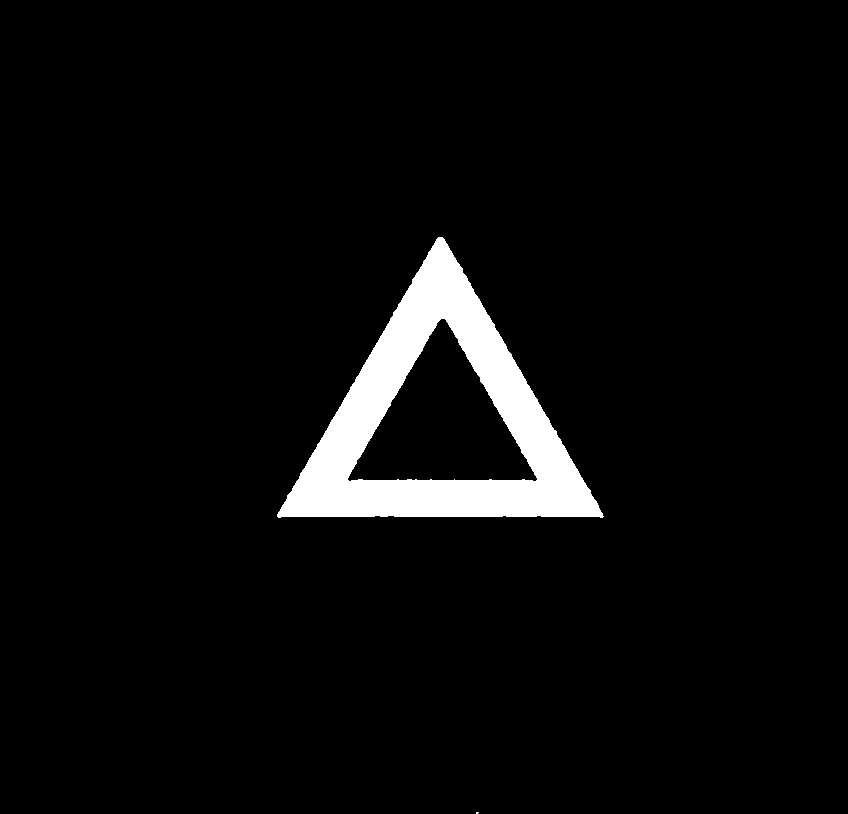

### syntheticlabelsgs12-10008.tif

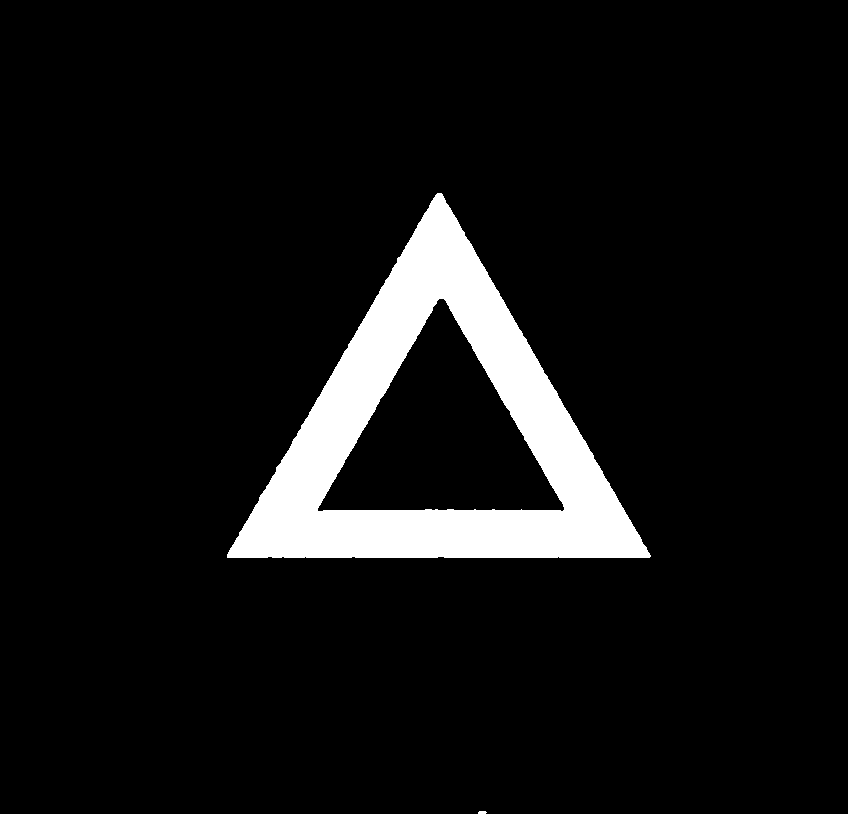

### syntheticlabelsgs12-10009.tif

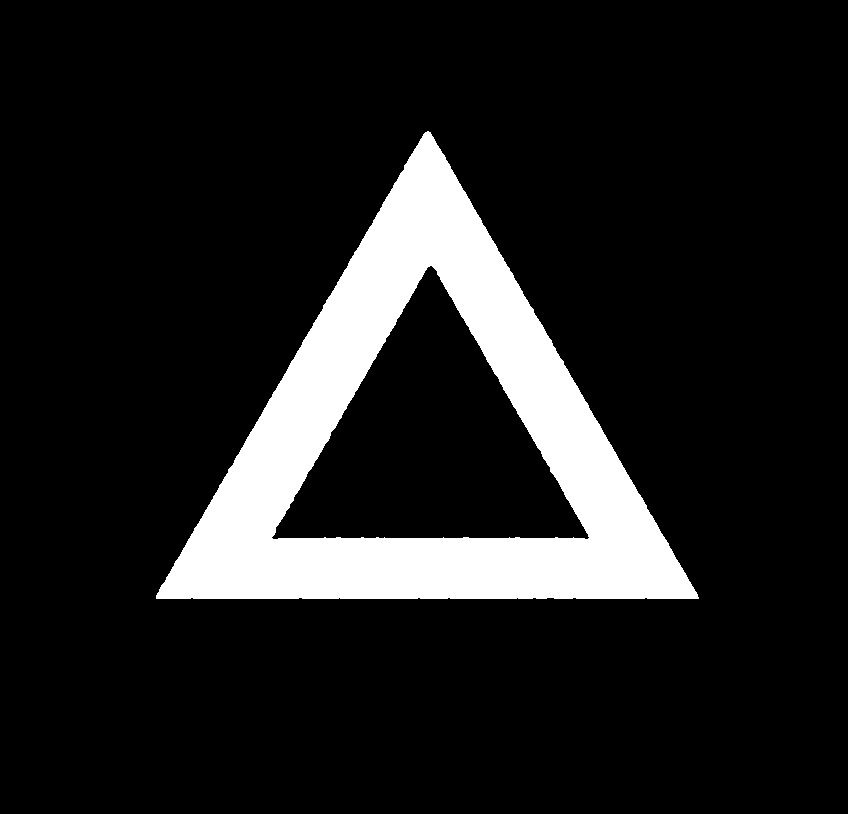

### syntheticlabelsgs12-10010.tif

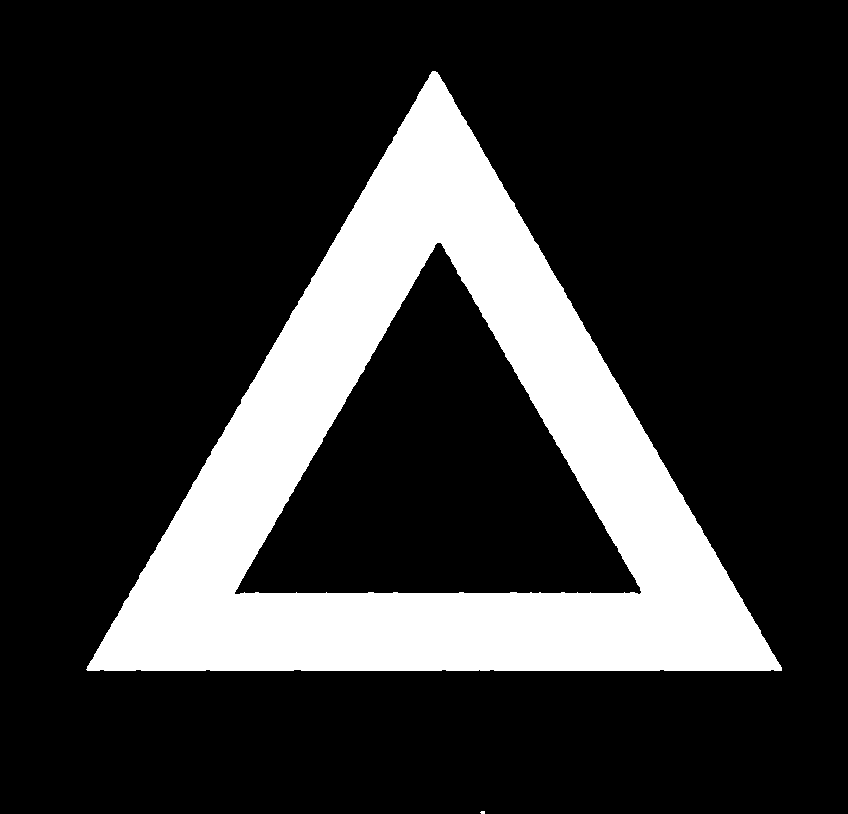

### syntheticlabelsgs12-10011.tif

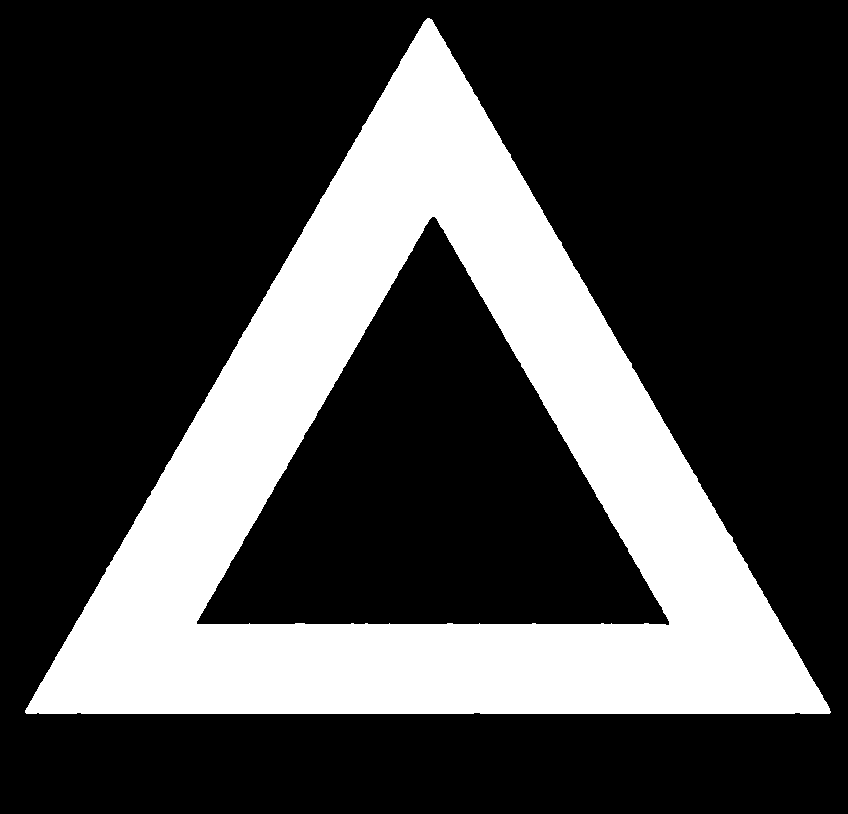
