## Supplementary material for "A machine learning based approach to the segmentation of micro CT data in archaeological and evolutionary sciences": S2. Weka_segmentation_simple_guide

A simple guide to using the trainable Weka segmentation in FIJI.

Open imageJ/FIJI

1. Import your sequence of images (for a single image, can just click open or drag and drop)

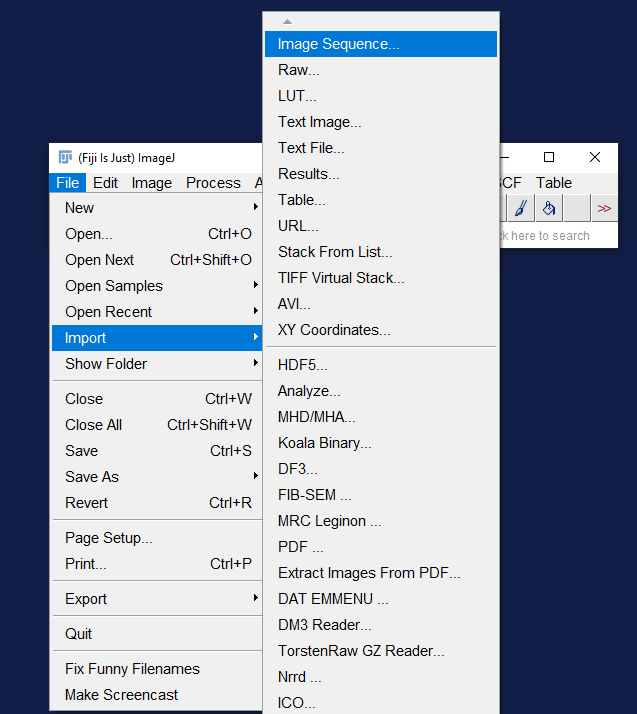

2. Click on one image in the folder

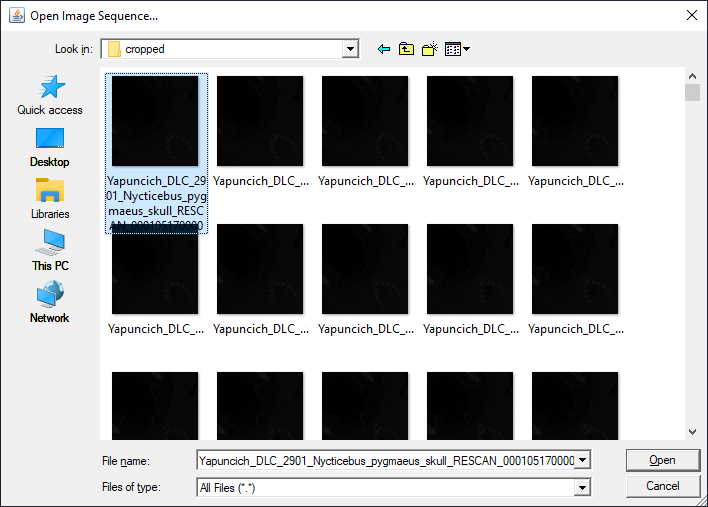

3. This will bring up a dialogue saying what is in the folder. If you only have the image stack in your folder, simply press ‘OK’. For speed of loading with large stacks, it is useful to tick the option ‘Use virtual stack’.

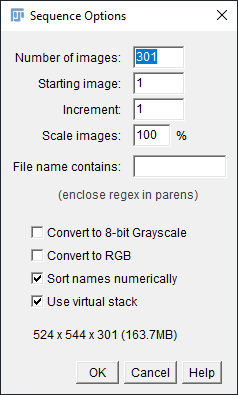

4. Adjust the Window/Level of your object to reflect the greyscale range of your data

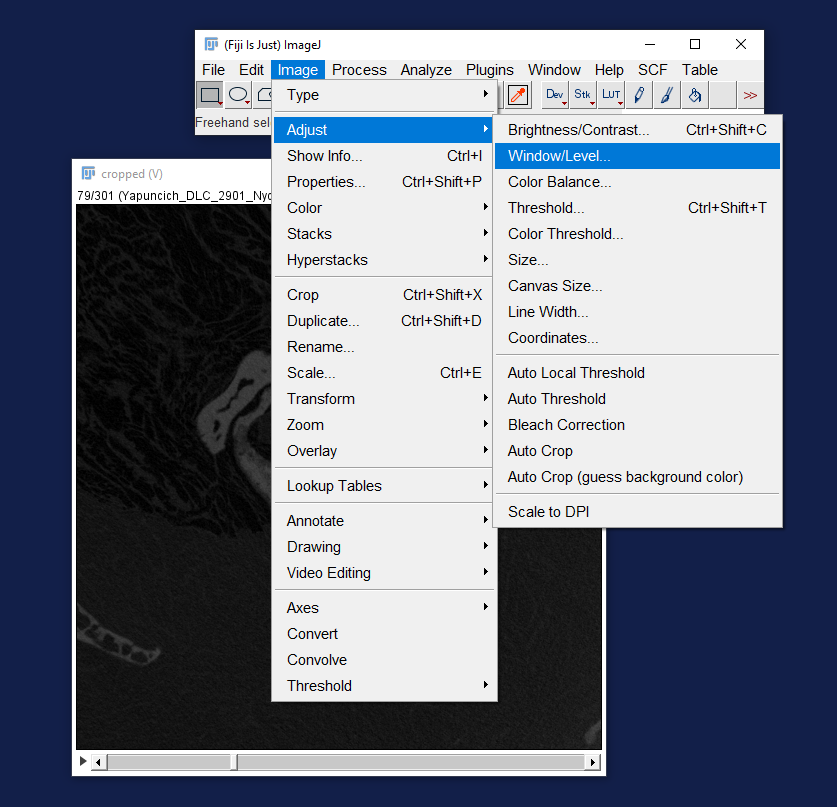

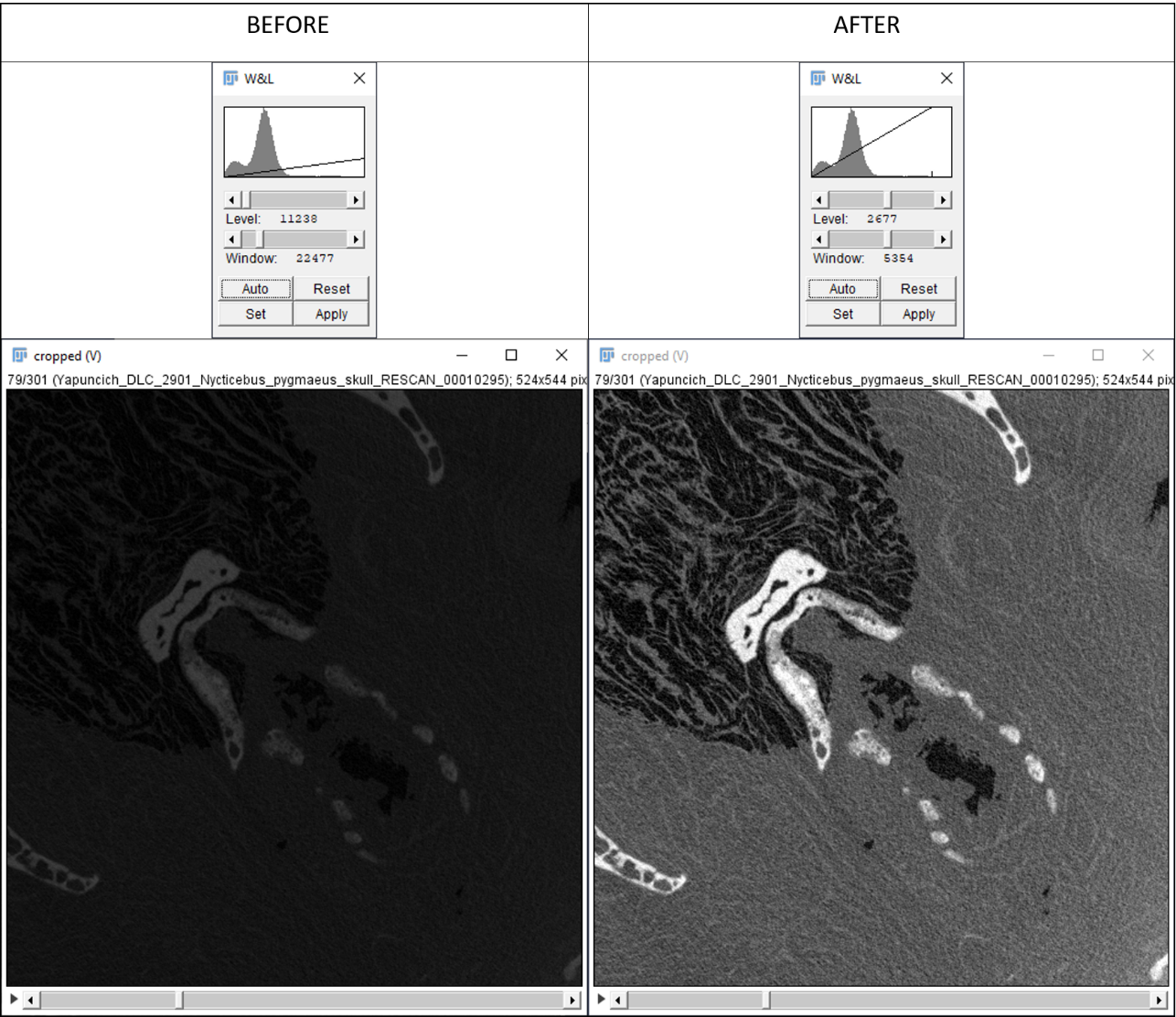

5. Save your stack now using File-Save as image sequence. Unless you put this in a different folder, it will over-write your original data with this new improved contrast stack.
6. Start the Weka Segmentation trainer (Plugins-Segmentation-Trainable Weka Segmentation). If you have a stack open, duplicate an image with lots of different data types in-then you can train on one image only (you can train over the whole stack, but this takes a long time and for many use cases, is not necessary).

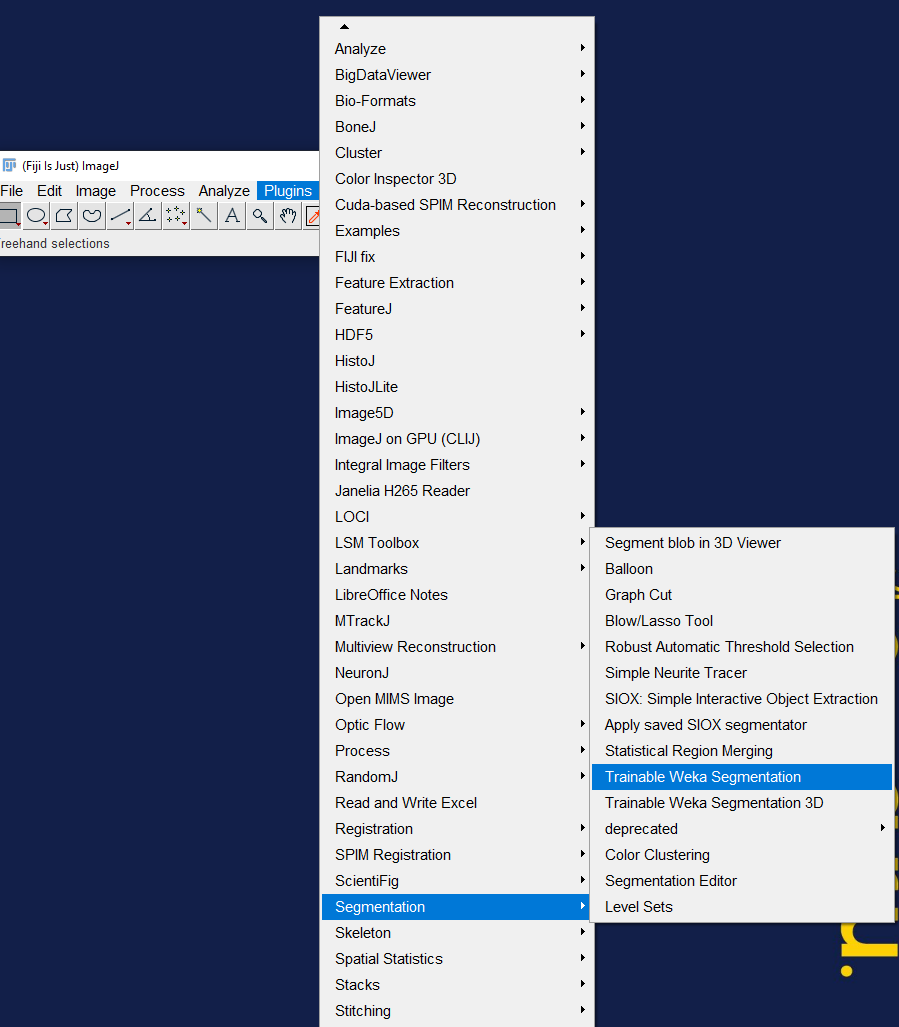

7. Adjust the number of material labels (referred to as class) in Weka. In this case we will have 3 classes. A dialogue box will pop up asking you to give the class a name.

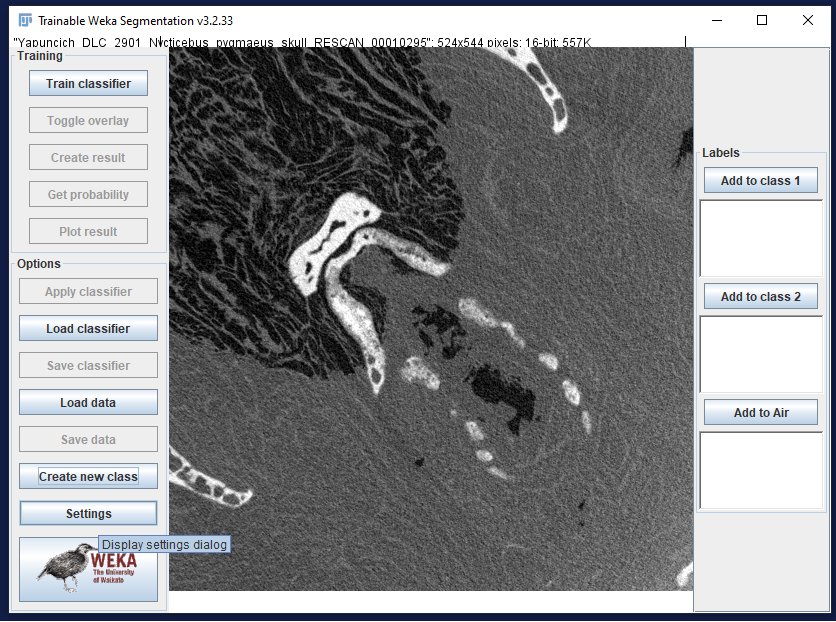

8. You can change the name of the class in ’Settings’. Here, you can also adjust different parameters such as the filters you apply to the data.

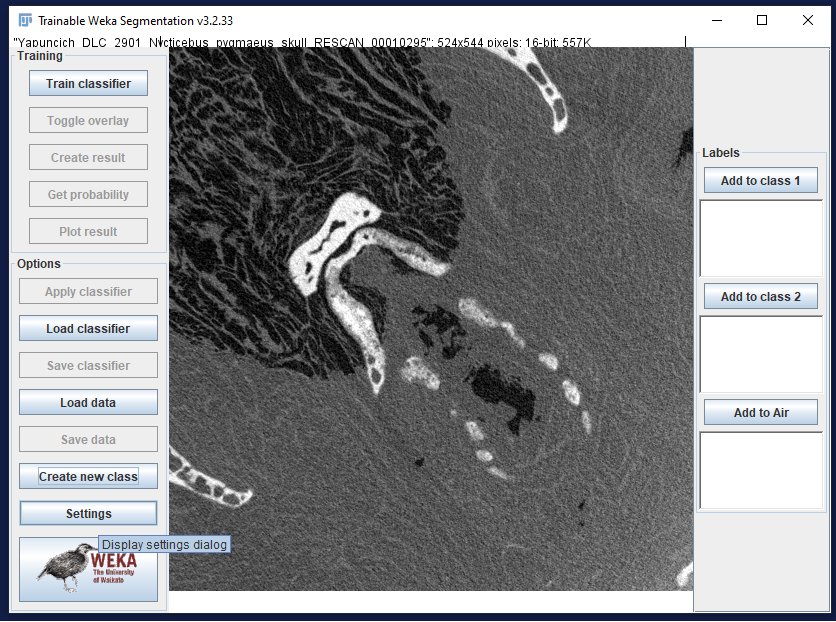

9. Here are the suggested settings for processing CT data, and you can see that we have changed the label names here as well.

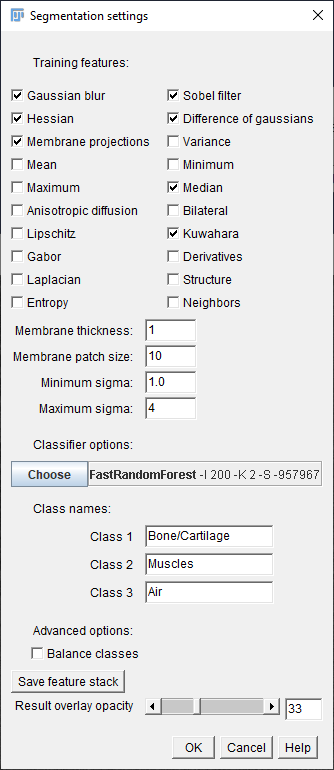

10. Start outlining the material of interest (a graphics tablet is advised for speed and precision). Your outline colour is yellow until you add it to a label, after which it is given a default label colour (in this case, bone is red and you can see that muscular tissue has been highlighted but not added to the label). To add to the label, click the appropriate button.

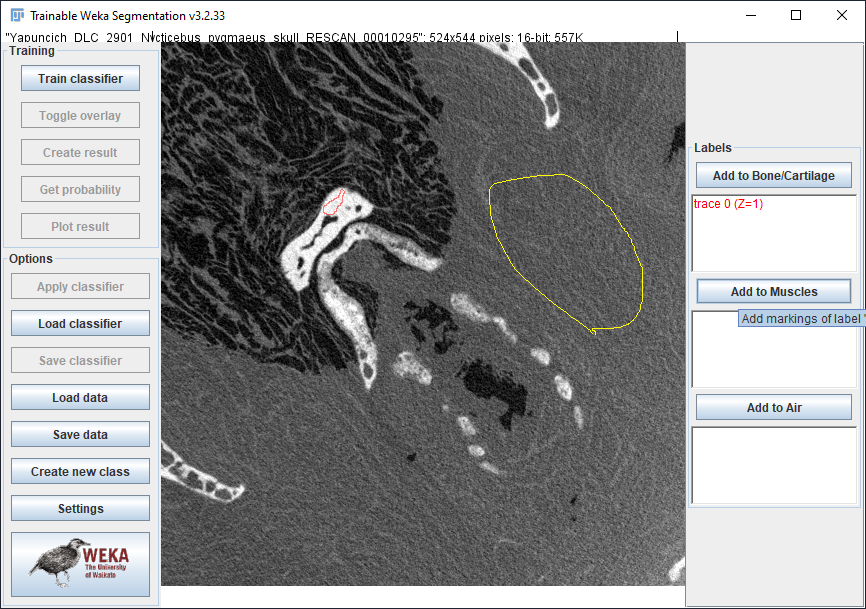

If you realise that you have added the highlighted area to the wrong label, simply click on the trace in the right hand part (highlighte in yellow above). This will highlight it. Double click to remove, or if you want to change label, click the correct label button, then double click to remove the mistaken label (they do not delete automatically after assigning a new label).

11. When you have enough areas assigned manually to labels (generally ony about <10% of the image needs to be marked up) click ‘Train classifier’. Once the training has completed, an overlay will appear on the image, which you can toggle on and off.

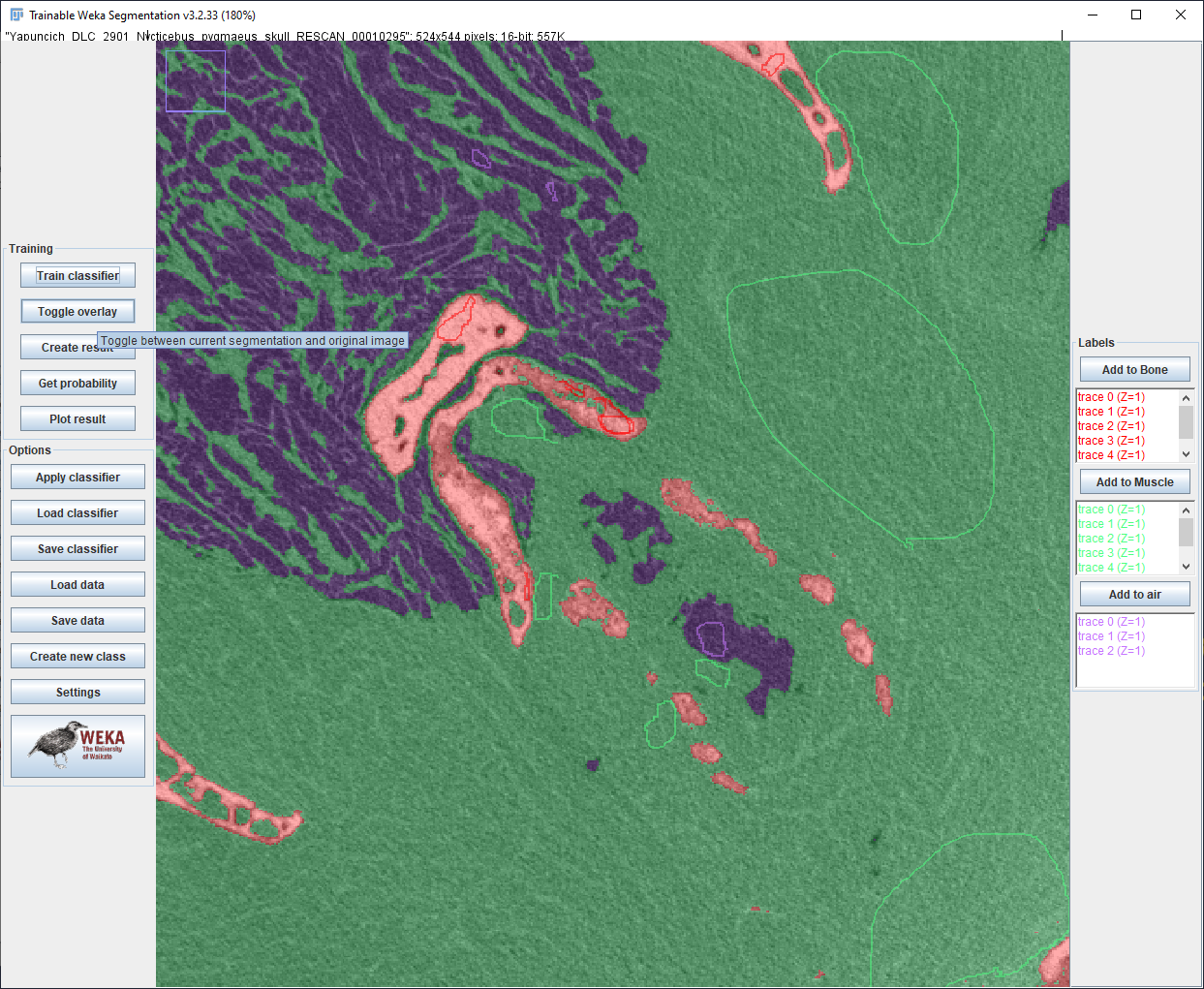

12. The training can be repeated as many times as you like as you add more manually traced data and until you are satisfied with the results. Once you are happy with the classification, it is important to save your classifier. Click ‘Save classifier’ and name the data as you wish.
13. To apply the classifier over a whole sequence of images, click ‘Apply classifier’. This will ask you to locate the images of interest and will classify them accordingly. If you are classifying more than 3 images, it will ask where to save them-it is advisable to create a new explicitly labelled folder for this.

### Scripting classification of images

As noted on the Weka segmentation website, it is often more efficient to script the classification of images when you have a whole folder to classify. Our experience has also found that multi-threading of the random forest algorithm works better when this is done.

To script this process:

1. Open a new Macro by clicking Plugins-Macro-New Macro
2. Copy the beanshell script from <https://imagej.net/Scripting_the_Trainable_Weka_Segmentation#Example:_apply_classifier_to_all_images_in_folder> and paste into the macro editor.
3. Specify the language as Beanshell in the Language tab.
4. Save the script as ‘Filename’, bsh (the extension is very important).
5. I you haven’t restarted ImageJ, just click Run. (If you have, you can drag and drop the file or open it via Plugins-Macros-Run…)

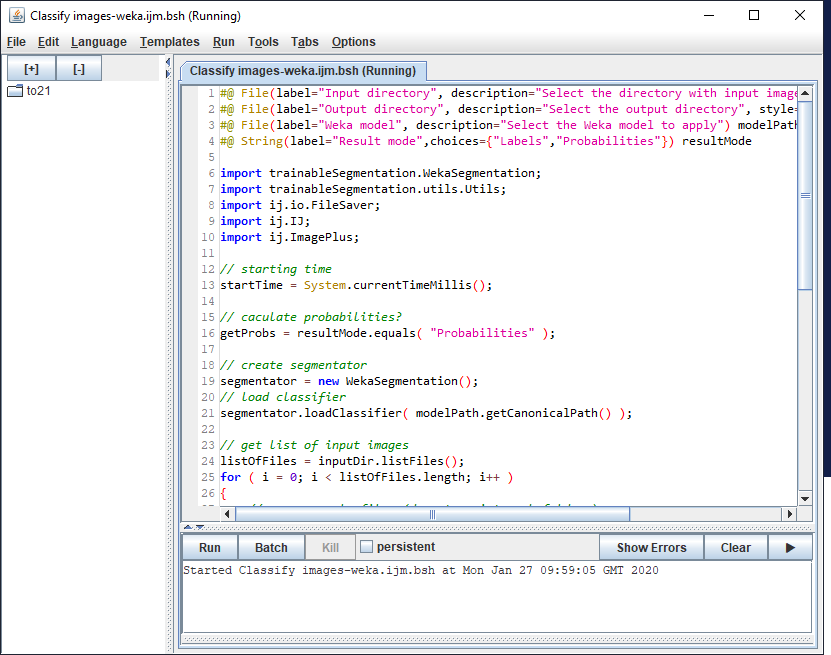

6. A dialogue box will pop up asking you to select the Input and output directorie, and the location of the model file. By default the result mode is ‘label’.

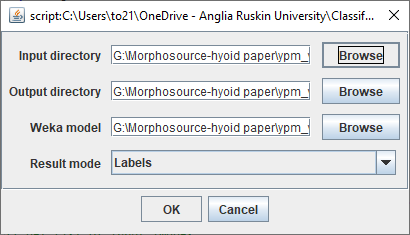
